# Spiny but photogenic: amateur sightings complement herbarium specimens to reveal the bioregions of cacti

**DOI:** 10.1101/2023.03.15.532806

**Authors:** Alice Calvente, Ana Paula Alves da Silva, Daniel Edler, Fernanda Antunes Carvalho, Mariana Ramos Fantinati, Alexander Zizka, Alexandre Antonelli

## Abstract

**Premise:** Cacti are characteristic elements of the Neotropical flora and of major interest for biogeographic, evolutionary, and ecological studies. Here we test global biogeographic boundaries for Neotropical Cactaceae using specimen-based occurrences coupled with data from visual observations, including citizen science records, as a means to tackle the known collection biases in the family.

**Methods:** Species richness and record density were assessed separately for preserved specimens and human observations and a bioregional scheme tailored to Cactaceae was produced using the interactive web application Infomap Bioregions based on data from 261,272 point records cleaned through automated and manual steps.

**Key Results:** We find that areas in Mexico and southwestern USA, Eastern Brazil and along the Andean region have the greatest density of records and the highest species richness. Human observations complement information from preserved specimens substantially, especially along the Andes. We propose 24 cacti bioregions, among which the most species-rich are, in decreasing order: northern Mexico/southwestern USA, central Mexico, southern central Mexico, Central America, Mexican Pacific coast, central and southern Andes, northwestern Mexico/extreme southwestern USA, southwestern Bolivia, northeastern Brazil, Mexico/Baja California.

**Conclusions:** The bioregionalization proposed shows novel or modified biogeographical boundaries specific to cacti, and can thereby aid further evolutionary, biogeographic, and ecological studies by providing a validated framework for further analyses. This classification builds upon, and is distinctive from, other expert-derived regionalization schemes for other taxa. Our results showcase how observation data, including citizen-science records, can complement traditional specimen-based data for biogeographic research, particularly for taxa with specific specimen collection and preservation challenges and those that are threatened or internationally protected.

## INTRODUCTION

Biogeographic regions are fundamental to the study of biogeography because they can inform on dynamic processes of origin, migration, and extinction of evolutionary lineages through time and space (Antonelli, 2017a; Ferrari, 2017; Morrone, 2018). Bioregions and related terms, including areas of endemism, biogeographic realms, phyto- or zoogeographic zones, biomes, ecoregions, or even ecosystems, have been used as operational units of regionalization (Morrone, 2018). Many of these units closely reflect continental divisions, as plate tectonics repeatedly isolated and connected continental biotas through evolutionary time (Antonelli, 2017b; Ficetola et al., 2017). Nevertheless, within continents, smaller patches of more cohesive biotas are also recognizable, but may be more difficult to define, depending on criteria (different methods used to delimit operational units) and context (focus on whole biotas, particular taxonomic groups, or even based on purely abiotic and geographical aspects).

Biotic and abiotic barriers and influences do not affect all organisms equally. Consider for instance that the permeability of a barrier can be different for species bearing animal- or wind-dispersed seeds (e.g, Antonelli, 2009; Nazareno et al., 2021) or even for species dispersed by non-flying small mammals or migrating bats (which can facilitate the connection of isolated populations; e.g., Shilton et al., 1999). Hence, while universal bioregions shared by many taxa delimited by large geographical barriers are important, to understand the significance of geological, climatic and other earth-history processes on the evolution of life (Parenti and Ebach, 2009), taxon-specific bioregionalization schemes are often more valuable for more specific applications, such as ancestral area reconstruction in historical biogeography (Edler et al., 2017). In this case, it is crucial to define the operational bioregions prior to analysis, because these bioregions influence the inference of ancestral areas and consequently the scenarios of origin and dispersal/vicariance events through time.

Metrics of dissimilarity and endemism, as well as species richness, abundance, and rarity, based on occurrence data for species (including the longitude, latitude, date, and other elements of meta-data) are the primary source of data used to delimit bioregions (Harold and Mooi, 1994; Olson and Dinerstein, 1998; Olson et al., 2001). In recent decades, bioregionalization schemes have risen in prominence as a means to analytically support evolutionary, ecological, biogeographic and conservation studies (Olson et al., 2001; Morrone, 2018; Montalvo-Mancheno et al., 2020), and have been propelled by the increase of digitization of biological collections, initiatives to promote public online occurrence databases, and the organization and publication of taxonomic research and inventories in electronic format (Soltis, 2017; Heberling et al., 2021). Recent approaches to bioregionalization also incorporate macroecological principles, ordination, and network methods (Kreft and Jetz, 2010; Vilhena and Antonelli, 2015; Edler et al., 2017; Droissart et al., 2018; Colli-Silva et al., 2019), although many biogeographic studies still use bioregions determined from ‘expert-based’ drawings on maps, a practice that is feasible for many taxa, but on the other hand it is largely subjective, not reproducible, and prevents estimates of uncertainty (Edler et al., 2017; Ferrari, 2017).

Several bioregionalization schemes have been proposed for the Neotropical region (e.g., Olson et al., 2001; De Nova et al., 2012; Hughes et al., 2013; Morrone, 2014; Zizka et al., 2018; Morrone et al., 2022). As probably the most species-rich region on Earth (Kier et al., 2005; Antonelli and Sanmartín, 2011; Zizka, 2019) with an exuberant faunal, fungal, floristic and geo-climatic diversity, the Neotropics comprise a heterogeneous landscape with a great variety of habitats, including interwoven areas of tropical rainforests, seasonally dry tropical forests, savannas, rocky fields, deserts, prairies, swamps, and high altitude and coastal ecosystems. Furthermore, the Neotropics are home to a diversity of life history strategies in plants (such as epiphytes, trees, and fire-adapted geophytes) that a single, general bioregionalization may fail to capture (Antonelli and Sanmartín, 2011; Hughes et al., 2013).

### Cacti as a study system for Neotropical biogeography

Neotropical areas under arid and semi-arid climates form broad, more or less well-defined units in the schemes already proposed for Neotropical regions and are noteworthy for their largely unique biota under high levels of threat (Dinerstein, 2017; Dryflor, 2021; Morrone et al., 2022). Cacti are among the most characteristic elements of the flora of seasonally dry Neotropical areas and are one of the groups of angiosperms best adapted to tropical aridity (Anderson, 2001). Cacti also occur in other Neotropical habitats, such as tropical and subtropical forests, usually occupying water-stressed niches, as epiphytes or lithophytes (Anderson, 2001; Barthlott et al., 2015). The cactus family is almost endemic to the Neotropical region; of around 1,850 known species in 130 genera, only one naturally occurs outside the Americas –the epiphytic and bird-dispersed *Rhipsalis baccifera* (J.S.Muell.) Stearn, which also reaches Africa, Asia, and Sri Lanka (Nyffeler, 2002; Hunt et al., 2006; Nyffeler and Eggli, 2010). The wide distribution, elevated richness and high levels of regional endemism make Cactaceae a suitable model for investigating the diversity, distribution and evolutionary history associated with dry habitats (Silva et al., 2018). Studies focusing on such groups with particular life histories are still poorly documented and may serve as reference to other groups of organisms with congruent distributions in similar conditions (Miranda et al., 2018; Colli-Silva and Pirani, 2020).

The geographic distribution of Cactaceae has been addressed generally, at the family level, based on taxonomic knowledge and checklists and using different analytical tools (Barthlott and Hunt, 1993; Anderson, 2001; Barthlott et al., 2015; Amaral et al., 2022). Traditionally, three centers of diversity are accepted: (1) Mexico and southwestern USA, which comprises the largest number of species; (2) the central Andean region of Peru, Bolivia, northern Argentina, and Chile; and (3) Eastern Brazil (Barthlott and Hunt, 1993; Anderson, 2001). These three centers were also identified by Taylor (1997), who reported a fourth center along central-western and southern Brazil, Paraguay, Uruguay, and central Argentina. Barthlott et al. (2015) estimated distribution ranges for individual species using data from the literature and other various sources and expanded the knowledge on diversity centers, recognizing seven additional subordinated centers: Chihuahua, Puebla-Oaxaca, Sonora-Sinaloan, Jalisco (all four north/central American); Southern central Andes, Caatinga and Mata Atlântica (all three south American).

Other approaches have associated distribution data with conservation assessments, as Cactaceae have historically been affected by illegal trade and habitat loss. For example, Goettsch et al. (2015, 2018) provided a formal global assessment of the conservation status of all cactus species based on point occurrences and expert-reviewed range maps. The study found nearly a third of the species are threatened with extinction (Goettsch et al., 2015, 2018). Amaral et al. (2022) used occurrence and phylogenetic data to investigate spatial patterns and factors associated with endemism and concluded that legally protected areas do not guarantee the evolutionary conservation of the family. Furthermore, these authors suggested that different abiotic factors may contribute to the prediction of endemism in the group. Pillet et al. (2022) used species distribution models to predict a negative impact of future climatic changes, with severe extinction risk for most species they assessed. Although these earlier studies generated a comprehensive general understanding of cacti distribution, historical biogeography studies on the group have relied on general schemes of Neotropical bioregionalizations (e.g., Ocampo and Columbus, 2010; Calvente et al., 2011; Vazquez-Sanchez et al., 2013; Hernandez-Hernandez et al. 2014; Lavor et al., 2018; Majure et al., 2022) and the validity of such general schemes for cacti remains to be tested.

Most studies of spatial biogeography in plants, including cacti, use preserved herbarium specimens point records as primary evidence (Funk and Richardson, 2002; Folk and Siniscalchi, 2021). However, this is problematic for cacti due to a known collection deficit associated with the difficult handling and preservation of specimens. Cacti are often ignored by collectors, because they are succulent, spiny and grow in extreme habitats, and the techniques to adequately preserve cactus specimens require special training (Anderson, 2001; Taylor and Zappi, 2004). The preservation of specimens of rare and endangered cacti can also be additionally challenging and biased, since many collectors avoid collecting these taxa, particularly when the whole plant must be collected for adequate preservation. It is also important to note that all cacti (except species in genera *Pereskia, Pereskiopsis* and *Quiabentia*) are legally protected by the Convention on International Trade in Endangered Species of Wild Fauna and Flora (CITES, appendices I and II) since 1975.

Despite those challenges, Cactaceae systematics has a wide appeal and even reaches the general public, since cacti are charismatic, well known due to their popularity as house plants, and easy to identify at family level. Consequently, synthesizing information about the distribution of cacti can benefit from networks of amateur enthusiasts and horticultural societies, and other sources of information such as personal communications, field observations and pictures. When combined with academic knowledge, these additional sources of information can provide an integrated understanding about the general occurrence of species (e.g., Taylor and Zappi, 2004; Goettsch et al., 2015, 2018). Despite legitimate concerns about the limits of data based solely on human observations, citizen-science data nevertheless hold the potential to complement the information provided exclusively by traditionally preserved specimens, particularly where the sources are well-referenced images linked to public databases that benefit from being curated to some degree (Troudet et al., 2018).

Here, we combine data from preserved specimens and human observations gathered from public databases to analyse diversity patterns for Cactaceae and then produce a bioregionalization scheme for the family based on network analysis, also integrating phylogenetic information. We aim to test the application of data from visual observations of cacti (including citizen-science-based data) in spatial biogeographical analyses and to test global biogeographic boundaries for the family. In particular, we address the following questions:

1. Are occurrence data currently available through public databases, including human observation records, consistent with scientific knowledge and therefore capable of providing a comprehensive distribution dataset for Cactaceae?
2. What are the bioregional boundaries for Cactaceae obtained from a large dataset?
3. Is a bioregional scheme tailored for Cactaceae compatible with general Neotropical bioregionalization schemes?

## MATERIALS AND METHODS

We downloaded a total of 261,272 georeferenced records from preserved specimens (28%) and human observations (72%) for Neotropical Cactaceae (Table 1) using rgbif v.1.3.0 (Chamberlain et al., 2019) implemented in R (R Core Team, 2020). To delimit the Neotropics, we used the polygon defined as: −34.7 32.8, −117.2 32.8, −117.2 −55.8, −34.7 − 55.8, −34.7 32.8 (Fig. 1). We included human observations to explore their potential to complement preserved specimen data due to the known collection bias in Cactaceae (Anderson, 2001; Taylor and Zappi, 2004). To access the contribution of each type of record on the resulting distribution and bioregionalization and to allow a clearer comparison of record type, we performed analyses separately for: (1) preserved specimens; (2) human observations (excluding *i*Naturalist, as these include mostly science-based and government datasets, but also includes less representative citizen-based datasets; Table S1); and (3) *i*Naturalist (the major source of citizen-based occurrences in our dataset, which included nearly 90% of all human observations; Table 1).

**Table 1.** Results for raw and cleaned datasets obtained from each record source.

|  | Preserved specimens |  | Human observations<br>(excl. *iNaturalist*) |  | *iNaturalist* |  | All sources |  |
| --- | --- | --- | --- | --- | --- | --- | --- | --- |
|  | Raw | Cleaned | Raw | Cleaned | Raw | Cleaned | Raw | Cleaned |
| No. of records | 71,957 | 29,898 | 21,078 | 9,599 | 168,237 | 98,163 | 261,272 | 137,660 |
| No. of species | 1,419 | 1,106 | 418 | 362 | 1,144 | 902 | 1,573 | 1,248 |

To minimize errors due to coordinate imprecision, data cleaning was performed through CoordinateCleaner v. 2.0-9 (Zizka et al., 2019) in R. There is no universal solution to clean and process species occurrence data (Zizka et al., 2020a). To visualize and help us evaluate the performance of steps of automated data cleaning for cacti, we produced richness and occurrence density maps of raw and cleaned datasets using the tidyverse v.1.3.2 (Wickham et al., 2019), speciesgeocodeR v.2.0-10 (Töpel et al., 2017), raster v.3.5-29 (Hijmans, 2018) and rgdal v.1.5-32 (Bivand et al., 2015) packages. Automated cleaning excluded records: (1) without geographical coordinates; (2) in a 10 km radius from country centroids; (3) in the headquarters of biodiversity institutions, such as museums, botanical gardens and universities; (4) in open sea; (5) with coordinates equal to zero; (6) with equal latitudes and longitudes; (7) collected prior to 1945 (except for preserved specimens from Argentina, for which we included all records, because they lacked data for collection dates); (8) with invalid or imprecise coordinates; (9) not identified at least to species level; and (10) duplicated (same coordinates) for the same species (Zizka et al., 2020b). We further manually standardized names following the classification adopted in Korotkova et al. (2021). To minimize the registered occurrence of cultivated and naturalized species (both in preserved specimens and human observation data) we checked all individual species distribution maps against the geographic distribution cited in the most recent comprehensive works produced for the family (Hunt et al., 2006; Barthlott et al., 2015, Hunt, 2016) and excluded non-matching records in QGis 3.6.1 (Qgis, 2021). In this final manual cleaning we excluded all records of the widely cultivated species *Opuntia cochenillifera* (L.) Mill., *O. ficus-indica* (L.) Mill., *Cereus hildmannianus* K.Schum., *Pereskia grandifolia* Haw., *Schlumbergera truncata* (Haw.) Moran and *Selenicereus undatus* (Haw.) D.R.Hunt and cleaned individual records of another 109 species (Table S2).

We inferred bioregions using the cleaned datasets of preserved specimens and human observations separately, and a combined dataset comprising both data sources. We used the interactive web application Infomap Bioregions v.1.2.0 (Edler et al., 2017) for bioregionalization, which is an easy-to-use method based on distribution data, bipartite networks, and network clustering to detect single non-hierarchical solution schemes (Vilhena and Antonelli, 2015; Kheirkhahzadeh et al., 2016). Infomap Bioregions first bins the world into grid cells with adaptive resolution based on the density of the data: starting from a maximum cell size, if there is at least a selected minimum number of records in a cell, it is included in the analysis. Conversely, if there is more than the pre-selected maximum number of records, the cell is recursively subdivided into four grid cells until the maximum capacity is respected or minimum cell size is reached. The software then creates a network between species and grid cells by connecting each species to all grid cells where it is found. It uses the Infomap network clustering algorithm to find an optimal partition of the network into groups of nodes more tightly inter-connected within than between the groups. The set of grid cells within each group makes up a bioregion.

Infomap has a resolution parameter called Markov time (called “cluster cost” in Infomap Bioregions v1) that can be used to zoom in and out for solutions on different scales (Kheirkhahzadeh et al., 2016). Further analyses were carried out in Infomap Bioregions v.2.6.1 using newly implemented tools to explore hierarchical solutions (Rosvall et al., 2011), detect interzones (transition zones) or fuzzy borders between bioregions where their taxa mix (Bloomfield et al., 2018) and using variable Markov time to adapt Infomap’s resolution to the network density (Edler et al., 2022). With a constant Markov time, increasing it to avoid fragmentation of sparse regions tends to collapse dense regions. Variable Markov time increases the range of scales of bioregions that Infomap explores by locally increasing Markov time on sparse regions.

Grid cell sizes of 1° x 1° and 2° x 2° and with an adaptive resolution of 1/8° to 4° were tested to choose the best fit for the data in the Neotropical area; for final figures we used the 1/2° to 2° adaptive grid cells and a maximum of 200 and a minimum of 3 records per cell, which showed the best fit as using the resolution of cells smaller than 1/2° and a minimum of less than 3 records per cell caused the fragmentation in sparse bioregions in areas with too few records. Using cells larger than 2° caused the loss of definition of bioregional borders in several areas, such as in Mexico and in the Andean region. We used a maximum of 200 records per cell as this was close to the number of records in the cell with the highest number of records. Due to unequal sampling efforts for Cactaceae along the Neotropics, we tested, but chose not to use, the weight on abundance option for the final results. Different cluster costs in 10 trial runs were tested (ranging from 0.5–2.0) to allow the search for larger or smaller bioregions (Kheirkhahzadeh et al., 2016; Edler et al., 2017) and the 0.92 cluster cost was selected to fit a wider continental scale inference. All these subdivisions were supported by the data and merely differ in their value for downstream applications: a smaller number of bioregions may be most useful for continental-scale biogeographic analyses, whereas a larger number of fragmented units may hold higher value for conservation purposes, which could be further explored and ground-truthed. Here we explore further a larger continental scale scheme highlighting bioregions larger than one 2° x 2° grid cell. Species richness and occurrence density maps were also produced in adaptive 1/2° to 2° grid cells in Infomap Bioregions (Edler et al., 2017) and were further edited in QGis 3.6.1 (Qgis, 2021).

Hierarchical solutions were also explored using Markov times varying from 0.7 to 1.2 and under the variable Markov time option. Optimal results with smaller bioregions in lower levels for South America and North America were found under different Markov times (1.18 and 1, respectively). Those were nearly identical to the single solution scheme in lower levels and are shown in the Supplementary Material (Figs. S6 and S7). An alternative hierarchical solution capturing larger bioregions that consistently emerged in schemes obtained under varying parameters is shown for North and South America simultaneously using the variable Markov time option under Markov time=1.

Bioregionalization solutions were also explored using a newly developed method implemented in Infomap Bioregions 2 that incorporates evolutionary relationships into species occurrence networks leading to the definition of more historically meaningful bioregions (Edler et al., 2023). It connects nodes from the phylogenetic tree to the grid cells where their descendant species occur, weighted by the amount of geographic information they provide. This makes the network more dense within areas with closely related species and tends to dissolve bioregional boundaries crossing such areas. Similar to ancestral nodes, individual wide-ranging species can also obscure and collapse modular patterns and transition zones defined by range-restricted species if the links between species and grid cells are unweighted. As narrowly distributed species are important for unveiling biogeographic patterns and evolutionary processes (Laffan et al., 2016; Quintero and Jetz 2018), we use range-weighted species by default in Infomap Bioregions 2, treating species nodes similar to ancestral nodes.

To perform this analysis, we built a phylogenetic backbone for Cactaceae using a top-down approach departing from a large phylogenetic tree for angiosperms, standardized according to the botanical nomenclature of The Plant List (GBOTB.extended.TPL.tre) implemented in the U.PhyloMaker package (Jin and Qian, 2023) and edited to include all species sampled in our occurrence records dataset. We initially built a tree pruned from the megatree including species that matched our dataset (616 spp.; Fig. S9). This tree captures relationships between major clades and genera of Cactaceae consistent with the literature (e.g. Nyffeler and Eggli, 2010; Guerrero et al., 2018). To avoid making assumptions about specific placements for species not yet sequenced, while making use of morphologically informed taxonomic classifications, we then added the remainder of the species binding them as polytomies to the first diverging node of their respective genus (when the genus was present in the megatree) or major clades in Cactaceae (when the genus was absent from the tree). We used the Open Tree of Life (Open Tree of Life, www.tree.opentreeoflife.org, searched in June 2023) as a reference backbone to bind genera to large clades (tree 2, available in the supplementary material). As an alternative, we built a second tree with species from genera absent in the megatree binded to the first diverging node of Cactaceae (tree 1, available in the supplementary material). Both trees were tested in the bioregionalization analyses and produced similar results. The final trees match the nomenclature of Korotkova et al. (2021) as our occurrence dataset was standardized according to it. For this analysis in Infomap Bioregions 2 we used the previously described settings and the options to integrate the whole tree, 100% tree weight, markov time=1 and to weight species by range.

## RESULTS

### Data source

The automated filtering and manual cleaning resulted in the exclusion of 58% of records and 22% of species names for preserved specimens, 54% records and 13% of species names for human observations (excluding *i*Naturalist), and 42% of records and 21% of species names for *i*Naturalist data (Table 1). The cleaned complete dataset with data from all records combined resulted in 137,660 records for 1,248 species (Table S2), which corresponds to 67% of all accepted species of Cactaceae listed in Korotkova et al. (2021). Data from preserved specimens included records for 60% of species, *i*Naturalist included records for 49% of species, and human observations (excluding *i*Naturalist) included records for 20% of species of Cactaceae (Table 1).

Each data category showed similar patterns of record density per species, with most species with few records and few species with numerous records (Fig. S3). Nevertheless, for human observations (including *i*Naturalist), a single species—the saguaro, *Carnegiea gigantea* (Engelm.) Britton & Rose—accounted for 16% of records (17,384 records), which is four times more records than the second most recorded species (*Cylindropuntia leptocaulis (DC.)* F.M.Knuth with 3,830 records). For preserved specimens, in contrast, the most recorded species—*Rhipsalis baccifera* (J.S.Muell) Stearn—had only a few more records than the second most recorded species (*Opuntia engelmannii* Salm-Dyck ex Engelm. with 853 records), accounting for only 3% of the records (891 records). The vast majority of records from *i*Naturalist were from Mexico and the USA, where 90,772 observations (93%) were recorded for 476 species, and only 7% were recorded in all other Neotropical countries together (7,391 observations for 442 species).

We found occurrences for species of cacti throughout the Neotropics, although in some patches of the core Amazonian region few species were documented (Fig. 1). Areas in Mexico and southwestern USA, Eastern Brazil and along the Andean region had the greatest density of records considering both preserved specimens and human observations (Fig. 1a-d, Table 2). Eastern Brazil was better sampled through preserved specimen data, and *i*Naturalist showed a greater density of records in Mexico, southeastern USA and Andean regions (Fig. 1, Table 2).

**Table 2.** Number of records and species in the three main centers of diversity for Neotropical Cactaceae.

|  | Preserved specimens |  | Human observations<br>(excl. *iNaturalist*) |  | *iNaturalist* |  | All sources |  |
| --- | --- | --- | --- | --- | --- | --- | --- | --- |
|  | Records | Species | Records | Species | Records | Species | Records | Species |
| Andean region | 2,426 | 298 | 800 | 36 | 4,497 | 292 | 8,367 | 376 |
| Brazil | 5,254 | 174 | 242 | 14 | 378 | 67 | 5,920 | 175 |
| Mexico/USA | 17,249 | 553 | 7,714 | 275 | 86,862 | 476 | 111,171 | 589 |

Species richness maps recovered three main diversity centers for Cactaceae (Fig. 1e-h, Table 2): Mexico/SW USA (589 spp., 81% of records), the Andean region (376 spp., 6% of records) and Eastern Brazil (175 spp., 4% of records). Although we recorded more species overall in the Andean region compared with Eastern Brazil, the number of preserved specimens from the former was lower (2,426 records for 376 species in the Andean region versus 5,254 for 175 species in Eastern Brazil). The addition of human observation records (including *i*Naturalist) increased both spatial coverage and density of records and increased the species richness estimated for the Andean region by 21% (Fig.1, Table 2). The addition of human observation records in Central/North America also led to a noteworthy increase of 6% in the estimation of species richness, while in Eastern Brazil this increase was only 0.6% (Table 2).

Most of the species (104 of 108 species) for which we obtained data only from human observations in our dataset occur mainly in the Andean (68%) and Central/North American (29%) centers of diversity and show local or subregional restricted distributions, not exceeding 600km^2^ in geographic range (Table 3). A significant portion (57%) of these species have few herbarium records in the Global Biodiversity Information Facility (GBIF, www.gbif.org, searched in September 2022), have some degree of threat (24%) or are Data Deficient (18%) in the IUCN Red List (IUCN, 2022), and a 6% were involved in some degree of taxonomic uncertainty in the last 15 years.

**Table 3.** Source of records, herbarium sampling, conservation status, distribution and taxonomic status for species recorded only from human observations in our dataset (108 in total) in two main centers of diversity. All species from the Brazilian center of diversity had preserved specimens in our dataset. The degree of sampling (<10 records is considered with few records/ > 10 is considered well recorded) is based on the number of herbarium records documented in GBIF for these species (these records were excluded during automated filtering steps due to low quality or absent georeferenced data). Threat categories included were CR, EN and VU according to the IUCN Red List (IUCN, 2022). Taxa included as taxonomic synonyms of other taxa in the last 15 years were considered as involved in taxonomic uncertainty. Geographic distributions are based on distribution ranges presented for species in Barthlott et al. (2015); species with ranges <200km^2^ are considered locally restricted, species with ranges >200km^2^ and <600km^2^ are considered sub regionally restricted.

|  | Mexico/ SW USA | Andean region |
| --- | --- | --- |
| No. Species | 31 (29%) | 73 (68%) |
| Source of records | 93% only from *iNaturalist*, 7% from *iNaturalist* and other sources of human observations | 95% only from *iNaturalist*, 4% from *iNaturalist* and other sources of human observations, 1% only from other sources of human observations |
| Herbarium sampling | 68% few records, 32% well recorded | 52% few records, 48% well recorded |
| Conservation status | 45% LC, 39% threatened, 16% DD | 63% LC or NT, 19% DD or not accessed, 18% threatened |
| Distribution | 81% local, 19% subregional | 70% local, 26% subregional, 4 % not included |
| Taxonomic status | 100% well-defined | 90% well-defined, 10% involved in taxonomic uncertainty |

### Bioregions

Based on combined human observation and preserved specimen data, we inferred 24 bioregions for Neotropical Cactaceae using the non-hierarchical solution method implemented in Infomap Bioregions v.1.2.0 (Fig. 2; Table 4; shape files available in the supplementary material). A scheme based exclusively on preserved specimens (Fig. S4) produced very similar results, with all major bioregions coinciding. The scheme based on combined data shows a clearer definition of bioregional borders in areas with lower density and spatial coverage of preserved specimen records (Fig. 1), such as in northwestern and midwestern South America, so we interpreted both results as complementary (Figs. 2, S4).

**Table 4.** Summary data for 24 bioregions of Neotropical Cactaceae: total number of species and number of species exclusive to the bioregion, estimated size (number of 2**°**x2**°** grid cells), geographical location and correspondence with biomes of Dinerstein et al. (2017) and biogeographic provinces and dominions of Morrone et al. (2022). The ten most species-rich bioregions are highlighted in bold. * Indicate bioregions including areas with higher phylogenetic diversity (0.06-0.241, according to Amaral et al., 2022)

| Bioregion | N° of species/<br>exclusives | N° of<br>grids | Geographic location | Biomes | Biogeographic provinces<br>(dominions/transition<br>zones) |
| --- | --- | --- | --- | --- | --- |
| **Bioregion**<br>**01\*** | 134/67 | 40 | Continuous in mid to southern Andean region, mostly in Argentina and Chile | DXS, MFWS, MGS, TBMF, TGSS, TSGSS, TSMBF | Chaco and Pampean (Chacoan); Puna, Atacama, Comechingones, Cuyan High Andean, and Monte (South American); small portions of Yungas (South Brazilian) |
| Bioregion 02 | 59/49 | 11 | Continuous in Chile and along the Argentinian border | DXS, MFWS, MGS, TBMF | Puna, Atacama, and Cuyan High Andean (South American) |
| **Bioregion 03\*** | 132/62 | 9 | Continuous mostly in southwestern Bolivia | DXS, MGS, TSDBF, TSGSS, TSMBF | Yungas and Rondônia (South Brazilian); Chaco (Chacoan); Puna and Monte (South American) |
| Bioregion 04 | 82/42 | 11 | Continuous mostly in southern Peru and northern Chile | DXS, MGS, TSMBF | Rodônia and Yungas (South Brazilian); Desert, Puna, and Atacama (South American) |
| Bioregion 05 | 47/24 | 15 | Continuous in Peru and Ecuador plus a disjunct small area in central Colombia | DSX, FGS, Ma, MGS, TSDBF, TSGSS, TSMBF | Cauca, Magdalena, Western Ecuador and Ecuadorian (Pacific); Napo (Boreal); Sabana (Pacific); Ucayali and Yungas (South Brazilian); Páramo, Desert and Puna (South American) |
| Bioregion 06 | 10/5 | 1 | Restricted to a small portion of Central Peru | TSMBF | Rodônia, Ucayali and Yungas (South Brazilian) |
| Bioregion 07* | 84/15 | 42 | Nearly continuous in eastern Argentina, eastern Bolivia, Paraguay, and smaller patches in Uruguay | FGS, MGS, TGSS, TSDBF, TSGSS, TSMBF | Rodônia, Madeira and Yungas (South Brazilian); Cerrado, Chaco and Pampean (Chacoan); Araucaria, Esteros del Iberá and Parana Forest (Paraná); Monte and Puna (South American) |
| Bioregion 08 | 21/7 | 5 | Disjunct patches in central Brazil (Cerrado region) | TSGSS, TSMBF | Cerrado (Chacoan) |
| Bioregion 09* | 91/31 | 31 | Continuous mostly in coastal northeastern, southeastern, southern Brazil and southern Paraguay | Ma, TSDBF, TSGSS, TSMBF | Caatinga, Cerrado, Chaco (Chacoan); Araucaria Forest, Atlantic, Esteros del Iberá, Parana Forest and Southern Espinhaço (Paraná) |
| Bioregion 10 | 59/16 | 80 | Mostly continuous in northern South America and with disjunct patches spread in South America | DXS, FGS, Ma, MGS, TSCF, TSDBF, TSGSS, TSMBF | All provinces in Mesoamerican, Pacific, Boreal Brazilian and South Brazilian dominions; Xingu-Tapajós (South-eastern Amazonian); Cerrado and Chaco (Chacoan) |
| Bioregion 11 | 44/19 | 7 | Continuous in extreme southern Brazil and Uruguay | TSGSS, TSMBF | Pampean (Chacoan); Esteros del Iberá and Parana Forest (Paraná) |
| Bioregion 12 | 13/2 | 9 | Continuous in central Brazil | FGS, TSDBF, TSGSS, TSMBF | Cerrado (Chacoan); Parana Forest (Paraná); and Rondônia (South Brazilian) |
| **Bioregion 13\*** | 121/75 | 35 | Continuous in northeastern Brazil, including north of Minas Gerais state | Ma, TSDBF, TSGSS, TSMBF | Caatinga and Cerrado (Chacoan); Pará (Boreal Brazilian); Atlantic, Chapada Diamantina, Parana Forest and Southern Espinhaço (Paraná) |
| **Bioregion 14\*** | 156/18 | 17 | Continuous along Mexican pacific coast | DXS, Ma, TSCF, TSDBF | Sierra Madre Occidental, Transmexican Volcanic Belt, and Sierra Madre del Sur (Mexican); Pacific Lowlands and Balsas Balsin (Mesoamerican) |
| **Bioregion 15** | 102/42 | 14 | Continuous in Mexico (Baja California) plus disjunct small patches along the Mexican pacific coast | DXS, Ma, MFWS, TSDBF | Pacific Lowlands (Mesoamerican) |
| **Bioregion 16\*** | 132/23 | 16 | Continuous in northwestern Mexico and extreme southwestern USA | DXS, Ma, MFWS, TCF, TSCF, TSDBF | Sierra Madre Occidental (Mexican); Pacific Lowlands (Mesoamerican) |
| **Bioregion 17\*** | 223/33 | 11 | Continuous in central Mexico | DXS, TSCF, TSDBF, TSMBF | Sierra Madre Occidental, Sierra Madre Oriental, and Transmexican Volcanic Belt (Mexican); Balsas Balsin, Pacific Lowlands and Veracruz (Mesoamerican) |
| **Bioregion 18\*** | 165/35 | 5 | Continuous in southern central Mexico and Oaxaca | DXS, TSCF, TSDBF, TSMBF | Transmexican Volcanic Belt and Sierra Madre del Sur (Mexican); Balsas Basin, Pacific Lowlands and Veracruz (Mesoamerican) |
| **Bioregion 19\*** | 283/99 | 37 | Continuous in northern Mexico, southwestern USA | DXS, Ma, TBMF, TCF, TGSS, TSCF, TSGSS, TSMBF | Sierra Madre Occidental and Sierra Madre Oriental (Mexican); Veracruz (Mesoamerican) |
| Bioregion 20 | 11/2 | 12 | Continuous in Florida (USA) plus patches in Cuba | DXS, FGS, Ma, TBMF, TGSS, TSCF, TSDBF, TSMBF | Cuban (Antillean) |
| Bioregion 21 | 39/28 | 14 | Disjunct in the Caribbean | DXS, FGS, Ma, TSCF, TSDBF, TSMBF | Cuban, Jamaica, Hispaniola, Puerto Rico (Antillean) |
| **Bioregion 22\*** | 161/24 | 30 | Continuous in Central America | DXS, Ma, TSCF, TSDBF, TSGSS, TSMBF | Chiapas Highlands and Sierra Madre del Sur (Mexican); Mosquito, Pacific Lowlands, Veracruz, Yucatán Peninsula (Mesoamerican) |
| Bioregion 23 | 42/13 | 6 | Continuous in the portions of Central America plus a disjunct patch in coastal Venezuela | DXS, TSCF, TSDBF, TSMBF | Guatuso-Talamanca, Puntarenas-Chiriqui and Venezuelan (Pacific); Pacific Lowlands (Mesoamerican) |
| Bioregion 24 | 3/3 | 3 | Continuous in Galapagos (Ecuador) | DXS | Galapagos Islands (Brazilian) |

Eight of these bioregions include more than 100 species and occupy 10 or more grid cells: bioregions 19 (Mexico/USA), 17 (central Mexico), 22 (Central America), 14 (Mexican Pacific coast), 1 (mid to southern Andean region), 16 (NW Mexico/extreme SW USA), 13 (NE Brazil) and 15 (Mexico/Baja California) (Table 4). Bioregions 3 (SW Bolivia) and 18 (southern central Mexico), although smaller bioregions, also included more than 100 species. These ten bioregions are also remarkable for the high number (18-99) of within-bioregion endemics and nine include areas with higher phylogenetic diversity (Table 4; Amaral et al., 2022).

Six of the most species-rich bioregions are in Mexico and Southwestern USA (Table 4). Bioregion 19 is continuous, spreading along northern Mexico and southwestern USA, and it is the most species rich, including 283 species, of which 99 occur exclusively in its area. Exclusive species of globular to subglobular cacti in the genera *Echinocereus, Coryphantha*, *Turbinicarpus*, *Mammillaria*, *Sclerocactus* and *Escobaria* are indicative of this bioregion (Table S5). Bioregion 17 is adjacent to the south of Bioregion 19, spreading continuously in Central Mexico. Exclusive globular to subglobular species in the genera *Stenocactus*, *Coryphantha*, *Echinocereus*, *Mammillaria*, *Thelocactus* and *Turbinicarpus* are indicative of this bioregion. Bioregion 14 is adjacent to the west, spreading through the west coast of Mexico. Various species of columnar, bushy, or globular cacti are indicative of this bioregion, including exclusive species of *Stenocereus*, *Pereskiopsis*, *Mammillaria*, *Pachycereus*, *Echinocereus*, *Selenicereus* and *Acanthocereus*. In the lower levels of an alternative hierarchical solution for North America, the northern section of Bioregion 14 emerges in a separate bioregion, in the intersection of Bioregion 16 (Fig. S7). Bioregion 16 ranges to the north of Bioregion 14 and to the west of Bioregion 19, and is characterised by exclusive globular to spiny, cylindrical species of *Sclerocactus*, *Echinocereus*, *Cochemiea* and *Grusonia*. Bioregion 15 is mostly in Baja California, and is characterised by species of globular to spiny, cylindrical, or flattened species of *Echinocereus*, *Ferocactus*, *Cylindropuntia*, *Cochemiea* and *Opuntia*. Finally, Bioregion 18 is in southern Mexico and Oaxaca, and is characterised by exclusive species of *Mammillaria*, *Stenocereus* and *Opuntia* with various stem forms.

The other four most species-rich bioregions occur further to the south in the Americas. Bioregion 22 is a large bioregion that ranges throughout extreme Southern Mexico and Central America. There, the indicative species are exclusive spiny scandent shrubs or epiphytes of *Selenicereus*, *Epiphyllum*, *Deamia* and *Acanthocereus* and also one columnar *Lemaireocereus*, one globular *Cochemiea* and one flattened *Opuntia*. Bioregions 1 and 3 are in the Andean region of Argentina, Chile, and Bolivia. In Bioregion 1 the indicative species are exclusive spiny cylindrical or globular species of *Rebutia*, *Echinopsis*, *Maihueniopsis*, *Tephrocactus*, *Acanthocalycium* and *Gymnocalycium* and one columnar *Soehrensia*. In Bioregion 3 the indicative species are various exclusive globular species of *Lobivia*, columnar species of *Harrisia*, *Cleistocactus* and *Vatricania* and one leafy species of *Pereskia*. Bioregion 13 is in Eastern Brazil and is characterised by exclusive species of various columnar genera such as *Pilosocereus*, *Micranthocereus*, *Leocereus* and *Stephanocereus* and also globular species of *Melocactus*.

Other bioregions rich in exclusive species (> 15 spp) are: 2 (Andean region/Chile), 4 (Andean region/Peru), 9 (SE/S Brazil), 21 (Caribbean), 5 (Andean region/Peru and Ecuador), 11 (extreme S Brazil and Uruguay), 10 (N South America) and 7 (central South America) (Table 4). Bioregions 7 and 9 also include areas with high phylogenetic diversity (Table 4; Amaral et al., 2022). Bioregions 23 (Central America), 8 and 12 (both in Central Brazil), 20 (USA/Florida), 6 (Small area in Peru) and 24 (Galapagos) include less than 50 species and less than 15 exclusive species. However, all of them include a cactus flora with at least 10 species and/or a high proportion of exclusive species. For example, Bioregion 24 (Galapagos) includes only three species of cacti but all of them are exclusive to this bioregion. In 19 of the 24 bioregions all indicative cactus species are restricted to their respective bioregion (Table S5).

The hierarchical solution capturing larger bioregions showed three bioregions in the first level (Fig. 3b): (1) a northern bioregion comprising North and Central America and northern South America; (2) one bioregion in the Galapagos Islands; and (3) one bioregion including eastern, western and southern South America (Fig. 3). Interzones are highlighted in central Brazil, Bolivia, Paraguay, Peru, Ecuador and Venezuela, corresponding to areas in bioregions 5, 6, 7, 8, 10 and 12 (Fig. 3a). Alternative first level hierarchical solutions (Fig. S8) show a further east/west division in South America. In the second level (Fig. 3c) 13 bioregions emerged in a scheme generally similar to the non-hierarchical solution (Fig. 2), although with a few larger bioregions. The areas corresponding to Bioregions 15 and 16 emerge in a single bioregion, as well as areas corresponding to bioregions 8 and 12. A single Mexican bioregion includes areas corresponding to bioregions 14, 17 and 18 and a single Andean bioregion includes areas corresponding to bioregions 2, 4 and 5. A large bioregion is stretched along Argentina, Uruguay, Bolivia and Paraguay, including areas corresponding to bioregions 1, 3, 7 and 11.

The two level hierarchical schemes obtained incorporating phylogenetic data (Fig. 4) are very similar to the previous schemes based only on occurrence data (non-hierarchical and hierarchical; Figs. 2 and 3). The first level includes 15 bioregions with Central and South American bioregions delimited similarly to the second level scheme and a single bioregion encompassing the whole Mexican and Southwestern USA center of diversity. Main differences include a different outline for bioregion 2, including an extreme Argentinean portion clustering with the southern Chilean region, and a portion of the Dominican Republic clustering with a large continental bioregion including bioregions 22, 23 and 10. In the second level, five Mexican bioregions emerge with differences only in a larger bioregion 18, including the southern portion of bioregion 14 and the tip of the Yucatán peninsula.

## DISCUSSION

### The pros and cons of specimen and observation data

Major advantages of biodiversity distribution data from public repositories or data providers, such as GBIF, are their free availability and the ease and speed of access. Verification and validation of data mostly overcomes issues such as taxonomic misidentification and georeferencing errors that are putatively problematic in such public databases. Data validation steps also help to identify other common issues of spatial analyses such as incomplete coverage due to geographically and taxonomically biased sampling (Maldonado et al., 2015; Meyer et al., 2016). Biased coverage can be particularly problematic for groups such as cacti, for which collection deficit must be particularly considered when estimating levels of diversity (Anderson, 2001; Taylor and Zappi, 2004). Nevertheless, the distribution, richness and density patterns obtained here agree with patterns previously documented and thoroughly verified and validated in the literature for Cactaceae in the Neotropical region (Barthlott and Hunt, 1993; Barthlott et al., 2015; Goettsch et al., 2015).

For analysing our large dataset, including both preserved specimens and human observation point occurrences for all Neotropical cacti, the automated filtering was particularly useful, and allowed faster standardized, and replicable data validation and cleaning (Zizka et al., 2019; Zizka et al., 2020b). Comparison to documented knowledge and estimations based on checklists and species lists validated by specialists (e.g., Hunt et al., 2006; Barthlott et al., 2015; Hunt, 2016; Korotkova et al., 2021) were also valuable to validating our dataset and analysing results. Similar validation through comparison has been performed previously for assessing limitations in inferring biodiversity patterns from point occurrences in public databases for other plant groups (e.g., Yesson et al., 2007; Maldonado et al., 2015).

Data obtained from preserved specimens allowed the identification of three main centers of diversity in arid and semiarid regions of Mexico/USA, Brazil and around the Andes, which were already largely acknowledged (Barthlott and Hunt 1993; Taylor, 1997; Anderson, 2001) and had also emerged as cores of high endemism and phylogenetic diversity in Amaral et al. (2022). The center of diversity in the Andean region was particularly underestimated by preserved specimen data and it was enhanced in record density and species richness with the addition of human observation data. This deficit of coverage of preserved specimens in the Andean region was particularly highlighted around the central and southern Andes and may have been caused either due to collection deficit or missing herbarium data in our dataset. The need for scientific collection in the central Andean region of Peru and Bolivia to shed light on the taxonomic and conservation status of several cacti has been highlighted by specialists (e.g., Taylor, 1997) and although there have been efforts to cover this knowledge deficit (Barthlott et al., 2015; Goettsch et al., 2015, 2018), preserved specimen data alone might still be insufficient for estimating levels of diversity in this region.

Our study documented higher species richness for each center of diversity when human observation data was added, as we obtained occurrence data exclusively from human observations for 108 species (these had missing data for herbarium specimens in our dataset). These included many species with particularly restricted distributions, and this may have hampered their collection and preservation, particularly if they were also located in remote places and associated with rarity. Forty-seven of these species have been classified as threatened or Data Deficient (IUCN, 2022), and 31 of these are indeed poorly represented in scientific collections, for example a few small and globular cacti of *Copiapoa*, *Parodia* and *Gymnocalycium* from Chile, Bolivia, and Argentina, respectively. This is also the case for several species of *Mammillaria* in Mexico. It is therefore clear that the data made available from human observations can be particularly valuable for complementing existing records and knowledge of these species.

On the other hand, all open georeferenced locality data for rare and threatened species have the potential to facilitate extractive collections and illegal trade; for this reason, some institutions avoid publicizing occurrence data online for threatened taxa or generalize the records so that exact locations cannot be determined (Chapman, 2020). The obscure coordinates option in *i*Naturalist is one of the tools to avoid the open publication of precise locality data for threatened species (www.inaturalist.org/, information given under threatened taxa, searched in September 2022). However, when there are many observations or records for a narrowly distributed taxon, the mapped occurrence (such as in *i*Naturalist or GBIF) can allow the inference of localities and environments where populations occur. The conservation status tag highlighted for each threatened taxon listed on *i*Naturalist nevertheless is a valuable information tool that reaches the community. That together with further information and recommendations can help to raise awareness and to potentially expand the network of people involved in conservation actions and to aid in the protection of taxa.

Another set of species sampled exclusively by human observations in our dataset (16 spp, 15%) had a relatively good sampling of preserved specimens (more than 10 records per species) in GBIF and a few of them had metadata with geographic coordinates. Data from these specimens were not included in our dataset as they were eliminated during automated and manual cleaning due to the quality criteria we followed. Human observations, such as those obtained from citizen science platforms, are often georeferenced with acceptable accuracy and migrated directly to the platform and subsequently to GBIF, without further editing of the coordinates. This reduces the potential for generating errors associated with coordinates, making it a potentially powerful source of data for spatial analysis. In contrast, different sources of errors with the geographical coordinates can accumulate in scientific collections-based data. Herbarium specimens often have geographical coordinates data manually transcribed from GPS data collected in the field or, for older specimens, the coordinates may appear in the labels but may lack precision or are subject to digitizing errors (Soltis, 2017). Furthermore, the metadata may undergo subsequent migrations through different databases until it is included in GBIF, when errors can accumulate.

Lack of data in GBIF may also have affected our assessment of species underrepresented by preserved specimens. Plant specimens deposited at most of the world’s about 3,000 herbaria have not yet been fully digitised, including many small herbaria with important collections of the local flora. Additional material may be available in those herbaria for several species for which the metadata were not digitized or were not included in GBIF, and therefore are not included in our dataset. This digitization gap has been identified previously as a major knowledge impediment for other groups of plants, such as the Fabaceae (Yesson et al., 2007) and may be the reason for the particularly incomplete coverage for the Andean region in the preserved specimen dataset. Even though the representation of herbaria from countries in the Global South in public databases has increased in the last 20 years (GBIF, www.gbif.org, searched in October 2021), occurrences from regional Andean herbaria remain underrepresented in our dataset. Of 89 herbaria in Peru, Bolivia, Chile and Argentina with potential collections for cacti registered in Index Herbariorum (http://sweetgum.nybg.org/science/ih/, searched in October 2021), 25 had datasets for preserved specimens in GBIF, 14 had occurrences for cacti and only seven of them included geographical coordinates. In contrast, the collections of herbaria in other South American countries such as Brazil and Colombia were well represented; we retrieved data from 73 Brazilian herbaria and from 21 Colombian herbaria. We also retrieved data for all countries in the Andean region from major and specialized herbarium collections (such as K, MO, NY, F, DES, E and B), which include duplicates from regional herbaria and may therefore represent their collections relatively well.

Future initiatives to enhance coverage of records for cacti in public databases should support not only an increase in sampling for areas and taxa potentially under-represented in collections, but also the digitization of specimens in local herbaria, and inclusion of the metadata produced in online databases. Nevertheless, our results support that, for spatial studies, publicly available human observation records, curated both scientifically and by citizens, represent a relevant tool to complement the occurrence data obtained only from preserved specimens. It is important to note, however, that such specimens are the basis of taxonomic research, providing information and material for many other applications, such as species description, morphological studies, DNA extraction, studies on herbivory and other ecological relationships, and biochemical studies (Folk and Siniscalchi, 2021). As discussed in Troudet et al. (2018), visual observations cannot be used as a basis for these types of studies and therefore should not be seen as a substitute for preserved specimens.

Although poor data quality and rapid identification error propagation are among the main concerns for data based on citizen science (Kosmala et al., 2016), initiatives to promote and support periodic validation and curation of data by specialists in citizen science databases can help mitigate these issues and make these data more readily reliable for research (Troudet et al., 2018). The manual cleaning performed in this work based on mapped point occurrences for individual species allowed a thorough verification of the citizen-science-based data, and most of the problematic records (excluded) were related to species commonly cultivated outside of their native range. Nevertheless, the data from preserved specimens also included occurrence records of cultivated cacti, which were excluded during our manual cleaning.

Cacti are particularly well documented in human observation datasets in GBIF, and it is possible that data from visual observations may be not as informative and complementary for other plant groups. Many species of cacti are severely threatened and targeted in conservation action plans, including government-based projects and initiatives to document the image and occurrence of species observed in natural areas such as in national parks (Table S1). Also, citizen science observation databases include a large amount of data for cacti, because they are charismatic and attractive to many people, including amateurs with a good knowledge of species identification and several of them work actively to curate observations. Cactaceae has more than ∼520,000 research grade observations in *i*Naturalist (i.e., independently validated by at least two experts or “knowledgeable people”, sensu *i*Naturalist) for 1,719 species in the wild (of a total of ∼1,900 in the family). A few charismatic groups show comparable results, such as Aizoaceae with ∼75,000 observations for 1,265 species (∼1,722 in the family). For the highly ornamental Orchidaceae, ∼820,000 research grade observations are listed but for only 8,307 of the ∼26,000 species in the family. Numbers are lower and even less representative for other charismatic and economically important families such as Myrtaceae, which had ∼123,000 research grade observations for 1,890 species (∼5,900 in the family), Bromeliaceae with ∼91,000 research grade observations for 1271 species (∼3,728 in the family), Arecaceae with ∼86,000 for 777 species (∼2,457 in the family), and Annonaceae with ∼39,000 for 356 species (∼2,430 in the family) (www.inaturalist.org/, searched July 6th, 2023).

Our data also suggest that citizen science, at least for the collection of species occurrences, is still in its infancy in many Neotropical countries, particularly in Central and South America, even though there are website translations to several languages including Spanish. Only 7% of all *i*Naturalist observations in our dataset were recorded in countries other than Mexico and USA. A similar pattern, with South American countries showing fewer contributions to observations in general, is revealed in searches for verifiable observations of plant families in *i*Naturalist, even when the size of countries is considered: 184,577 were made in Ecuador, 126,040 in Brazil, 55,954 in Peru and 51,302 in Bolivia, compared to 1,054,004 in Mexico and 17,289,264 in USA (www.inaturalist.org/, searched October 16th, 2021). Even though *i*Naturalist data contributed to enhancing the density of records in the Andean region in our dataset, the number of observations in general was lower for Andean countries when compared to northern America countries, to a degree highly disproportionate to the difference in the number of species. Further efforts from the scientific community and government initiatives in Central and South America can help to engage citizens to contribute to databases, promoting the importance of science and basic science education in general.

### The bioregions of cacti

As succulent plants with life history strategies and adaptations associated with dry environments, cacti are expected to be concentrated in arid and semi-arid climate regions (Gregory-Wodzicki, 2000; Arakaki et al., 2011). Our data support this, with higher species richness and major bioregions centered around the Sonoran (Bioregion 16), Baja Californian (Bioregion 15), Chihuahuan (Bioregion 19) and Atacaman (bioregions 1, 2, 3, and 4) deserts and on the Caatinga dry forest (Bioregion 13). Although a few cactus species inhabit the Amazonian domain (such as the epiphytic and wide-ranging *Rhipsalis baccifera* and *Epiphyllum phyllanthus*), the extra-Amazonian pattern is pronounced and suggests that the Amazonian tropical forest may also act as a barrier for the dispersal of most cacti as it does for many taxa from open habitats (Colli-Silva, 2021). In contrast, the Atlantic Forest, which is the second largest Neotropical forest, is home to several cactus lineages, such as the epiphytic Rhipsalideae, which can also inhabit patches of humid forests in the Andean Yungas (Calvente et al., 2011; Barthlott et al., 2015).

From a Neotropic-wide perspective, topography also seems to influence regional patterns observed for cacti. Elevation, in combination with many factors such as dynamics of air masses and humidity, temperature, and exposure to direct sunlight (also associated with the presence or absence of forested environments) and soil have been associated with the distribution, phylogenetic diversity and endemism patterns for cacti and other plant groups (Guerrero et al., 2011; Moeslund et al., 2013; Amaral et al., 2022). Andean and Mesoamerican higher elevations seem to play a major role in shaping diversity and distribution patterns for cacti, and consequently may be associated with the delimitation of bioregions. A major bioregion (Bioregion 1) with high species richness and endemism, is stretched along higher elevation Andean regions adjacent to five smaller ones (bioregions 2, 3, 4, 5 and 6), which may also have been shaped under elevation dynamics. The three most species-rich bioregions (bioregions 19, 17 and 18) are also under the influence of elevation dynamics in Mexico, spreading along the Central Mexican Plateau, interspersed between Sierra Madre Occidental, Sierra Madre Oriental, the Trans-Mexican Volcanic Belt and Sierra Madre del Sur.

Fire may also be a relevant factor in our bioregionalization scheme. Closely linked to the distribution, composition and structure of Neotropical savannas, fire influences the occurrence of several lineages, such as Melastomataceae (e.g., Microlicieae), Fabaceae (e.g., *Mimosa*, *Andira*), Malvaceae (e.g., *Eriotheca*), Asteraceae (e.g., *Viguiera*), and Poaceae (e.g., *Actinocladum*) among many others (Soderstrom, 1981; Fritsch et al., 2004; Simon et al., 2009; Simon and Pennington, 2012). A pattern of seasonal fires in the Cerrado seems not to favor the occurrence of cacti, which commonly do not have adaptations to resist these fire cycles as is the case for other succulent taxa (Pennington et al., 2009). Frequently, cacti in Cerrado occur in rocky outcrops that act as refuges away from the fires (Taylor and Zappi, 2004; Lavor et al., 2018). These conditions may be the key element shaping bioregions 8 and 12, which occur in marginal disjunct patches around core Cerrado regions in Central Brazil.

Although much of the diversity of cacti is confined within continental land masses, bioregional patterns are also created by marine island systems. In the Pacific, there is the isolated bioregion 24 placed away from the continent on the Galapagos islands. In the Caribbean, the bioregional delimitation suggests a complex scenario with disjunct patches of the same bioregions on different islands and on the continent (bioregions 20 and 21 and 22, 23 and 10 as supported by phylogenetic data) and with a further subdivision of bioregion 21 in three smaller bioregions in the third level of an alternative hierarchical solution for this region (Fig. S7). The distance among land masses in the Caribbean is much shorter, thereby aiding dispersal, but other factors may also have facilitated a dynamic dispersal of cacti in that region. Pleistocene glacial cycles as well as major meteorological events such as hurricanes may have increased the opportunity for dispersal among islands, also influencing the diversification of several Pleistocene aged lineages such as *Consolea*, *Harrisia*, *Melocactus* and *Pilosocereus* (Frank et al., 2013; Lavor et al., 2018; Majure et al., 2021; Majure et al., 2022).

Major bioregions 9, 10 and 13 and bioregion 11 extend through continuous units that have emerged in previous Neotropical regional classifications based on patterns observed for whole vegetations and biotas (Griffith et al., 1998; Omernick and Griffith, 2014; Dinerstein et al., 2017; IBGE, 2019; Morrone et al., 2022; Fig. 5). Among them, bioregion 11, which extends through the Uruguayan savanna (Dinerstein et al, 2017; Fig. 5b, 5c) in Brazil, Uruguay, and Argentina, has a remarkable cactus flora with several globose species of *Parodia* and *Frailea* inhabiting open grassland habitats. Bioregions 9 and 10 occupy extensive areas of humid forests included in the Tropical and Subtropical Moist Broadleaf Forests biome (Dinerstein et al., 2017; Fig. 5b). Bioregion 10 also extends in disjunct patches into the adjacent Tropical & Subtropical Grasslands, Savannas & Shrublands biome (Dinerstein et al., 2017; Fig. 5b), including forest taxa which can expand their range into riparian forests along a savanna-like matrix, or lithophyte taxa that occur in rock outcrops embedded in forests or savannas.

Although Bioregion 13 corresponds mostly to dry and seasonally marked areas of the Caatinga (an ecoregion in Dinerstein et al., 2017 and Griffith et al., 1998 and a province in Morrone et al., 2022; Fig. 5), it also expands through adjacent provinces and ecoregions; it includes ecotonal areas between Caatinga and Cerrado and between Caatinga and Atlantic Forest already included in the expanded definition of the Caatinga according to the most recent evaluation of the Instituto Brasileiro de Geografia e Estatística (IBGE, 2019). It also includes hotspot areas of the campo rupestre provinces Chapada Diamantina and Southern Espinhaço in the Espinhaço Range (Colli-Silva et al., 2019) with remarkable endemism for cacti (Amaral et al. 2022).

### Hierarchical schemes and comparison with other classifications

The North/South pattern shown for Cactaceae in first levels of the hierarchical scheme (Fig. 3) surpasses the scale of traditional large scale biogeographical divisions for Cactaceae, which describe three main large diversity centers. The Andean center and the Eastern Brazil center appear within a continuous large southern bioregion pointing to some degree of biogeographical connection between them. Alternative first level hierarchical solutions show a further east/west subdivision of this large southern bioregion with contrasting relationships for the central area in between them. Those areas in Argentina, Bolivia and Paraguay can appear either clustered to the East or to the West and occupy large stretches of transition zones (Fig. S8). Other large transition zones among bioregions occur along the Andes, in northern South America and in Central America.

Lower level subdivisions along this central South American transition zone were inconsistent in our results, also when phylogenetic data was integrated, leading to conflicting solutions with divergent merging and subdivisions of bioregions. A more conservative approach would be to recognize a single bioregion including bioregions 1, 3 and 7 and 11 and another single bioregion including bioregions 8 and 12 (as in Fig.3). The same applies to bioregion 6 that may be merged into bioregion 5. Nevertheless, the non-hierarchical scheme presented here in which these areas are divided into smaller bioregions (Fig. 2) offers the most consistent outcome for finer-scale analysis based on our dataset, although the fit of this particular scheme might need to be better evaluated when focusing on specific analyses, such as ancestral area reconstruction for narrowly distributed lineages.

Overall, phylogenetic data supported solutions based exclusively on occurrence data for our dataset. Although using different methodological approaches, Holt et al. (2013) and Slik et al. (2020) found more pronounced discrepancies when phylogenetic data was integrated in the classification of zoological regions and tropical forests, respectively. Amaral et al. (2022) also found similar patterns between taxon richness and phylogenetic diversity for Cactaceae and nearly half of our non-hierarchical scheme include areas where they detected high phylogenetic diversity. Nevertheless, incorporating phylogenetic information here was a powerful tool to highlight that most bioregions outlined are also meaningful evolutionarily. Differences observed in the outline of a few bioregions highlights the phylogenetic uniqueness or a narrower definition for a few bioregions. In some cases, such as for bioregions 14 and 21, some of their bordering portions were clustered into more widespread bioregions by their shared occurrence of widespread taxa. With many exclusive lineages such as *Eriosyce*, *Copiapoa* and *Eulychnia*, the phylogenetic uniqueness of Bioregion 2 was marked, as it emerged distinct from northern Andean bioregions.

The bioregionalization scheme proposed here for Neotropical cacti is singular in many aspects. Although overall results are comparable to biogeographic units based on whole terrestrial biotas (e.g., Olson et al., 2001; Morrone, 2014; Dinerstein et al., 2017; Morrone et al., 2022; Fig. 5), the exact delineation and regionalization scales are different in most cases. For example, the second level of the hierarchical scheme (Fig. 3) shows a single bioregion stretching along the Andean Chile and Peru what is comparable to some extent of the South American transition zone (Morrone et al., 2022), although north, south and east borders are much narrower. In Mesoamerica and in the Andean region, bioregions for cacti are too wide to fit ecoregions and too narrow to fit the biomes in Dinerstein et al. (2017). Bioregion 3 for example contains the Central Andean Dry Puna and part of the Central Andean Puna and several other ecoregions of five different biomes (Table 4). Nevertheless, Bioregion 2 nearly corresponds to the Chilean Matorral ecoregion within the Mediterranean Forests, Woodlands & Scrub biome.

In general, the same is observed in Mesoamerican and Andean regions for the ecoregions of Griffith et al. (1998), with cacti bioregions being larger than third level ecoregions and narrower than second level ecoregions. Comparison with provinces and dominions or zones of Morrone et al. (2022) lead to a similar interpretation: overall cacti bioregions are more generalized than provinces and narrower than dominions. In Brazil on the other hand, bioregions 9, 13 and 12/8 show a similar outline of operational units defined in previous schemes (Table 4, Figs. 3 and 4).

These differences in relation to other schemes indicate that the distribution of cacti seem to exceed the general borders of the whole biota and for others it may be more restricted. Such singularity in the distribution patterns in a group with unique life history strategies and adaptations is expected, as these plants may deal differently with extreme factors that limit the distribution of other groups. For instance, dry regions and habitats are mostly barriers to the dispersal of forest taxa, but corridors and cradles of diversification for dry-adapted taxa such as cacti. Richness and endemism analyses conducted with Neotropical Bromeliaceae also resulted in some degree of singularity when compared to macroecological schemes based on plant and animal data, likely due to differences in methodology and evolutionary history of the taxa examined (Zizka et al., 2019). Differences in source and availability of data, criteria used and scale, which are critical for the exact delineation of biogeographic units, are other likely reasons for differences among bioregionalization schemes (Kreft & Jetz, 2010). However, the bioregionalization obtained here is highly consistent with global biodiversity patterns previously described for cacti.

Most centers of diversity recognized by Barthlott et al. (2015) emerged inside corresponding bioregions. The Caatinga center is contained within Bioregion 13. The Mata Atlântica center is within Bioregion 9, the southern central Andes center is stretched along bioregions 1 and 3, and the Puebla-Oaxaca center is in Bioregion 18. The very broad Chihuahua center on the other hand, which has the greatest species and generic richness, ranges across three bioregions in Mexico (bioregions 16, 17 and 19). Bioregion 16 also includes the Sonora-Sinaloan center and Bioregion 14 contains the Jalisco center. Although our results are overall consistent at this continental scale, smaller scale regionalization, particularly on Andean and Mexican regions with a heterogeneous landscape and topography associated with high species richness and endemism, may improve the resolution of regional boundaries, and improve the fit of the scheme, especially for restricted and narrowly distributed taxa. An improved dataset with the addition of more high-quality data on species occurrence for still poorly documented areas and taxa and with more data from herbaria yet to be digitized may bring complementary results on a more refined smaller scale.

## CONCLUSIONS

Citizen-science observations can greatly complement traditional specimen-derived data for biodiversity information, particularly for taxa such as cacti that are challenging to collect due to intrinsic or legal factors. We propose a new bioregionalization scheme for Cactaceae which is overall comparable to other regionalization schemes proposed but complements them with clearer and better-supported biogeographical boundaries. These results provide a new comparative backbone to support the investigation of patterns and processes underlying the biogeographic history and evolutionary diversification of lineages of cacti and of the dry Neotropical flora.

## Supporting information

Phylogenetic trees 1 and 2

non-hierarchical scheme shapefile

## ACKNOWLEDGMENTS

The authors thank Pró-reitoria de Pesquisa of UFRN and the Systematics and Evolution graduate program for providing a visiting professor grant to AA; to the collectors, observators and institutions that provided and made available the metadata from preserved specimen and human observations of cacti used in this study; to Rhian Smith for science and language editing; to Leonardo Versieux for suggestions on the manuscript; and to three anonymous reviewers for valuable suggestions that helped improve this article. AA acknowledges financial support from the Swedish Research Council (2019-05191) and the Royal Botanic Gardens, Kew. APAS acknowledges a PhD fellowship from CAPES. We acknowledge individual credits for species imagens in Figure 2 and thank: Jan Doležal (*<u>E. aurea</u>*), Miguel A. Casado (*<u>C. cinerascens</u>*), Martin Lowry (*<u>L. calorubra</u>*), Christian Bravard (*<u>C. brevistylus</u>*), Oscar Johnson (*<u>B. microsperma</u>*), Manuel Roncal (*<u>C. substerile</u>*), María Zeta (*<u>A. rhodotrichum</u>*), Thales Santos (*<u>C. pierre-braunianus</u>*), Juliana Zuluaga-Carrero (*<u>M. schatzlii</u>*), Martin Coronel Varela (*<u>P. scopa</u>*), Mattheus Mota (*<u>C. bicolor</u>*), William Bruno (*<u>M. zehntneri</u>*), Juan Ramón Manjarrez (*<u>S. martinezii</u>*), Vince Scheidt, CNPS (*<u>E. maritimus</u>*), Marc Faucher (*<u>S. erectocentrus</u>*), Aaron Balam (*<u>S. phyllacanthus</u>*), Ignacio Torres García (*<u>M. pectinifera</u>*), Ana Luisa Fernández Fuentes (*<u>E. longisetus</u>*), Eric M Powell (*<u>O. drummondii</u>*), Yolanda M. Leon (*<u>M. lemairei</u>*), Luis Humberto Vicente Rivera (*<u>S. pteranthus</u>*), Yamilette Herrera Estévez (*<u>E. hookeri</u>*), William Stephens (*<u>B. nesioticus</u>*).

## AUTHOR CONTRIBUTIONS

AC, APAS, AA and MRF designed the study. AC, DE and APAS analysed data and results and AC, APAS and FAC wrote the first draft. All authors reviewed, commented, and contributed intellectually to improve the analyses and presented results and text.

## DATA AVAILABILITY STATEMENT

Datasets used in these studies contain the location of many endangered or legally protected species and therefore are not publicly archived. Original datasets may be requested through direct contact with the authors. Phylogenetic trees and shapefiles of the non-hierarchical scheme are available in the supplementary material and may be used freely.

## FIGURE LEGENDS

**Figure 1.** Density of records (a-d) and species richness (e-h) of Neotropical Cactaceae on adaptive 0.5° to 2° grid cells, based on: (a, e) Preserved specimens only, (b, f) Human observations excluding *i*Naturalist, (c, g) *i*Naturalist only, (d, h) complete dataset including preserved specimens and human observations. Darker shades highlight major diversity centers for the family: North and Central America, Andean Region, Eastern Brazil. Histograms of records from 1945 to 2021 for preserved specimen (i) and human observation (j) datasets.

**Figure 2.** Bioregionalization for Neotropical cacti based on preserved specimens and human observations resulting in 24 bioregions highlighted in colors and numbered. The images show the indicative species or the most common (in bold) species for each bioregion: 1. *Echinopsis aurea*, 2. *Copiapoa cinerascens*, 3. *Lobivia calorubra*, 4. *Corryocactus brevistylus*, 5. *Browningia microsperma*, 6. *Calymmanthium substerile*, 7. *Acanthocalycium rhodotrichum*, 8. *Cereus pierre-braunianus*, 9. *Rhipsalis pulchra*, 10. *Melocactus schatzlii*, 11. *Parodia scopa*, 12. *Cereus bicolor*, 13. *Melocactus zehntneri*, 14. *Stenocereus martinezii*, 15. *Echinocereus maritimus*, 16. *Sclerocactus erectocentrus*, 17. *Stenocactus phyllacanthus*, 18. *Mammillaria pectinifera*, 19. *Echinocereus longisetus*, 20. *Opuntia drummondii*, 21. *Melocactus lemairei*, 22. *Selenicereus pteranthus*, 23. ***Epiphyllum hookeri***, 24. *Brachycereus nesioticus* (images of species obtained from *i*Naturalist, individual credits in acknowledgements). Note that some bioregions contain multiple species that are equally indicative or common (see scores in item S5 below).

**Figure 3:** Bioregionalization for Neotropical cacti based on a hierarchical solution method and variable Markov time using preserved specimens and human observations. Two hierarchical levels show three bioregions in the first level (a, b) and 13 bioregions in the second level (c). Decreased opacity in cells (a) highlight interzones (fuzzy borders) between bioregions of the first level.

**Figure 4:** Bioregional schemes for the Neotropics: maps adapted from (a) Morrone et al. (2022), (b) Dinerstein et al. (2017), (c) Griffith et al. (1998), and (d) EPA (2022). Biomes abbreviations: **DXS**: Deserts and Xeric Shrublands; **FGS**: Flooded Grasslands and Savannas; **Ma:** Mangroves; **MFWS**: Mediterranean Forest Woodlands, and Scrub; **MGS:** Montane Grasslands and Shrublands; **TBMF**: Temperate Broadleaf and Mixed Forests; **TCF:** Temperate Coniferous Forests; **TGSS**: Temperate Grasslands, Savannas and Shrublands; **TSCF:** Tropical and Subtropical Coniferous Forests; **TSDBF**: Tropical and Subtropical Dry Broadleaf Forests; **TSGSS**: Tropical and Subtropical Grasslands, Savannas and Shrubland; **TSMBF**: Tropical and Subtropical Moist Broadleaf Forests.

