## Supplementary material for "Spiny but photogenic: amateur sightings complement herbarium specimens to reveal the bioregions of cacti": Figures and supplementary material

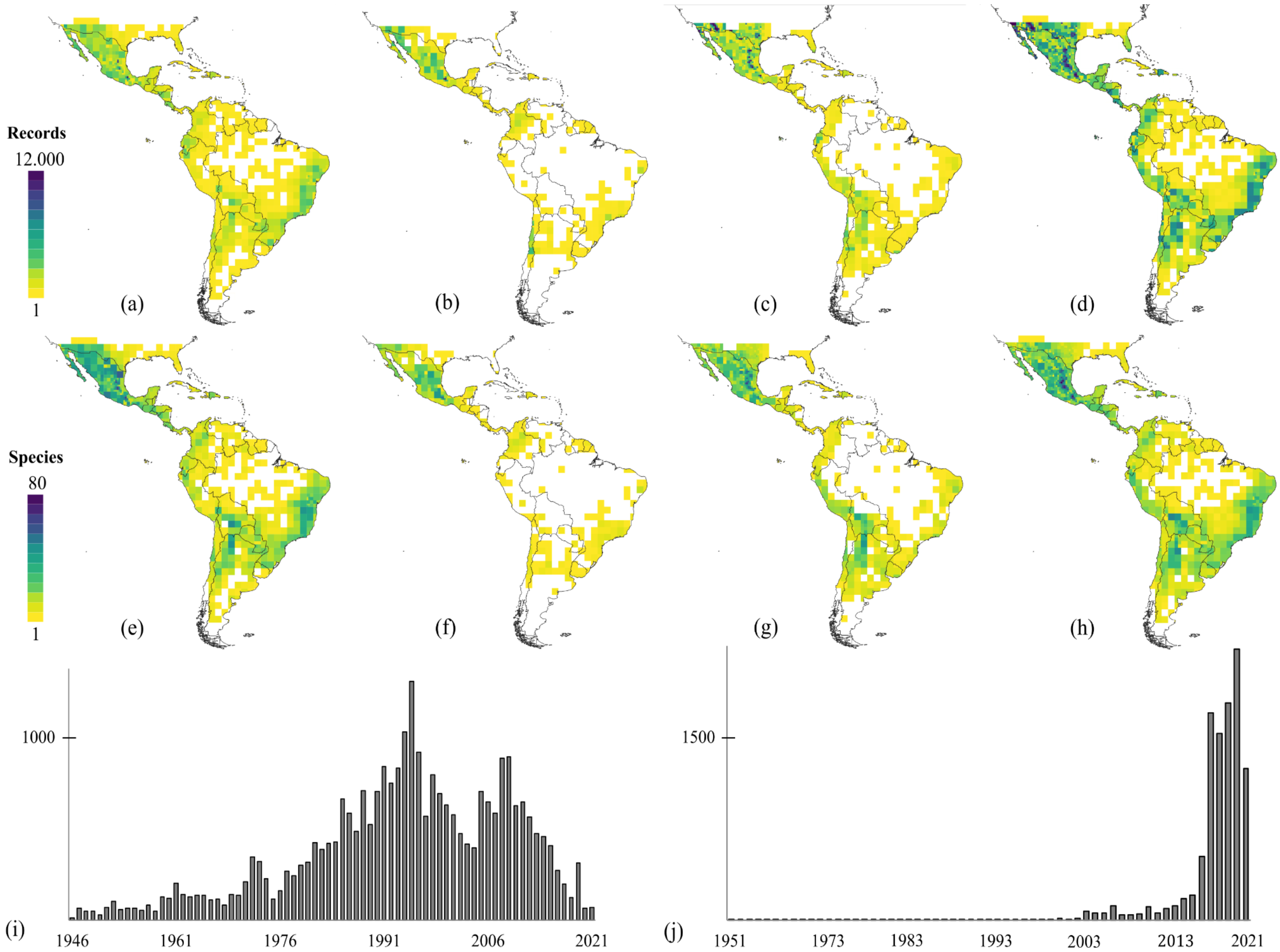

**Figure 1**: Density of records (a-d) and species richness (e-h) of Neotropical Cactaceae on adaptive 0.5° to 2° grid cells, based on: (a, e) Preserved specimens only, (b, f) Human observations excluding *i*Naturalist, (c, g) *i*Naturalist only, (d, h) complete dataset including preserved specimens and human observations. Darker shades highlight major diversity centers for the family: North and Central America, Andean Region, Eastern Brazil. Histograms of records from 1945 to 2021 for preserved specimen (i) and human observation (j) datasets.

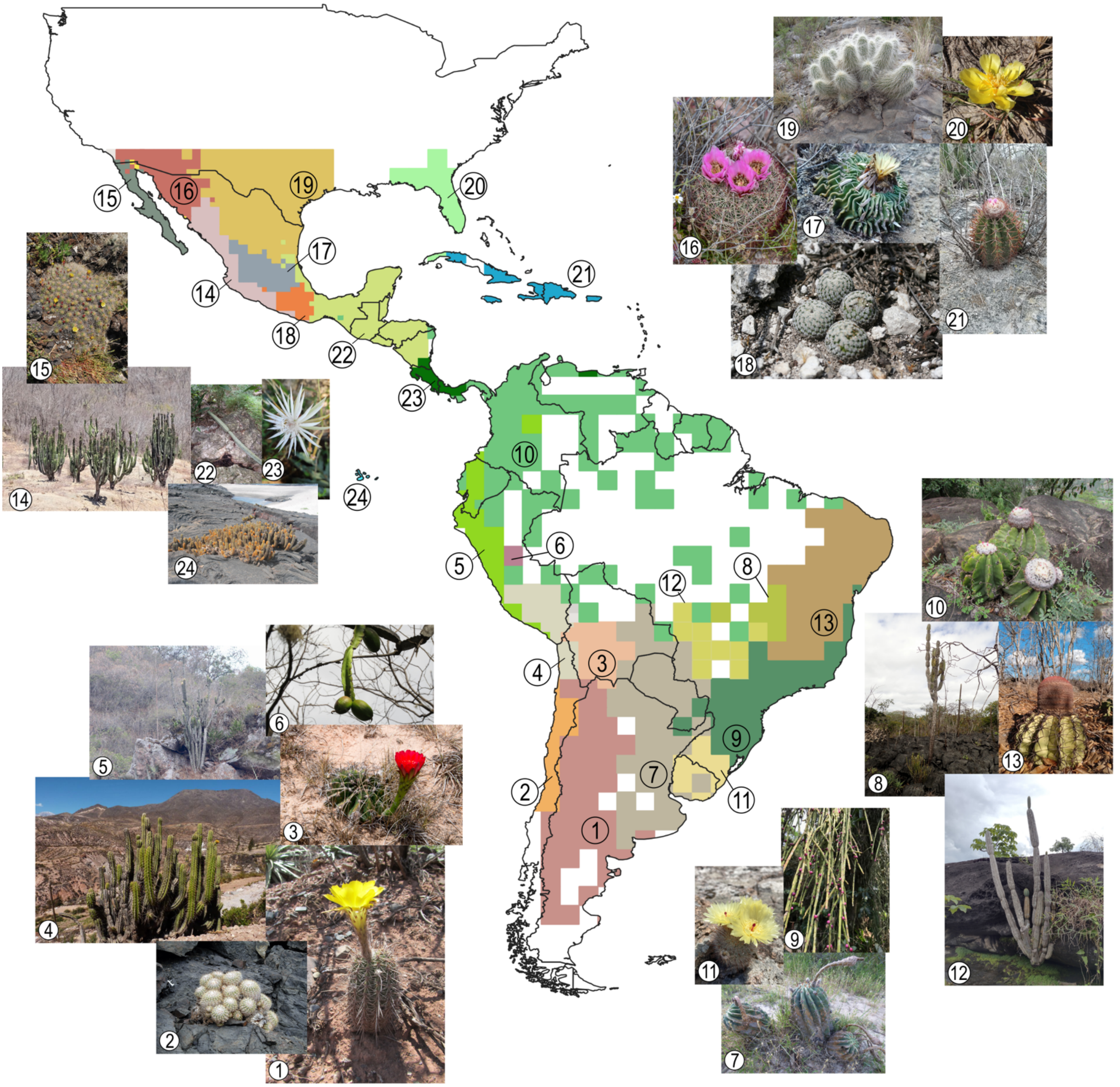

**Figure 2:** Bioregional scheme for Neotropical cacti based on a non-hierarchical solution method using preserved specimens and human observations: 24 bioregions are highlighted in colors and numbered. The images show the indicative species or the most common (in bold) species for each bioregion: 1. *Echinopsis aurea*, 2. *Copiapoa cinerascens*, 3. *Lobivia calorubra*, 4. *Corryocactus brevistylus*, 5. *Browningia microsperma*, 6. *Calymmanthium substerile*, 7. *Acanthocalycium rhodotrichum*, 8. *Cereus pierre-braunianus*, 9. *Rhipsalis pulchra*, 10. *Melocactus schatzlii*, 11. *Parodia scopa*, 12. *Cereus bicolor*, 13. *Melocactus zehntneri*, 14. *Stenocereus martinezii*, 15. *Echinocereus maritimus*, 16. *Sclerocactus erectocentrus*, 17. *Stenocactus phyllacanthus*, 18. *Mammillaria pectinifera*, 19. *Echinocereus longisetus*, 20. *Opuntia drummondii*, 21. *Melocactus lemairei*, 22. *Selenicereus*

*pteranthus*, 23. ***Epiphyllum hookeri***, 24. *Brachycereus nesioticus* (images of species obtained from *i*Naturalist, individual credits in acknowledgements). Note that some bioregions contain multiple species that are equally indicative or common (see scores in table S5).

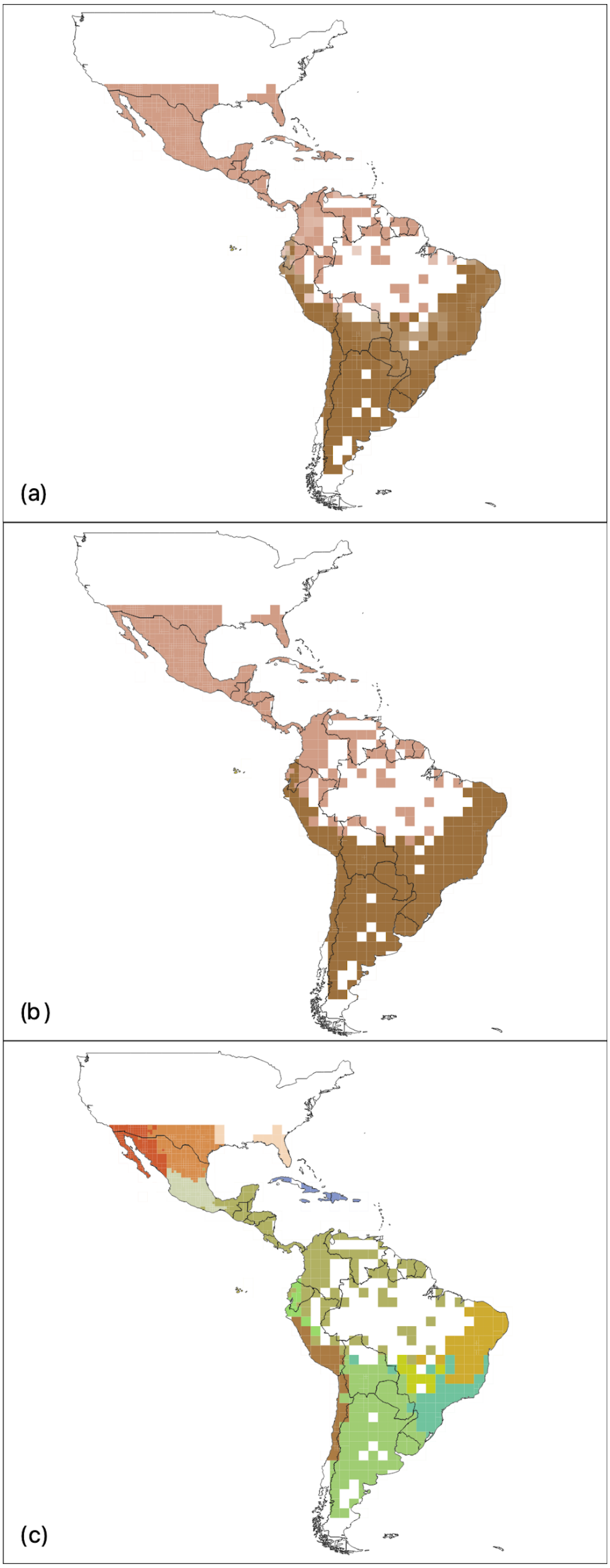

**Figure 3:** Bioregionalization for Neotropical cacti based on a hierarchical solution method and variable Markov time using preserved specimens and human observations. Two hierarchical levels show three bioregions in the first level (a, b) and 13 bioregions in the second level (c). Decreased opacity in cells (a) highlights interzones (fuzzy borders) between bioregions of the first level.

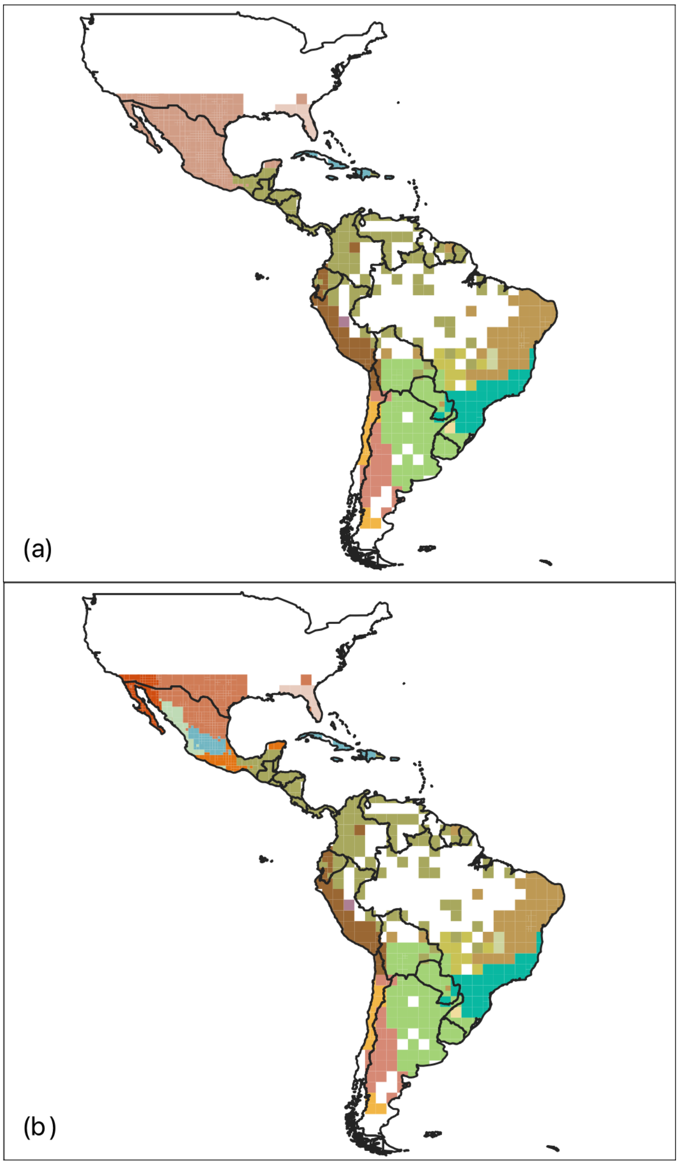

**Figure 4:** Bioregionalization for Neotropical cacti based on a hierarchical solution method incorporating phylogenetic information, preserved specimens and human observations. Two hierarchical levels show 15 bioregions in the first level (a) and 33 bioregions in the second level (b).

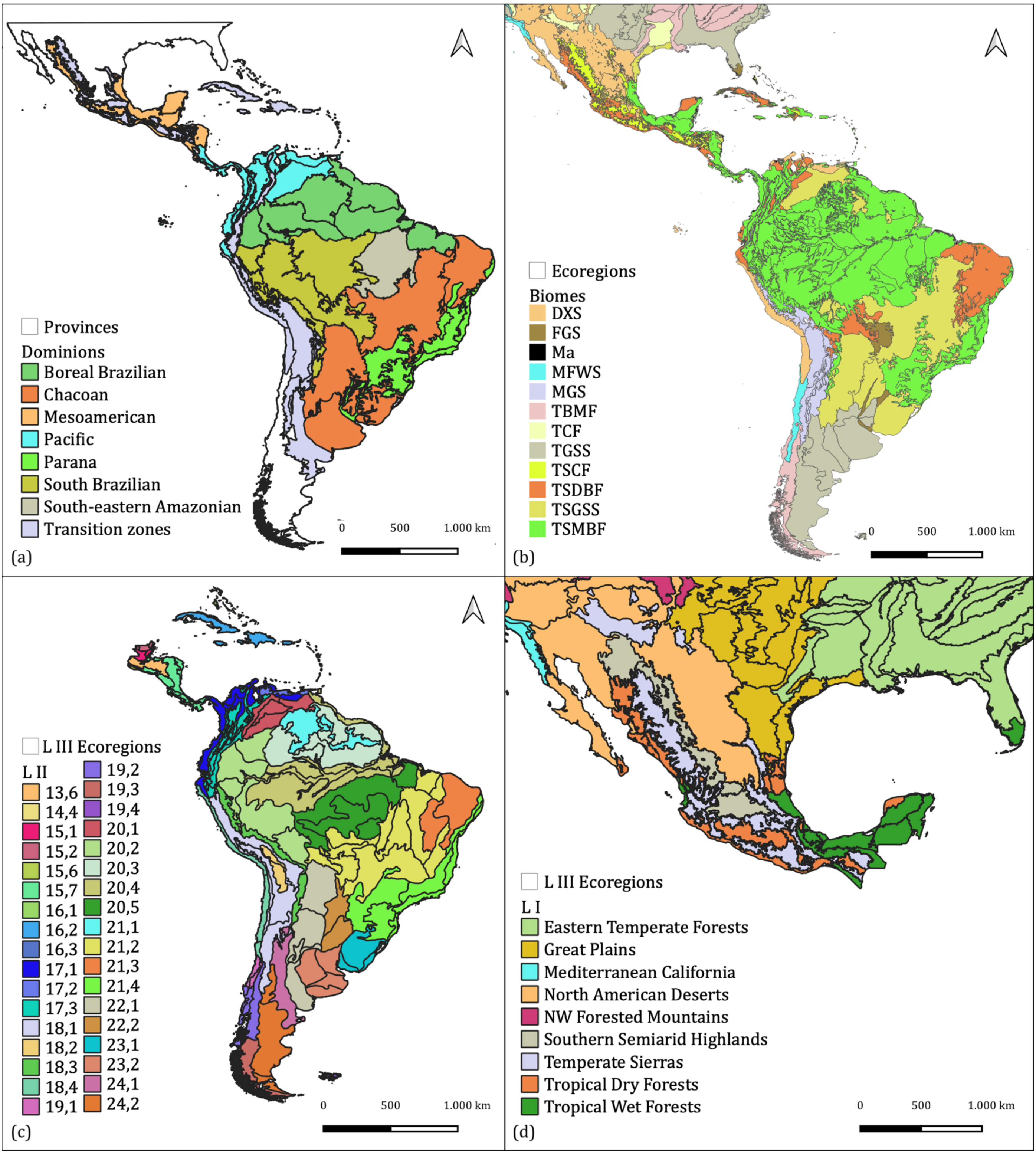

**Figure 5:** Bioregional schemes for the Neotropics: maps adapted from (a) Morrone et al. (2022), (b) Dinerstein et al. (2017), (c) Griffith et al. (1998), and (d) EPA (2022). Biomes abbreviations: **DXS**: Deserts and Xeric Shrublands; **FGS**: Flooded Grasslands and Savannas; **Ma:** Mangroves; **MFWS**: Mediterranean Forest Woodlands, and Scrub; **MGS:** Montane Grasslands and Shrublands; **TBMF**: Temperate Broadleaf and Mixed Forests; **TCF:** Temperate Coniferous Forests; **TGSS**: Temperate Grasslands, Savannas and Shrublands; **TSCF:** Tropical and Subtropical Coniferous Forests; **TSDBF**: Tropical and Subtropical Dry Broadleaf Forests;

**TSGSS**: Tropical and Subtropical Grasslands, Savannas and Shrubland; **TSMBF**: Tropical and Subtropical Moist Broadleaf Forests.

### Supplementary Material

S1. List of major datasets (datasets with less than 2 records were excluded from this list) and number of records for each dataset in human observations data, excluding *i*Naturalist.

| Dataset | Number of records |
|---|---|
| Comisión nacional para el conocimiento y uso de la biodiversidad | 6,265 |
| Pl@nNet automatically identified occurrences | 2,090 |
| Ministerio del Medio Ambiente de Chile | 805 |
| Sistema de Información sobre Biodiversidad de la Colombia - SiB | 330 |
| naturgucker | 117 |
| fieldmuseum | 107 |
| The Vascular Plant Collection at the Botanische Staatssammlung München | 74 |
| MEXU/Flora de Oaxaca-UNIBIO, IBUNAM | 42 |
| Sistema de Informação sobre a Biodiversidade Brasileira - SiBBr | 41 |
| Taxon occurrence data for the FungalRoot database | 41 |
| Pl@ntNet observations | 39 |
| BLM - National Landscape Monitoring Framework - Plants | 36 |
| Instituto de Investigación de Recursos Biológicos Alexander von Humboldt | 33 |
| Observation.org, Nature data from around the World | 29 |
| Pontificia Universidad Católica de Valparaíso | 24 |
| Bioimages-Vanderbilt University, | 22 |
| International Barcode of Life project (iBOL) | 22 |
| Ministerio del Ambiente - Ecuador | 22 |
| Instituto Amazónico de Investigaciones Científicas - SINCHI | 20 |
| ArOBIS Centro Nacional Patagónico | 19 |
| Fundación Natura Colombia | 19 |
| Laboratorio de Invasiones Biológicas (LIB) | 17 |
| UMS PatriNat (OFB-CNRS-MNHN), Paris | 15 |
| ipttest.conabio | 11 |
| Florida Museum of Natural History | 8 |
| Plantas Vasculares del Uruguay (Observaciones Mai & Rossado) | 8 |
| UNIBIO, IBUNAM | 8 |
| Universidad Icesi | 8 |
| Arizona State University Biocollections | 7 |
| Corporación Paisajes Rurales | 7 |
| UNIBIO, IBUNAM | 7 |
| Carbones del Cerrejón Limited | 6 |
| Conservation International | 6 |

| | |
|---|---|
| Oleoducto Bicentenario | 5 |
| Parques Nacionales Naturales de Colombia | 5 |
| Patrimonio Natural Fondo para la Biodiversidad y Áreas Protegidas "Patrimonio Natural" | 5 |
| BioExpertise Engine (BEE) | 4 |
| Instituto para la Investigación y la Preservación del Patrimonio Cultural y Natural del Valle del Cauca - INCIVA | 4 |
| Promigas S.A E.S.P | 4 |
| Instituto de Investigaciones Ambientales del Pacífico John Von Neumann - IIAP | 3 |
| Parques Nacionales Naturales de Colombia-Fauna y Flora de Cinaruco - 2014- 2016 | 3 |
| api.datacite | 2 |
| CVC - Corporación Autónoma Regional del Valle del Cauca | 2 |
| Universidad de Ciencias Aplicadas y Ambientales U.D.C.A | 2 |

S2. List of species with point occurrence records in this study. The “source of records" column highlights species recorded only in one basis of records, either preserved specimens or human observations (blank means the species was recorded in both). The “changes column” describe the magnitude of changes made during the final manual cleaning (less than 10 records excluded = Minor, more than 10 records excluded = Major); in the final manual cleaning we examined and revised individual species distribution maps resulting from automated cleaning aiming mostly to minimize errors due to the registered occurrence of cultivated specimens.

| **Species** | **Source of records** | **Changes (final manual cleaning)** |
|---|---|---|
| *Acanthocalycium leucanthum* | | |
| *Acanthocalycium rhodotrichum* | | Minor |
| *Acanthocalycium spiniflorum* | Only preserved specimens | |
| *Acanthocalycium thionanthum* | Only preserved specimens | |
| *Acanthocereus chiapensis* | | |
| *Acanthocereus cuixmalensis* | Only preserved specimens | |
| *Acanthocereus fosterianus* | | |
| *Acanthocereus hirschtianus* | | |
| *Acanthocereus macdougallii* | Only preserved specimens | |
| *Acanthocereus maculatus* | Only preserved specimens | |
| *Acanthocereus oaxacensis* | | |
| *Acanthocereus rosei* | | |
| *Acanthocereus tepalcatepecanus* | Only *i*Naturalist | |
| *Acanthocereus tetragonus* | | |
| *Acharagma aguirreanum* | Only preserved specimens | |
| *Acharagma roseanum* | | |

| | | |
|---|---|---|
| *Airampoa ayrampo* | | |
| *Airampoa corrugata* | Only preserved specimens | |
| *Airampoa erectoclada* | Only preserved specimens | |
| *Airampoa microdisca* | Only preserved specimens | |
| *Aporocactus flagelliformis* | | Minor |
| *Aporocactus martianus* | Only preserved specimens | |
| *Ariocarpus agavoides* | | |
| *Ariocarpus bravoanus* | Only preserved specimens | |
| *Ariocarpus fissuratus* | | |
| *Ariocarpus kotschoubeyanus* | | |
| *Ariocarpus retusus* | | |
| *Ariocarpus scaphirostris* | | |
| *Ariocarpus trigonus* | | |
| *Armatocereus cartwrightianus* | | |
| *Armatocereus godingianus* | | |
| *Armatocereus laetus* | Only preserved specimens | |
| *Armatocereus matucanensis* | | |
| *Armatocereus procerus* | | |
| *Armatocereus rauhii* | | |
| *Armatocereus riomajensis* | Only *i*Naturalist | |
| *Arrojadoa bahiensis* | | |
| *Arrojadoa dinae* | Only preserved specimens | Minor |
| *Arrojadoa marylanae* | Only preserved specimens | |
| *Arrojadoa penicillata* | | |
| *Arrojadoa rhodantha* | | |
| *Arthrocereus glaziovii* | | |
| *Arthrocereus melanurus* | | |
| *Arthrocereus rondonianus* | Only preserved specimens | |
| *Astrophytum asterias* | | |
| *Astrophytum capricorne* | | |
| *Astrophytum caput-medusae* | Only *i*Naturalist | |
| *Astrophytum coahuilense* | Only preserved specimens | |
| *Astrophytum myriostigma* | | Minor |
| *Astrophytum ornatum* | | Minor |
| *Austrocactus bertinii* | | |
| *Austrocactus ferrarii* | | |
| *Austrocactus philippii* | Only *i*Naturalist | |
| *Austrocactus spiniflorus* | | |
| *Austrocylindropuntia cylindrica* | | |

| | | |
|---|---|---|
| *Austrocylindropuntia floccosa* | | |
| *Austrocylindropuntia pachypus* | | |
| *Austrocylindropuntia shaferi* | | |
| *Austrocylindropuntia subulata* | | |
| *Austrocylindropuntia vestita* | | |
| *Aylostera deminuta* | Only preserved specimens | |
| *Aztekium hintonii* | Only preserved specimens | |
| *Aztekium ritteri* | | |
| *Bergerocactus emoryi* | | |
| *Blossfeldia liliputana* | | |
| *Borzicactus acanthurus* | | |
| *Borzicactus fieldianus* | Only *i*Naturalist | |
| *Borzicactus icosagonus* | | |
| *Borzicactus leonensis* | Only preserved specimens | |
| *Borzicactus neoroezlii* | Only preserved specimens | |
| *Borzicactus pachycladus* | Only *i*Naturalist | |
| *Borzicactus plagiostoma* | Only preserved specimens | |
| *Borzicactus sepium* | | |
| *Borzicactus sextonianus* | | |
| *Borzicactus sulcifer* | Only *i*Naturalist | |
| *Borzicactus tenuiserpens* | Only preserved specimens | |
| *Brachycereus nesioticus* | | |
| *Brasilicereus markgrafii* | | |
| *Brasilicereus phaeacanthus* | Only preserved specimens | |
| *Brasiliopuntia brasiliensis* | | Major |
| *Browningia altissima* | | |
| *Browningia candelaris* | | |
| *Browningia hernandezii* | Only preserved specimens | |
| *Browningia hertlingiana* | Only *i*Naturalist | |
| *Browningia microsperma* | | |
| *Browningia pilleifera* | | |
| *Browningia utcubambensis* | Only preserved specimens | |
| *Calymmanthium substerile* | Only preserved specimens | |
| *Carnegiea gigantea* | | |
| *Castellanosia caineana* | | |
| *Cephalocereus apicicephalium* | | |
| *Cephalocereus columna-trajani* | | |

| | | |
|---|---|---|
| *Cephalocereus euphorbioides* | | |
| *Cephalocereus fulviceps* | | |
| *Cephalocereus macrocephalus* | | |
| *Cephalocereus mezcalaensis* | | |
| *Cephalocereus nudus* | | |
| *Cephalocereus polylophus* | | |
| *Cephalocereus sanchez-mejoradae* | Only preserved specimens | |
| *Cephalocereus scoparius* | | |
| *Cephalocereus senilis* | | |
| *Cephalocereus tetetzo* | | |
| *Cereus aethiops* | | |
| *Cereus albicaulis* | | |
| *Cereus bicolor* | | Minor |
| *Cereus fernambucensis* | | Major |
| *Cereus hankeanus* | | |
| *Cereus hexagonus* | | |
| *Cereus hildmannianus* | | Major/ all excluded |
| *Cereus insularis* | | |
| *Cereus jamacaru* | | Minor |
| *Cereus lanosus* | Only preserved specimens | |
| *Cereus mirabella* | Only preserved specimens | |
| *Cereus phatnospermus* | Only preserved specimens | Minor |
| *Cereus pierre-braunianus* | | |
| *Cereus repandus* | | Minor |
| *Cereus saddianus* | Only preserved specimens | |
| *Cereus spegazzinii* | | |
| *Cereus stenogonus* | | |
| *Cereus vargasianus* | Only *i*Naturalist | |
| *Cereus yungasensis* | | |
| *Chamaecereus saltensis* | | |
| *Chamaecereus silvestrii* | | Minor |
| *Chamaecereus stilowianus* | Only *i*Naturalist | |
| *Cipocereus bradei* | Only preserved specimens | |
| *Cipocereus crassisepalus* | Only preserved specimens | |
| *Cipocereus laniflorus* | Only preserved specimens | |
| *Cipocereus minensis* | | |
| *Cipocereus pusilliflorus* | Only preserved specimens | |
| *Cleistocactus baumannii* | | |

| | | |
|---|---|---|
| *Cleistocactus brookeae* | | |
| *Cleistocactus buchtienii* | | |
| *Cleistocactus candelilla* | | |
| *Cleistocactus hyalacanthus* | | Minor |
| *Cleistocactus laniceps* | | Minor |
| *Cleistocactus luribayensis* | Only *i*Naturalist | |
| *Cleistocactus morawetzianus* | Only *i*Naturalist | |
| *Cleistocactus parviflorus* | | |
| *Cleistocactus pungens* | Only *i*Naturalist | |
| *Cleistocactus reae* | | |
| *Cleistocactus samaipatanus* | Only preserved specimens | |
| *Cleistocactus smaragdiflorus* | | |
| *Cleistocactus strausii* | Only preserved specimens | |
| *Cochemiea albicans* | | |
| *Cochemiea armillata* | | |
| *Cochemiea barbata* | | |
| *Cochemiea blossfeldiana* | | |
| *Cochemiea boolii* | | |
| *Cochemiea capensis* | | |
| *Cochemiea cerralboa* | Only preserved specimens | |
| *Cochemiea conoidea* | | |
| *Cochemiea dioica* | | |
| *Cochemiea grahamii* | | |
| *Cochemiea guelzowiana* | | |
| *Cochemiea halei* | | Minor |
| *Cochemiea hutchisoniana* | | |
| *Cochemiea insularis* | | |
| *Cochemiea macdougallii* | Only preserved specimens | |
| *Cochemiea mainiae* | | |
| *Cochemiea palmeri* | | |
| *Cochemiea phitauiana* | | |
| *Cochemiea pondii* | | |
| *Cochemiea poselgeri* | | |
| *Cochemiea saboae* | | |
| *Cochemiea schumannii* | | |
| *Cochemiea tetrancistra* | | |
| *Cochemiea theresae* | | |
| *Cochemiea thornberi* | | |
| *Cochemiea wrightii* | | |
| *Coleocephalocereus aureus* | Only preserved specimens | |

| | |
|---|---|
| *Coleocephalocereus buxbaumianus* | Only preserved specimens |
| *Coleocephalocereus fluminensis* | |
| *Coleocephalocereus goebelianus* | Only preserved specimens |
| *Coleocephalocereus pluricostatus* | Only preserved specimens |
| *Coleocephalocereus purpureus* | Only preserved specimens |
| *Consolea macracantha* | |
| *Consolea moniliformis* | |
| *Consolea rubescens* | Only preserved specimens |
| *Consolea spinosissima* | |
| *Copiapoa calderana* | |
| *Copiapoa cinerascens* | |
| *Copiapoa cinerea* | |
| *Copiapoa conglomerata* | Only *i*Naturalist |
| *Copiapoa coquimbana* | |
| *Copiapoa dealbata* | |
| *Copiapoa decorticans* | Only *i*Naturalist |
| *Copiapoa echinoides* | |
| *Copiapoa fiedleriana* | Only preserved specimens |
| *Copiapoa humilis* | |
| *Copiapoa hypogaea* | |
| *Copiapoa longistaminea* | Only *i*Naturalist |
| *Copiapoa marginata* | |
| *Copiapoa megarhiza* | |
| *Copiapoa serpentisulcata* | |
| *Copiapoa solaris* | |
| *Copiapoa taltalensis* | Only preserved specimens |
| *Corryocactus aureus* | |
| *Corryocactus ayacuchoensis* | Only *i*Naturalist |
| *Corryocactus brachypetalus* | |
| *Corryocactus brevistylus* | |
| *Corryocactus chachapoyensis* | Only *i*Naturalist |
| *Corryocactus erectus* | |
| *Corryocactus melanotrichus* | |
| *Corryocactus squarrosus* | Only *i*Naturalist |
| *Corryocactus tarijensis* | Only *i*Naturalist |
| *Coryphantha clavata* | |

| | |
|---|---|
| *Coryphantha compacta* | |
| *Coryphantha cornifera* | Only preserved specimens |
| *Coryphantha delaetiana* | |
| *Coryphantha delicata* | |
| *Coryphantha difficilis* | |
| *Coryphantha durangensis* | |
| *Coryphantha echinoidea* | |
| *Coryphantha echinus* | |
| *Coryphantha elephantidens* | |
| *Coryphantha erecta* | |
| *Coryphantha georgii* | Only *i*Naturalist |
| *Coryphantha glanduligera* | |
| *Coryphantha glassii* | |
| *Coryphantha gracilis* | |
| *Coryphantha hintoniorum* | |
| *Coryphantha jalpanensis* | |
| *Coryphantha kracikii* | |
| *Coryphantha longicornis* | |
| *Coryphantha macromeris* | |
| *Coryphantha maiz-tablasensis* | |
| *Coryphantha neglecta* | Only *i*Naturalist |
| *Coryphantha nickelsiae* | |
| *Coryphantha octacantha* | |
| *Coryphantha ottonis* | |
| *Coryphantha pallida* | |
| *Coryphantha poselgeriana* | |
| *Coryphantha potosiana* | Only *i*Naturalist |
| *Coryphantha pseudoechinus* | |
| *Coryphantha pseudonickelsiae* | |
| *Coryphantha pulleineana* | Only *i*Naturalist |
| *Coryphantha pycnacantha* | |
| *Coryphantha ramillosa* | |
| *Coryphantha recurvata* | |
| *Coryphantha retusa* | |
| *Coryphantha robustispina* | |
| *Coryphantha salinensis* | Only *i*Naturalist |
| *Coryphantha sulcata* | |
| *Coryphantha tripugionacantha* | Only *i*Naturalist |

| | | |
|---|---|---|
| *Coryphantha vaupeliana* | | |
| *Coryphantha vogtherriana* | Only *i*Naturalist | |
| *Coryphantha werdermannii* | | |
| *Coryphantha wohlschlageri* | | |
| *Cumarinia odorata* | | |
| *Cumulopuntia boliviana* | | |
| *Cumulopuntia chichensis* | | |
| *Cumulopuntia leucophaea* | Only *i*Naturalist | |
| *Cumulopuntia sphaerica* | | |
| *Cylindropuntia acanthocarpa* | | |
| *Cylindropuntia alcahes* | | |
| *Cylindropuntia anteojoensis* | | |
| *Cylindropuntia arbuscula* | | |
| *Cylindropuntia bigelovii* | | |
| *Cylindropuntia californica* | | |
| *Cylindropuntia caribaea* | | |
| *Cylindropuntia cedrosensis* | | |
| *Cylindropuntia cholla* | | |
| *Cylindropuntia davisii* | | |
| *Cylindropuntia echinocarpa* | | |
| *Cylindropuntia fulgida* | | |
| *Cylindropuntia ganderi* | | |
| *Cylindropuntia hystrix* | Only preserved specimens | |
| *Cylindropuntia imbricata* | | |
| *Cylindropuntia kleiniae* | | |
| *Cylindropuntia leptocaulis* | | |
| *Cylindropuntia lindsayi* | | |
| *Cylindropuntia molesta* | | |
| *Cylindropuntia munzii* | | |
| *Cylindropuntia prolifera* | | |
| *Cylindropuntia ramosissima* | | |
| *Cylindropuntia sanfelipensis* | | |
| *Cylindropuntia santamaria* | | |
| *Cylindropuntia tesajo* | | |
| *Cylindropuntia thurberi* | | Minor |
| *Cylindropuntia tunicata* | | Major |
| *Cylindropuntia whipplei* | | |
| *Cylindropuntia wolfii* | | |
| *Deamia chontalensis* | Only preserved specimens | |
| *Deamia testudo* | | |

| | | |
|---|---|---|
| *Denmoza rhodacantha* | | |
| *Discocactus bahiensis* | Only preserved specimens | |
| *Discocactus catingicola* | Only preserved specimens | |
| *Discocactus diersianus* | Only preserved specimens | |
| *Discocactus ferricola* | Only preserved specimens | |
| *Discocactus hartmannii* | Only preserved specimens | |
| *Discocactus heptacanthus* | | |
| *Discocactus horstii* | Only preserved specimens | |
| *Discocactus placentiformis* | | |
| *Discocactus pseudoinsignis* | Only preserved specimens | |
| *Discocactus zehntneri* | Only preserved specimens | Minor |
| *Disocactus ackermannii* | | |
| *Disocactus anguliger* | | |
| *Disocactus biformis* | Only preserved specimens | |
| *Disocactus crenatus* | | |
| *Disocactus lepidocarpus* | Only preserved specimens | |
| *Disocactus macdougallii* | | |
| *Disocactus macranthus* | | |
| *Disocactus nelsonii* | | |
| *Disocactus phyllanthoides* | | Minor |
| *Disocactus quezaltecus* | Only preserved specimens | |
| *Disocactus speciosus* | | Minor |
| *Echinocactus horizonthalonius* | | |
| *Echinocactus platyacanthus* | | |
| *Echinocereus acifer* | Only preserved specimens | |
| *Echinocereus adustus* | | |
| *Echinocereus arizonicus* | | |
| *Echinocereus barthelowanus* | | |
| *Echinocereus berlandieri* | | |
| *Echinocereus bonkerae* | | |
| *Echinocereus brandegeei* | | |
| *Echinocereus bristolii* | | |
| *Echinocereus chisosensis* | Only *i*Naturalist | |
| *Echinocereus cinerascens* | | |
| *Echinocereus coccineus* | | |
| *Echinocereus dasyacanthus* | | |
| *Echinocereus engelmannii* | | |
| *Echinocereus enneacanthus* | | |
| *Echinocereus fendleri* | | |
| *Echinocereus ferreiranus* | | |

| | |
|---|---|
| *Echinocereus grandis* | |
| *Echinocereus klapperi* | |
| *Echinocereus knippelianus* | |
| *Echinocereus ledingii* | |
| *Echinocereus leucanthus* | Only preserved specimens |
| *Echinocereus longisetus* | |
| *Echinocereus mapimiensis* | |
| *Echinocereus maritimus* | |
| *Echinocereus mombergerianus* | Only preserved specimens |
| *Echinocereus nicholii* | |
| *Echinocereus nivosus* | Only *i*Naturalist |
| *Echinocereus ortegae* | |
| *Echinocereus pacificus* | Only preserved specimens |
| *Echinocereus palmeri* | |
| *Echinocereus pamanesii* | |
| *Echinocereus papillosus* | Only *i*Naturalist |
| *Echinocereus parkeri* | |
| *Echinocereus pectinatus* | |
| *Echinocereus pentalophus* | |
| *Echinocereus platyacanthus* | Only preserved specimens |
| *Echinocereus polyacanthus* | |
| *Echinocereus poselgeri* | |
| *Echinocereus primolanatus* | |
| *Echinocereus pseudopectinatus* | |
| *Echinocereus pulchellus* | |
| *Echinocereus reichenbachii* | |
| *Echinocereus rigidissimus* | |
| *Echinocereus russanthus* | |
| *Echinocereus salm-dyckianus* | |
| *Echinocereus scheeri* | |
| *Echinocereus schmollii* | |
| *Echinocereus sciurus* | |
| *Echinocereus scopulorum* | |
| *Echinocereus spinigemmatus* | Only *i*Naturalist |
| *Echinocereus stolonifer* | |
| *Echinocereus stramineus* | |
| *Echinocereus subinermis* | Only preserved specimens |
| *Echinocereus triglochidiatus* | |

| | | |
|---|---|---|
| *Echinocereus viereckii* | | |
| *Echinocereus viridiflorus* | | |
| *Echinocereus websterianus* | | |
| *Echinocereus yavapaiensis* | Only preserved specimens | |
| *Echinopsis aurea* | | |
| *Echinopsis calochlora* | | |
| *Echinopsis chrysantha* | | |
| *Echinopsis densispina* | | |
| *Echinopsis haematantha* | | |
| *Echinopsis jajoana* | | |
| *Echinopsis lamprochlora* | Only preserved specimens | |
| *Echinopsis marsoneri* | | |
| *Echinopsis oxygona* | | |
| *Echinopsis tubiflora* | | |
| *Epiphyllum cartagense* | | |
| *Epiphyllum chrysocardium* | Only preserved specimens | Minor |
| *Epiphyllum grandilobum* | Only preserved specimens | |
| *Epiphyllum hookeri* | | Minor |
| *Epiphyllum laui* | Only preserved specimens | |
| *Epiphyllum oxypetalum* | | |
| *Epiphyllum phyllanthus* | | |
| *Epiphyllum pumilum* | Only preserved specimens | |
| *Epiphyllum thomasianum* | | Minor |
| *Epithelantha bokei* | | |
| *Epithelantha micromeris* | | |
| *Eriosyce aspillagae* | Only preserved specimens | |
| *Eriosyce aurata* | | |
| *Eriosyce bulbocalyx* | Only *i*Naturalist | |
| *Eriosyce calderana* | Only preserved specimens | |
| *Eriosyce chilensis* | | |
| *Eriosyce crispa* | | |
| *Eriosyce curvispina* | | |
| *Eriosyce engleri* | | |
| *Eriosyce eriosyzoides* | Only preserved specimens | |
| *Eriosyce esmeraldana* | | |
| *Eriosyce garaventae* | | |
| *Eriosyce heinrichiana* | | |
| *Eriosyce iquiquensis* | Only *i*Naturalist | |
| *Eriosyce islayensis* | | |
| *Eriosyce kunzei* | | |

| | | |
|---|---|---|
| *Eriosyce napina* | | |
| *Eriosyce occulta* | Only preserved specimens | |
| *Eriosyce odieri* | | |
| *Eriosyce paucicostata* | | |
| *Eriosyce recondita* | | |
| *Eriosyce rodentiophila* | | |
| *Eriosyce senilis* | | |
| *Eriosyce simulans* | | |
| *Eriosyce strausiana* | | |
| *Eriosyce subgibbosa* | | |
| *Eriosyce taltalensis* | | |
| *Eriosyce umadeave* | | |
| *Eriosyce villicumensis* | Only preserved specimens | |
| *Eriosyce villosa* | | |
| *Escobaria chihuahuensis* | | |
| *Escobaria dasyacantha* | | |
| *Escobaria duncanii* | | Minor |
| *Escobaria emskoetteriana* | | |
| *Escobaria hesteri* | Only *i*Naturalist | |
| *Escobaria laredoi* | | |
| *Escobaria lloydii* | Only preserved specimens | |
| *Escobaria minima* | Only *i*Naturalist | |
| *Escobaria missouriensis* | | |
| *Escobaria robbinsorum* | Only *i*Naturalist | |
| *Escobaria sneedii* | | |
| *Escobaria tuberculosa* | | Minor |
| *Escobaria vivipara* | Only preserved specimens | |
| *Escobaria zilziana* | | |
| *Escontria chiotilla* | | |
| *Espostoa blossfeldiorum* | | |
| *Espostoa calva* | Only *i*Naturalist | |
| *Espostoa frutescens* | | |
| *Espostoa lanata* | | |
| *Espostoa melanostele* | | |
| *Espostoa mirabilis* | Only *i*Naturalist | |
| *Espostoopsis dybowskii* | Only preserved specimens | |
| *Eulychnia acida* | Only Human Observation (including *i*Naturalist) | |
| *Eulychnia breviflora* | Only Human Observation (including *i*Naturalist) | |
| *Eulychnia castanea* | Only *i*Naturalist | |

| | | |
|---|---|---|
| *Eulychnia iquiquensis* | | |
| *Facheiroa cephaliomelana* | Only preserved specimens | |
| *Facheiroa squamosa* | Only preserved specimens | |
| *Facheiroa ulei* | Only preserved specimens | |
| *Ferocactus alamosanus* | | |
| *Ferocactus chrysacanthus* | | |
| *Ferocactus cylindraceus* | | |
| *Ferocactus diguetii* | | |
| *Ferocactus echidne* | | |
| *Ferocactus emoryi* | | |
| *Ferocactus flavovirens* | | |
| *Ferocactus fordii* | | |
| *Ferocactus glaucescens* | | |
| *Ferocactus gracilis* | | |
| *Ferocactus haematacanthus* | | Minor |
| *Ferocactus hamatacanthus* | | |
| *Ferocactus herrerae* | Only preserved specimens | |
| *Ferocactus histrix* | | |
| *Ferocactus johnstonianus* | Only preserved specimens | |
| *Ferocactus latispinus* | | |
| *Ferocactus lindsayi* | Only preserved specimens | |
| *Ferocactus macrodiscus* | | |
| *Ferocactus peninsulae* | | |
| *Ferocactus pilosus* | | |
| *Ferocactus pottsii* | | |
| *Ferocactus robustus* | | |
| *Ferocactus tiburonensis* | Only preserved specimens | |
| *Ferocactus townsendianus* | Only preserved specimens | |
| *Ferocactus uncinatus* | Only preserved specimens | |
| *Ferocactus viridescens* | | |
| *Ferocactus wislizeni* | | |
| *Frailea buenekeri* | Only preserved specimens | |
| *Frailea castanea* | Only preserved specimens | |
| *Frailea cataphracta* | Only preserved specimens | |
| *Frailea chiquitana* | Only *i*Naturalist | |
| *Frailea fulviseta* | Only preserved specimens | |
| *Frailea gracillima* | | |
| *Frailea mammifera* | Only preserved specimens | |
| *Frailea phaeodisca* | Only preserved specimens | |
| *Frailea pumila* | Only preserved specimens | |

| | | |
|---|---|---|
| *Frailea pygmaea* | | |
| *Frailea schilinzkyana* | | |
| *Geohintonia mexicana* | | |
| *Grusonia aggeria* | Only preserved specimens | |
| *Grusonia bradtiana* | | |
| *Grusonia bulbispina* | Only preserved specimens | |
| *Grusonia emoryi* | Only preserved specimens | |
| *Grusonia grahamii* | Only preserved specimens | |
| *Grusonia invicta* | Only preserved specimens | |
| *Grusonia kunzei* | Only preserved specimens | |
| *Grusonia marenae* | Only preserved specimens | |
| *Grusonia moelleri* | Only preserved specimens | |
| *Grusonia parishii* | Only preserved specimens | |
| *Grusonia reflexispina* | Only preserved specimens | |
| *Grusonia robertsii* | Only preserved specimens | |
| *Grusonia schottii* | Only preserved specimens | |
| *Grusonia vilis* | Only preserved specimens | |
| *Gymnocalycium amerhauseri* | Only *i*Naturalist | |
| *Gymnocalycium andreae* | | |
| *Gymnocalycium anisitsii* | | |
| *Gymnocalycium baldianum* | Only preserved specimens | |
| *Gymnocalycium bayrianum* | | |
| *Gymnocalycium calochlorum* | Only *i*Naturalist | |
| *Gymnocalycium capillense* | | |
| *Gymnocalycium castellanosii* | | |
| *Gymnocalycium chiquitanum* | Only *i*Naturalist | |
| *Gymnocalycium denudatum* | Only preserved specimens | |
| *Gymnocalycium friedrichii* | Only preserved specimens | |
| *Gymnocalycium gibbosum* | Only preserved specimens | Minor |
| *Gymnocalycium hossei* | Only *i*Naturalist | |
| *Gymnocalycium hyptiacanthum* | Only preserved specimens | |
| *Gymnocalycium kieslingii* | Only preserved specimens | |
| *Gymnocalycium marsoneri* | | |
| *Gymnocalycium mesopotamicum* | Only preserved specimens | |
| *Gymnocalycium mihanovichii* | | |
| *Gymnocalycium monvillei* | | Minor |

| | | |
|---|---|---|
| *Gymnocalycium mostii* | | |
| *Gymnocalycium pflanzii* | | |
| *Gymnocalycium pugionacanthum* | | |
| *Gymnocalycium quehlianum* | | |
| *Gymnocalycium ragonesii* | Only preserved specimens | |
| *Gymnocalycium reductum* | | |
| *Gymnocalycium rhodantherum* | Only *i*Naturalist | |
| *Gymnocalycium ritterianum* | Only *i*Naturalist | |
| *Gymnocalycium robustum* | | |
| *Gymnocalycium saglionis* | | |
| *Gymnocalycium schickendantzii* | | Minor |
| *Gymnocalycium schroederianum* | | |
| *Gymnocalycium spegazzinii* | | |
| *Gymnocalycium striglianum* | Only *i*Naturalist | |
| *Gymnocalycium taningaense* | Only *i*Naturalist | |
| *Haageocereus acranthus* | | |
| *Haageocereus chilensis* | | |
| *Haageocereus decumbens* | | |
| *Haageocereus kagenekii* | | |
| *Haageocereus platinospinus* | | |
| *Haageocereus tenuis* | Only preserved specimens | |
| *Haageocereus versicolor* | | |
| *Harrisia adscendens* | | Minor |
| *Harrisia bonplandii* | Only preserved specimens | |
| *Harrisia eriophora* | | |
| *Harrisia gracilis* | | Minor |
| *Harrisia martinii* | | |
| *Harrisia pomanensis* | | Minor |
| *Harrisia regelii* | | Minor |
| *Harrisia tetracantha* | | Minor |
| *Harrisia tortuosa* | | Minor |
| *Hatiora salicornioides* | | Minor |
| *Homalocephala parryi* | | |
| *Homalocephala polycephala* | | |
| *Homalocephala texensis* | | |
| *Isolatocereus dumortieri* | | |
| *Jasminocereus thouarsii* | | |
| *Kadenicarpus horripilus* | Only preserved specimens | |

| | | |
|---|---|---|
| *Kadenicarpus pseudomacrochele* | Only preserved specimens | |
| *Kimnachia ramulosa* | Only preserved specimens | |
| *Kroenleinia grusonii* | | |
| *Lemaireocereus hollianus* | | |
| *Lemaireocereus lepidanthus* | Only preserved specimens | |
| *Leocereus bahiensis* | Only preserved specimens | |
| *Lepismium cruciforme* | | Minor |
| *lepismium houlletianum* | | Minor |
| *Lepismium lorentzianum* | | |
| *Lepismium lumbricoides* | | Minor |
| *Lepismium warmingianum* | | |
| *Leptocereus arboreus* | | |
| *Leptocereus leonii* | Only *i*Naturalist | |
| *Leptocereus nudiflorus* | | |
| *Leptocereus paniculatus* | | |
| *Leptocereus quadricostatus* | | |
| *Leptocereus sylvestris* | | |
| *Leptocereus undulosus* | | |
| *Leptocereus velozianus* | Only preserved specimens | |
| *Leptocereus weingartianus* | | |
| *Leuchtenbergia principis* | | |
| *Leucostele atacamensis* | | |
| *Leucostele chiloensis* | Only Human Observation (including *i*Naturalist) | |
| *Leucostele deserticola* | Only preserved specimens | |
| *Leucostele terscheckii* | | |
| *Leucostele tunariensis* | Only *i*Naturalist | |
| *Leucostele werdermanniana* | Only *i*Naturalist | |
| *Leuenbergeria aureiflora* | Only preserved specimens | |
| *Leuenbergeria bleo* | | Minor |
| *Leuenbergeria guamacho* | | |
| *Leuenbergeria lychnidiflora* | | |
| *Leuenbergeria marcanoi* | Only preserved specimens | |
| *Leuenbergeria portulacifolia* | Only preserved specimens | |
| *Leuenbergeria quisqueyana* | | |
| *Leuenbergeria zinniiflora* | Only preserved specimens | Minor/ all records in Mexico excluded |
| *Lobivia ancistrophora* | | |
| *Lobivia arachnacantha* | Only *i*Naturalist | |
| *Lobivia ayopayana* | Only *i*Naturalist | |
| *Lobivia backebergii* | | |

| | | |
|---|---|---|
| *Lobivia bridgesii* | | |
| *Lobivia caineana* | | |
| *Lobivia calorubra* | | |
| *Lobivia chrysochete* | | |
| *Lobivia cinnabarina* | Only *i*Naturalist | |
| *Lobivia ferox* | | |
| *Lobivia hertrichiana* | Only *i*Naturalist | |
| *Lobivia lateritia* | Only *i*Naturalist | |
| *Lobivia mamillosa* | | |
| *Lobivia maximiliana* | | |
| *Lobivia obrepanda* | | |
| *Lobivia pampana* | | |
| *Lobivia pamparuizii* | | Minor |
| *Lobivia pentlandii* | | |
| *Lobivia pugionacantha* | | |
| *Lobivia rauschii* | Only *i*Naturalist | |
| *Lobivia schieliana* | Only *i*Naturalist | |
| *Lobivia taratensis* | Only *i*Naturalist | |
| *Lobivia tegeleriana* | Only *i*Naturalist | |
| *Lobivia tiegeliana* | Only *i*Naturalist | |
| *Lophocereus gatesii* | Only preserved specimens | |
| *Lophocereus marginatus* | | Minor |
| *Lophocereus schottii* | | |
| *Lophophora diffusa* | | |
| *Lophophora williamsii* | | |
| *Lymanbensonia brevispina* | Only preserved specimens | |
| *Lymanbensonia crenata* | Only preserved specimens | |
| *Lymanbensonia incachacana* | | |
| *Maihuenia patagonica* | | |
| *Maihuenia poeppigii* | | |
| *Maihueniopsis conoidea* | Only preserved specimens | |
| *Maihueniopsis crassispina* | Only *i*Naturalist | |
| *Maihueniopsis darwinii* | | |
| *Maihueniopsis glomerata* | | |
| *Maihueniopsis hickenii* | | |
| *Maihueniopsis minuta* | | |
| *Maihueniopsis ovata* | | |
| *Maihueniopsis platyacantha* | Only *i*Naturalist | |
| *Mammillaria albicoma* | | |
| *Mammillaria albiflora* | Only *i*Naturalist | |

| | | |
|---|---|---|
| *Mammillaria albilanata* | | |
| *Mammillaria aureilanata* | Only *i*Naturalist | |
| *Mammillaria baumii* | | |
| *Mammillaria beneckei* | | |
| *Mammillaria bocasana* | | |
| *Mammillaria bocensis* | | |
| *Mammillaria boelderliana* | Only *i*Naturalist | |
| *Mammillaria bombycina* | | |
| *Mammillaria brandegeei* | | |
| *Mammillaria candida* | | |
| *Mammillaria carmenae* | | |
| *Mammillaria carnea* | | |
| *Mammillaria carretii* | Only *i*Naturalist | |
| *Mammillaria coahuilensis* | | |
| *Mammillaria columbiana* | | |
| *Mammillaria compressa* | | |
| *Mammillaria crinita* | | |
| *Mammillaria crucigera* | | |
| *Mammillaria decipiens* | | |
| *Mammillaria deherdtiana* | | |
| *Mammillaria densispina* | | |
| *Mammillaria discolor* | | |
| *Mammillaria dixanthocentron* | | |
| *Mammillaria duoformis* | | |
| *Mammillaria eichlamii* | Only preserved specimens | |
| *Mammillaria elongata* | | Minor |
| *Mammillaria eriacantha* | | |
| *Mammillaria erythrosperma* | | |
| *Mammillaria evermanniana* | | |
| *Mammillaria fittkaui* | | |
| *Mammillaria flavicentra* | Only preserved specimens | |
| *Mammillaria formosa* | | |
| *Mammillaria gasseriana* | Only preserved specimens | |
| *Mammillaria geminispina* | | Minor |
| *Mammillaria gigantea* | | |
| *Mammillaria glassii* | | |
| *Mammillaria glochidiata* | Only preserved specimens | |
| *Mammillaria grusonii* | | |
| *Mammillaria guerreronis* | | |
| *Mammillaria guillauminiana* | Only preserved specimens | |

| | | |
|---|---|---|
| *Mammillaria haageana* | | |
| *Mammillaria hahniana* | | |
| *Mammillaria hamata* | Only preserved specimens | |
| *Mammillaria hernandezii* | | |
| *Mammillaria herrerae* | | |
| *Mammillaria heyderi* | | |
| *Mammillaria huitzilopochtli* | | |
| *Mammillaria humboldtii* | Only Human Observation (including *i*Naturalist) | |
| *Mammillaria jaliscana* | | |
| *Mammillaria johnstonii* | Only *i*Naturalist | |
| *Mammillaria karwinskiana* | | |
| *Mammillaria klissingiana* | | |
| *Mammillaria knippeliana* | Only *i*Naturalist | |
| *Mammillaria kraehenbuehlii* | | |
| *Mammillaria lasiacantha* | | |
| *Mammillaria lenta* | | |
| *Mammillaria longiflora* | | |
| *Mammillaria longimamma* | | Minor |
| *Mammillaria magnifica* | | |
| *Mammillaria magnimamma* | | |
| *Mammillaria mammillaris* | Only HO excluding *i*Naturalist | |
| *Mammillaria marksiana* | | |
| *Mammillaria mathildae* | | |
| *Mammillaria mazatlanensis* | | |
| *Mammillaria melaleuca* | | |
| *Mammillaria melanocentra* | | |
| *Mammillaria mercadensis* | | |
| *Mammillaria meyranii* | | |
| *Mammillaria microhelia* | | |
| *Mammillaria moelleriana* | | |
| *Mammillaria muehlenpfordtii* | | |
| *Mammillaria mystax* | | Minor |
| *Mammillaria nana* | Only *i*Naturalist | |
| *Mammillaria napina* | | |
| *Mammillaria nivosa* | | |
| *Mammillaria nunezii* | | |
| *Mammillaria orcuttii* | | |
| *Mammillaria oteroi* | | |
| *Mammillaria parkinsonii* | | |

| | | |
|---|---|---|
| *Mammillaria pectinifera* | | |
| *Mammillaria peninsularis* | | |
| *Mammillaria pennispinosa* | | |
| *Mammillaria perbella* | | |
| *Mammillaria petrophila* | | |
| *Mammillaria petterssonii* | | |
| *Mammillaria picta* | | |
| *Mammillaria pilispina* | | |
| *Mammillaria plumosa* | | Minor |
| *Mammillaria polyedra* | | |
| *Mammillaria polythele* | | |
| *Mammillaria pottsii* | | |
| *Mammillaria pringlei* | | |
| *Mammillaria prolifera* | | |
| *Mammillaria rekoi* | | |
| *Mammillaria rettigiana* | | |
| *Mammillaria rhodantha* | | Minor |
| *Mammillaria roseoalba* | Only preserved specimens | |
| *Mammillaria sanchez-mejoradae* | | |
| *Mammillaria sartorii* | | |
| *Mammillaria schiedeana* | | |
| *Mammillaria scrippsiana* | | |
| *Mammillaria senilis* | | |
| *Mammillaria sinistrohamata* | Only preserved specimens | |
| *Mammillaria sonorensis* | | |
| *Mammillaria sphacelata* | | |
| *Mammillaria sphaerica* | | |
| *Mammillaria spinosissima* | | |
| *Mammillaria standleyi* | | |
| *Mammillaria supertexta* | | |
| *Mammillaria surculosa* | | |
| *Mammillaria tonalensis* | Only Human Observation (including *i*Naturalist) | |
| *Mammillaria uncinata* | | |
| *Mammillaria varieaculeata* | | |
| *Mammillaria vetula* | | |
| *Mammillaria wagneriana* | | |
| *Mammillaria weingartiana* | | |
| *Mammillaria wiesingeri* | | |
| *Mammillaria winterae* | | |

| | | |
|---|---|---|
| *Mammillaria xaltianguensis* | Only *i*Naturalist | |
| *Mammillaria zephyranthoides* | | |
| *Marshallocereus aragonii* | Only preserved specimens | Minor |
| *Matucana aurantiaca* | Only *i*Naturalist | |
| *Matucana formosa* | | |
| *Matucana haynei* | | |
| *Matucana madisoniorum* | Only preserved specimens | |
| *Matucana oreodoxa* | Only *i*Naturalist | |
| *Matucana weberbaueri* | | |
| *Melocactus andinus* | | |
| *Melocactus azureus* | Only preserved specimens | |
| *Melocactus bahiensis* | | |
| *Melocactus bellavistensis* | | |
| *Melocactus concinnus* | Only preserved specimens | |
| *Melocactus conoideus* | Only preserved specimens | |
| *Melocactus curvispinus* | | |
| *Melocactus deinacanthus* | Only preserved specimens | |
| *Melocactus ernestii* | | |
| *Melocactus estevesii* | Only preserved specimens | |
| *Melocactus ferreophilus* | Only preserved specimens | |
| *Melocactus glaucescens* | Only preserved specimens | |
| *Melocactus harlowii* | | |
| *Melocactus inconcinnus* | Only preserved specimens | |
| *Melocactus intortus* | | |
| *Melocactus lanssensianus* | Only preserved specimens | |
| *Melocactus lemairei* | | |
| *Melocactus levitestatus* | Only preserved specimens | |
| *Melocactus macracanthos* | | |
| *Melocactus matanzanus* | | Minor |
| *Melocactus mazelianus* | Only preserved specimens | |
| *Melocactus neryi* | | |
| *Melocactus oreas* | Only preserved specimens | Minor |
| *Melocactus pachyacanthus* | Only preserved specimens | |
| *Melocactus paucispinus* | Only preserved specimens | |
| *Melocactus peruvianus* | | |
| *Melocactus praerupticola* | Only preserved specimens | |
| *Melocactus salvadorensis* | Only preserved specimens | |
| *Melocactus schatzlii* | | |
| *Melocactus smithii* | | |
| *Melocactus violaceus* | | Minor |

| | | |
|---|---|---|
| *Melocactus zehntneri* | | |
| *Micranthocereus albicephalus* | Only preserved specimens | |
| *Micranthocereus auri-azureus* | | |
| *Micranthocereus dolichospermaticus* | Only preserved specimens | |
| *Micranthocereus estevesii* | | |
| *Micranthocereus flaviflorus* | | |
| *Micranthocereus polyanthus* | Only preserved specimens | |
| *Micranthocereus purpureus* | | |
| *Micranthocereus streckeri* | Only preserved specimens | |
| *Micranthocereus violaciflorus* | Only preserved specimens | |
| *Mila caespitosa* | | |
| *Miqueliopuntia miquelii* | | |
| *Morangaya pensilis* | | |
| *Myrtillocactus cochal* | | |
| *Myrtillocactus geometrizans* | | Minor |
| *Myrtillocactus schenckii* | | |
| *Neoraimondia arequipensis* | | |
| *Neoraimondia herzogiana* | | |
| *Neowerdermannia chilensis* | | |
| *Neowerdermannia vorwerkii* | | |
| *Nyctocereus serpentinus* | Only preserved specimens | |
| *Obregonia denegrii* | | |
| *Opuntia aciculata* | Only *i*Naturalist | |
| *Opuntia atrispina* | | |
| *Opuntia auberi* | | |
| *Opuntia aurantiaca* | Only *i*Naturalist | |
| *Opuntia aurea* | Only preserved specimens | |
| *Opuntia aureispina* | | |
| *Opuntia basilaris* | | |
| *Opuntia boldinghii* | Only preserved specimens | |
| *Opuntia bonaerensis* | Only preserved specimens | |
| *Opuntia bonplandii* | Only preserved specimens | |
| *Opuntia bravoana* | | |
| *Opuntia caracassana* | | Minor |
| *Opuntia chaffeyi* | Only preserved specimens | |
| *Opuntia chlorotica* | | |
| *Opuntia cochenillifera* | | Major/ all excluded |
| *Opuntia curassavica* | Only *i*Naturalist | |

| | | |
|---|---|---|
| *Opuntia deamii* | Only preserved specimens | |
| *Opuntia decumbens* | | |
| *Opuntia dejecta* | | |
| *Opuntia depressa* | | |
| *Opuntia discolor* | | |
| *Opuntia drummondii* | | |
| *Opuntia elata* | | Minor |
| *Opuntia elatior* | Only preserved specimens | |
| *Opuntia engelmannii* | | |
| *Opuntia excelsa* | | |
| *Opuntia feroacantha* | Only preserved specimens | |
| *Opuntia ficus-indica* | | Major/ all excluded |
| *Opuntia fragilis* | Only preserved specimens | |
| *Opuntia fuliginosa* | | |
| *Opuntia galapageia* | | |
| *Opuntia gosseliniana* | | |
| *Opuntia guatemalensis* | Only preserved specimens | Minor |
| *Opuntia guilanchi* | Only preserved specimens | |
| *Opuntia huajuapensis* | | |
| *Opuntia humifusa* | Only preserved specimens | Minor |
| *Opuntia hyptiacantha* | | |
| *Opuntia inaperta* | Only preserved specimens | |
| *Opuntia karwinskiana* | Only preserved specimens | |
| *Opuntia lagunae* | | |
| *Opuntia lasiacantha* | | |
| *Opuntia leucotricha* | | |
| *Opuntia littoralis* | | |
| *Opuntia macrocentra* | | |
| *Opuntia macrorhiza* | | |
| *Opuntia megapotamica* | | |
| *Opuntia megarrhiza* | | |
| *Opuntia microdasys* | | |
| *Opuntia monacantha* | | Minor |
| *Opuntia oricola* | | |
| *Opuntia parviclada* | | |
| *Opuntia phaeacantha* | | |
| *Opuntia pilifera* | | |
| *Opuntia pittieri* | | |
| *Opuntia polyacantha* | Only preserved specimens | |
| *Opuntia pottsii* | | |

| | | |
|---|---|---|
| *Opuntia puberula* | | |
| *Opuntia pubescens* | | Minor |
| *Opuntia pycnantha* | | |
| *Opuntia quimilo* | | |
| *Opuntia quitensis* | | |
| *Opuntia repens* | | |
| *Opuntia rioplatense* | Only *i*Naturalist | |
| *Opuntia robinsonii* | Only preserved specimens | |
| *Opuntia robusta* | | Minor |
| *Opuntia rufida* | | |
| *Opuntia scheeri* | Only preserved specimens | |
| *Opuntia schumannii* | | |
| *Opuntia setispina* | Only preserved specimens | |
| *Opuntia soederstromiana* | | Minor |
| *Opuntia spinulifera* | Only preserved specimens | |
| *Opuntia stenopetala* | | |
| *Opuntia streptacantha* | | |
| *Opuntia stricta* | | Major |
| *Opuntia strigil* | | |
| *Opuntia sulphurea* | | Minor |
| *Opuntia tapona* | Only preserved specimens | |
| *Opuntia tehuacana* | | |
| *Opuntia tomentosa* | | Minor |
| *Opuntia triacantha* | | Minor |
| *Opuntia tuna* | Only preserved specimens | Minor/ only record in Colombia excluded |
| *Opuntia velutina* | | |
| *Opuntia wilcoxii* | | |
| *Oreocereus celsianus* | Only preserved specimens | |
| *Oreocereus doelzianus* | Only preserved specimens | |
| *Oreocereus hempelianus* | Only preserved specimens | |
| *Oreocereus leucotrichus* | Only preserved specimens | |
| *Oreocereus trollii* | Only preserved specimens | |
| *Oroya borchersii* | Only preserved specimens | |
| *Pachycereus gaumeri* | | |
| *Pachycereus grandis* | | |
| *Pachycereus militaris* | | |
| *Pachycereus pecten-aboriginum* | | |
| *Pachycereus pringlei* | | |
| *Pachycereus tepamo* | | |

| | | |
|---|---|---|
| *Pachycereus weberi* | | |
| *Parodia alacriportana* | Only preserved specimens | |
| *Parodia aureicentra* | | |
| *Parodia ayopayana* | | |
| *Parodia buiningii* | Only preserved specimens | |
| *Parodia carambeiensis* | | |
| *Parodia chrysacanthion* | | |
| *Parodia columnaris* | | |
| *Parodia comarapana* | | |
| *Parodia commutans* | Only *i*Naturalist | |
| *Parodia concinna* | | |
| *Parodia crassigibba* | | |
| *Parodia curvispina* | Only preserved specimens | |
| *Parodia erinacea* | | Minor |
| *Parodia formosa* | Only preserved specimens | Minor/ only record in Brazil excluded |
| *Parodia fusca* | Only preserved specimens | |
| *Parodia gaucha* | Only preserved specimens | |
| *Parodia gibbulosa* | Only *i*Naturalist | |
| *Parodia haselbergii* | | |
| *Parodia hausteiniana* | Only *i*Naturalist | |
| *Parodia horstii* | Only preserved specimens | Minor |
| *Parodia langsdorfii* | | |
| *Parodia linkii* | Only preserved specimens | |
| *Parodia maassii* | | |
| *Parodia magnifica* | | |
| *Parodia maldonadensis* | Only *i*Naturalist | |
| *Parodia mammulosa* | | |
| *Parodia microsperma* | | Minor |
| *Parodia mueller-melchersii* | | |
| *Parodia muricata* | Only preserved specimens | |
| *Parodia nigrispina* | Only preserved specimens | |
| *Parodia nivosa* | | Minor |
| *Parodia ocampoi* | Only *i*Naturalist | |
| *Parodia otaviana* | Only *i*Naturalist | |
| *Parodia ottonis* | | |
| *Parodia oxycostata* | Only preserved specimens | |
| *Parodia prestoensis* | Only *i*Naturalist | |
| *Parodia procera* | Only *i*Naturalist | |
| *Parodia ritteri* | Only *i*Naturalist | |
| *Parodia schumanniana* | Only preserved specimens | |

| | | |
|---|---|---|
| *Parodia schwebsiana* | Only *i*Naturalist | |
| *Parodia scopa* | | |
| *Parodia stockingeri* | Only preserved specimens | |
| *Parodia stuemeri* | | |
| *Parodia subterranea* | Only *i*Naturalist | |
| *Parodia taratensis* | Only preserved specimens | |
| *Parodia tenuicylindrica* | Only preserved specimens | |
| *Parodia tuberculata* | Only *i*Naturalist | |
| *Parodia warasii* | Only preserved specimens | |
| *Pediocactus simpsonii* | Only preserved specimens | |
| *Pelecyphora aselliformis* | | |
| *Pelecyphora strobiliformis* | | |
| *Peniocereus greggii* | | |
| *Peniocereus johnstonii* | | |
| *Peniocereus lazaro-cardenasii* | | |
| *Peniocereus marianus* | Only preserved specimens | |
| *Peniocereus striatus* | | |
| *Peniocereus viperinus* | | |
| *Peniocereus zopilotensis* | Only preserved specimens | |
| *Pereskia aculeata* | | |
| *Pereskia bahiensis* | | |
| *Pereskia diaz-romeroana* | Only preserved specimens | |
| *Pereskia grandifolia* | | Major/ all excluded |
| *Pereskia horrida* | Only preserved specimens | |
| *Pereskia nemorosa* | Only preserved specimens | |
| *Pereskia sacharosa* | | |
| *Pereskia stenantha* | Only preserved specimens | |
| *Pereskia weberiana* | | |
| *Pereskiopsis aquosa* | | |
| *Pereskiopsis blakeana* | | |
| *Pereskiopsis diguetii* | | |
| *Pereskiopsis kellermanii* | | Minor |
| *Pereskiopsis porteri* | | |
| *Pereskiopsis rotundifolia* | | |
| *Pfeiffera asuntapatensis* | Only preserved specimens | |
| *Pfeiffera boliviana* | Only preserved specimens | |
| *Pfeiffera ianthothele* | | |
| *Pfeiffera monacantha* | | |
| *Pfeiffera paranganiensis* | Only *i*Naturalist | |
| *Pilosocereus albisummus* | Only preserved specimens | |

| | | |
|---|---|---|
| *Pilosocereus alensis* | | Minor |
| *Pilosocereus arrabidae* | | |
| *Pilosocereus aureispinus* | Only preserved specimens | |
| *Pilosocereus aurisetus* | | |
| *Pilosocereus azulensis* | Only preserved specimens | |
| *Pilosocereus bohlei* | Only preserved specimens | |
| *Pilosocereus brasiliensis* | | |
| *Pilosocereus catingicola* | | Minor |
| *Pilosocereus chrysacanthus* | | Minor |
| *Pilosocereus chrysostele* | | |
| *Pilosocereus collinsii* | | |
| *Pilosocereus densiareolatus* | Only preserved specimens | |
| *Pilosocereus diersianus* | Only preserved specimens | |
| *Pilosocereus flavipulvinatus* | Only preserved specimens | |
| *Pilosocereus flexibilispinus* | Only preserved specimens | |
| *Pilosocereus floccosus* | Only preserved specimens | |
| *Pilosocereus fulvilanatus* | | |
| *Pilosocereus glaucochrous* | Only preserved specimens | |
| *Pilosocereus hermii* | Only preserved specimens | |
| *Pilosocereus jauruensis* | Only preserved specimens | |
| *Pilosocereus lanuginosus* | | |
| *Pilosocereus leucocephalus* | | |
| *Pilosocereus machrisii* | | |
| *Pilosocereus magnificus* | Only preserved specimens | |
| *Pilosocereus multicostatus* | Only preserved specimens | |
| *Pilosocereus pachycladus* | | Minor |
| *Pilosocereus parvus* | Only preserved specimens | |
| *Pilosocereus pentaedrophorus* | | Minor |
| *Pilosocereus piauhyensis* | Only preserved specimens | |
| *Pilosocereus polygonus* | Only preserved specimens | |
| *Pilosocereus purpusii* | | |
| *Pilosocereus pusillibaccatus* | Only preserved specimens | |
| *Pilosocereus quadricentralis* | | |
| *Pilosocereus royenii* | | |
| *Pilosocereus splendidus* | Only preserved specimens | |
| *Pilosocereus ulei* | | |
| *Pilosocereus vilaboensis* | | |
| *Polaskia chende* | | |
| *Polaskia chichipe* | | |
| *Praecereus euchlorus* | | |

| | | |
|---|---|---|
| *Praecereus saxicola* | Only preserved specimens | Minor |
| *Pseudorhipsalis acuminata* | Only preserved specimens | |
| *Pseudorhipsalis alata* | Only preserved specimens | |
| *Pseudorhipsalis amazonica* | | |
| *Pseudorhipsalis himantoclada* | Only preserved specimens | |
| *Pseudorhipsalis lankesteri* | Only preserved specimens | |
| *Pterocactus araucanus* | Only preserved specimens | |
| *Pterocactus australis* | Only preserved specimens | |
| *Pterocactus fischeri* | | |
| *Pterocactus gonjianii* | | |
| *Pterocactus hickenii* | Only preserved specimens | |
| *Pterocactus reticulatus* | Only *i*Naturalist | |
| *Pterocactus tuberosus* | | |
| *Pterocactus valentinii* | | |
| *Punotia lagopus* | | |
| *Pygmaeocereus bylesianus* | | |
| *Quiabentia verticillata* | | |
| *Quiabentia zehntneri* | Only preserved specimens | Minor |
| *Rapicactus beguinii* | Only preserved specimens | |
| *Rapicactus mandragora* | Only preserved specimens | |
| *Rapicactus subterraneus* | Only preserved specimens | |
| *Rebutia minuscula* | | |
| *Rebutia steinbachii* | Only preserved specimens | |
| *Reicheocactus famatinensis* | Only preserved specimens | |
| *Rhipsalidopsis gaertneri* | | Minor |
| *Rhipsalidopsis rosea* | Only preserved specimens | Minor |
| *Rhipsalis agudoensis* | Only preserved specimens | |
| *Rhipsalis baccifera* | | Minor |
| *Rhipsalis burchellii* | Only preserved specimens | |
| *Rhipsalis campos-portoana* | Only preserved specimens | |
| *Rhipsalis cereoides* | Only preserved specimens | |
| *Rhipsalis cereuscula* | | |
| *Rhipsalis clavata* | | |
| *Rhipsalis crispata* | | |
| *Rhipsalis cuneata* | Only preserved specimens | |
| *Rhipsalis dissimilis* | Only preserved specimens | Minor |
| *Rhipsalis elliptica* | | |
| *Rhipsalis flagelliformis* | Only preserved specimens | |
| *Rhipsalis floccosa* | | |
| *Rhipsalis goebeliana* | Only preserved specimens | |

| | | |
|---|---|---|
| *Rhipsalis grandiflora* | | Minor |
| *Rhipsalis hoelleri* | Only preserved specimens | |
| *Rhipsalis juengeri* | Only preserved specimens | |
| *Rhipsalis lindbergiana* | | |
| *Rhipsalis mesembryanthemoides* | | Minor |
| *Rhipsalis micrantha* | | |
| *Rhipsalis neves-armondii* | Only preserved specimens | |
| *Rhipsalis oblonga* | Only preserved specimens | |
| *Rhipsalis occidentalis* | Only preserved specimens | |
| *Rhipsalis olivifera* | Only preserved specimens | Minor |
| *Rhipsalis ormindoi* | Only preserved specimens | |
| *Rhipsalis pacheco-leonis* | Only preserved specimens | |
| *Rhipsalis pachyptera* | | Minor |
| *Rhipsalis paradoxa* | | |
| *Rhipsalis pentaptera* | Only preserved specimens | |
| *Rhipsalis pilocarpa* | | |
| *Rhipsalis pulchra* | Only preserved specimens | |
| *Rhipsalis puniceodiscus* | Only preserved specimens | Minor |
| *Rhipsalis russellii* | Only preserved specimens | |
| *Rhipsalis teres* | | Minor |
| *Rhipsalis triangularis* | Only preserved specimens | |
| *Rhipsalis trigona* | | |
| *Salmonopuntia salmiana* | | |
| *Salmonopuntia schickendantzii* | Only preserved specimens | Minor |
| *Samaipaticereus corroanus* | | |
| *Schlumbergera kautskyi* | Only preserved specimens | |
| *Schlumbergera lutea* | Only preserved specimens | |
| *Schlumbergera microsphaerica* | Only preserved specimens | |
| *Schlumbergera opuntioides* | | |
| *Schlumbergera russelliana* | Only preserved specimens | |
| *Schlumbergera truncata* | | Major/ all excluded |
| *Sclerocactus brevihamatus* | | |
| *Sclerocactus erectocentrus* | | |
| *Sclerocactus intertextus* | | |
| *Sclerocactus johnsonii* | Only *i*Naturalist | |
| *Sclerocactus mariposensis* | | |
| *Sclerocactus papyracanthus* | Only *i*Naturalist | |
| *Sclerocactus scheeri* | | |

| | | |
|---|---|---|
| *Sclerocactus unguispinus* | | |
| *Sclerocactus warnockii* | | |
| *Selenicereus anthonyanus* | | Minor |
| *Selenicereus atropilosus* | Only preserved specimens | |
| *Selenicereus calcaratus* | Only preserved specimens | |
| *Selenicereus costaricensis* | Only preserved specimens | Minor |
| *Selenicereus escuintlensis* | Only preserved specimens | |
| *Selenicereus extensus* | | |
| *Selenicereus glaber* | Only preserved specimens | |
| *Selenicereus grandiflorus* | | |
| *Selenicereus guatemalensis* | Only preserved specimens | |
| *Selenicereus hamatus* | | |
| *Selenicereus inermis* | | |
| *Selenicereus megalanthus* | | |
| *Selenicereus minutiflorus* | Only preserved specimens | |
| *Selenicereus monacanthus* | Only preserved specimens | Minor |
| *Selenicereus nelsonii* | Only *i*Naturalist | |
| *Selenicereus ocamponis* | Only preserved specimens | |
| *Selenicereus pteranthus* | Only preserved specimens | Minor |
| *Selenicereus setaceus* | | |
| *Selenicereus spinulosus* | | Minor |
| *Selenicereus stenopterus* | Only preserved specimens | |
| *Selenicereus tonduzii* | Only preserved specimens | |
| *Selenicereus triangularis* | Only preserved specimens | Minor |
| *Selenicereus tricae* | | |
| *Selenicereus undatus* | | Major/ all excluded |
| *Selenicereus vagans* | | |
| *Selenicereus validus* | Only *i*Naturalist | |
| *Setiechinopsis mirabilis* | | |
| *Soehrensia angelesiae* | Only *i*Naturalist | |
| *Soehrensia arboricola* | Only preserved specimens | |
| *Soehrensia camarguensis* | | |
| *Soehrensia candicans* | | |
| *Soehrensia caulescens* | Only *i*Naturalist | |
| *Soehrensia formosa* | | |
| *Soehrensia grandiflora* | | |
| *Soehrensia hahniana* | Only preserved specimens | Minor/ only record in US excluded |
| *Soehrensia huascha* | | |
| *Soehrensia quadratiumbonata* | | |
| *Soehrensia schickendantzii* | Only *i*Naturalist | |

| | | |
|---|---|---|
| *Soehrensia serpentina* | | |
| *Soehrensia strigosa* | | |
| *Soehrensia tarijensis* | | |
| *Soehrensia thelegona* | | |
| *Soehrensia thelegonoides* | | |
| *Soehrensia vasquezii* | | |
| *Soehrensia volliana* | | |
| *Soehrensia walteri* | | |
| *Stenocactus coptonogonus* | | |
| *Stenocactus crispatus* | | |
| *Stenocactus dichroacanthus* | Only preserved specimens | |
| *Stenocactus multicostatus* | | |
| *Stenocactus obvallatus* | | |
| *Stenocactus ochoterenianus* | | |
| *Stenocactus phyllacanthus* | | |
| *Stenocactus sulphureus* | | |
| *Stenocactus vaupelianus* | | |
| *Stenocereus alamosensis* | | |
| *Stenocereus beneckei* | | |
| *Stenocereus chacalapensis* | | |
| *Stenocereus chrysocarpus* | | |
| *Stenocereus eruca* | | |
| *Stenocereus fricii* | | |
| *Stenocereus griseus* | | |
| *Stenocereus gummosus* | | |
| *Stenocereus heptagonus* | | |
| *Stenocereus humilis* | Only preserved specimens | |
| *Stenocereus kerberi* | | |
| *Stenocereus martinezii* | | |
| *Stenocereus montanus* | | |
| *Stenocereus pruinosus* | | |
| *Stenocereus queretaroensis* | | Minor |
| *Stenocereus quevedonis* | | |
| *Stenocereus standleyi* | | |
| *Stenocereus stellatus* | | |
| *Stenocereus thurberi* | | Minor |
| *Stenocereus treleasei* | | |
| *Stenocereus yunckeri* | Only preserved specimens | |
| *Stenocereus zopilotensis* | Only preserved specimens | |
| *Stephanocereus leucostele* | Only preserved specimens | |

| | | |
|---|---|---|
| *Stephanocereus luetzelburgii* | Only preserved specimens | |
| *Stetsonia coryne* | | |
| *Strombocactus corregidorae* | Only preserved specimens | |
| *Strombocactus disciformis* | | |
| *Strophocactus brasiliensis* | Only preserved specimens | |
| *Strophocactus wittii* | | |
| *Tacinga braunii* | Only preserved specimens | |
| *Tacinga funalis* | Only preserved specimens | Minor |
| *Tacinga inamoena* | | Minor |
| *Tacinga palmadora* | | Minor |
| *Tacinga saxatilis* | | |
| *Tacinga werneri* | Only preserved specimens | Minor |
| *Tephrocactus alexanderi* | | |
| *Tephrocactus aoracanthus* | | |
| *Tephrocactus articulatus* | | |
| *Tephrocactus bonnieae* | Only preserved specimens | |
| *Tephrocactus halophilus* | Only preserved specimens | |
| *Tephrocactus molinensis* | | Minor |
| *Tephrocactus nigrispinus* | | Minor |
| *Tephrocactus recurvatus* | | |
| *Tephrocactus verschaffeltii* | Only preserved specimens | |
| *Tephrocactus weberi* | | |
| *Thelocactus bicolor* | | |
| *Thelocactus buekii* | | |
| *Thelocactus conothelos* | | |
| *Thelocactus hastifer* | | |
| *Thelocactus hexaedrophorus* | | |
| *Thelocactus leucacanthus* | | |
| *Thelocactus macdowellii* | | |
| *Thelocactus rinconensis* | Only preserved specimens | |
| *Thelocactus setispinus* | | |
| *Thelocactus tulensis* | | |
| *Trichocereus bridgesii* | | Minor |
| *Trichocereus chalaensis* | Only HO excluding *i*Naturalist | |
| *Trichocereus clavatus* | Only preserved specimens | |
| *Trichocereus cuzcoensis* | | |
| *Trichocereus johnsonii* | Only preserved specimens | |
| *Trichocereus macrogonus* | | Minor |
| *Trichocereus tacaquirensis* | Only *i*Naturalist | |
| *Turbinicarpus alonsoi* | | |

| | | |
|---|---|---|
| *Turbinicarpus hoferi* | | |
| *Turbinicarpus lophophoroides* | | |
| *Turbinicarpus pseudopectinatus* | | |
| *Turbinicarpus saueri* | | |
| *Turbinicarpus schmiedickeanus* | | |
| *Turbinicarpus valdezianus* | | |
| *Turbinicarpus viereckii* | Only preserved specimens | |
| *Uebelmannia buiningii* | Only preserved specimens | |
| *Uebelmannia gummifera* | Only preserved specimens | |
| *Uebelmannia pectinifera* | | |
| *Vatricania guentheri* | | |
| *Weberbauerocereus cephalomacrostibas* | | |
| *Weberbauerocereus cuzcoensis* | Only *i*Naturalist | |
| *Weberbauerocereus madidiensis* | | |
| *Weberbauerocereus rauhii* | | |
| *Weberbauerocereus weberbaueri* | | |
| *Weberocereus bradei* | Only preserved specimens | |
| *Weberocereus frohningiorum* | Only preserved specimens | |
| *Weberocereus imitans* | Only preserved specimens | |
| *Weberocereus rosei* | Only preserved specimens | |
| *Weberocereus trichophorus* | Only preserved specimens | |
| *Weberocereus tunilla* | Only preserved specimens | Minor |
| *Weingartia cintia* | Only *i*Naturalist | |
| *Weingartia fidana* | | |
| *Weingartia neocumingii* | Only preserved specimens | |
| *Xiquexique frewenii* | Only preserved specimens | |
| *Xiquexique gounellei* | | |
| *Xiquexique tuberculatus* | | |
| *Yavia cryptocarpa* | Only preserved specimens | |
| *Yungasocereus inquisivensis* | | |

S3. Histogram of records per species in each basis of records: human observations excluding *i*Naturalist, preserved specimens, and *i*Naturalist only.

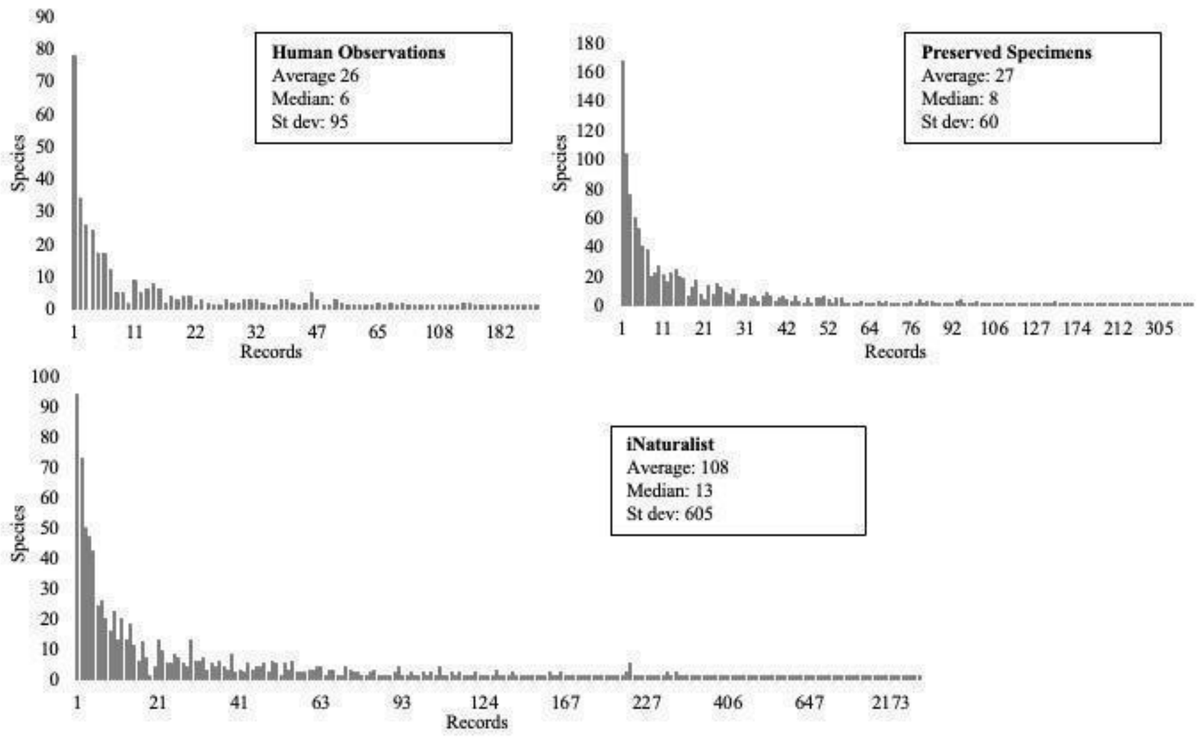

S4. Bioregionalization for neotropical cacti based on preserved specimens data only and a single solution method, resulting in 34 bioregions highlighted in colors (cluster cost= 0.92).

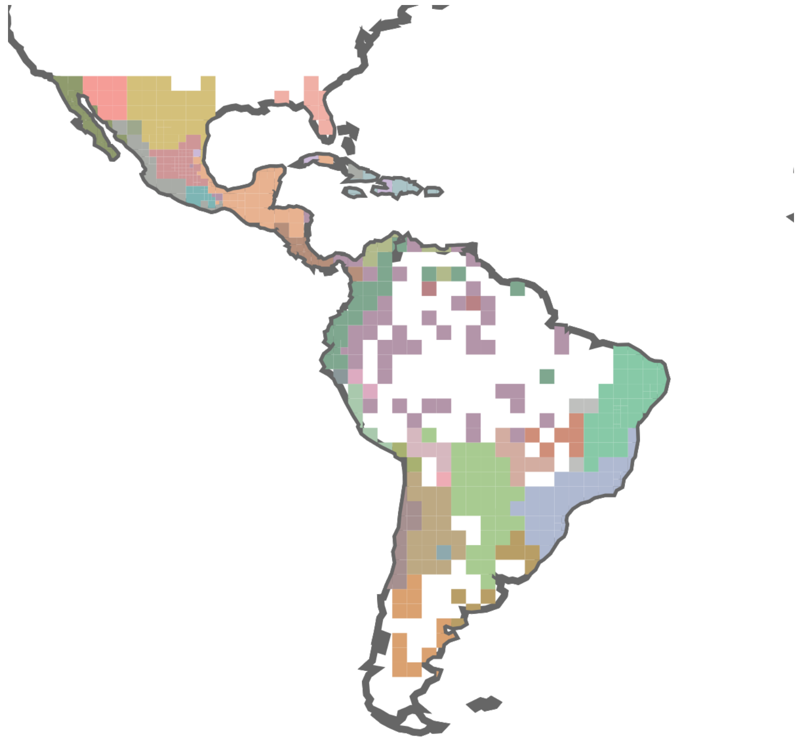

S5. Summary data for 24 bioregions of Neotropical Cactaceae: number of records, species, most common species, and most indicative species.

**BIOREGION 01** (2534 records of 134 species)

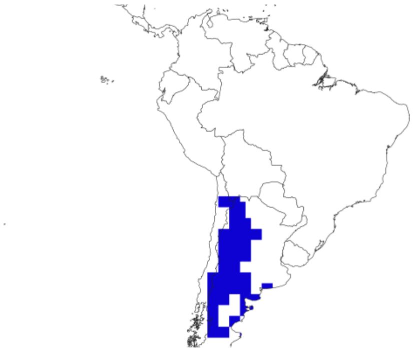

| Most common species (count) | Most indicative species (Score) |
|---|---|
| *Opuntia sulphurea* (190) | ***Soehrensia huascha**** (91.6) |
| ***Soehrensia candicans**** (148) | ***Rebutia minuscula**** (91.6) |
| ***Acanthocalycium leucanthum**** (141) | ***Echinopsis haematantha**** (91.6) |
| *Leucostele atacamensis* (136) | ***Echinopsis aurea**** (91.6) |
| ***Soehrensia formosa**** (125) | ***Maihueniopsis hickenii**** (91.6) |
| *Cereus aethiops* (105) | ***Tephrocactus molinensis**** (91.6) |
| ***Denmoza rhodacantha**** (103) | ***Soehrensia candicans**** (91.6) |
| *Leucostele terscheckii* (91) | ***Acanthocalycium leucanthum**** (91.6) |
| *Maihuenia patagonica* (65) | ***Soehrensia formosa**** (91.6) |
| *Maihueniopsis glomerata* (64) | ***Gymnocalycium monvillei**** (91.6) |

*Species restricted to the bioregion are marked in bold.

**BIOREGION 02** (2009 records of 59 species)

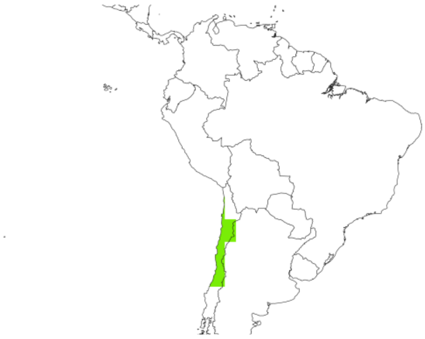

| Most common species (count) | Most indicative species (Score) |
|---|---|
| ***Leucostele chiloensis**** (359) | ***Copiapoa cinerascens**** (48.5) |
| ***Copiapoa cinerea**** (323) | ***Eriosyce napina**** (48.5) |
| ***Eriosyce paucicostata**** (180) | ***Eriosyce eriosyzoides**** (48.5) |

| | |
|---|---|
| ***Eriosyce curvispina**** (178) | ***Eriosyce heinrichiana**** (48.5) |
| ***Eulychnia breviflora**** (163) | ***Copiapoa dealbata**** (48.5) |
| ***Eriosyce subgibbosa**** (74) | ***Eriosyce chilensis**** (48.5) |
| *Cumulopuntia sphaerica* (72) | ***Copiapoa serpentisulcata**** (48.5) |
| ***Copiapoa coquimbana**** (60) | ***Copiapoa hypogaea**** (48.3) |
| ***Eulychnia iquiquensis**** (49) | ***Eriosyce crispa**** (48.5) |
| ***Copiapoa humilis**** (48) | ***Eriosyce senilis**** (48.5) |

*Species restricted to the bioregion are marked in bold.

**BIOREGION 03** (1639 records of 132 species)

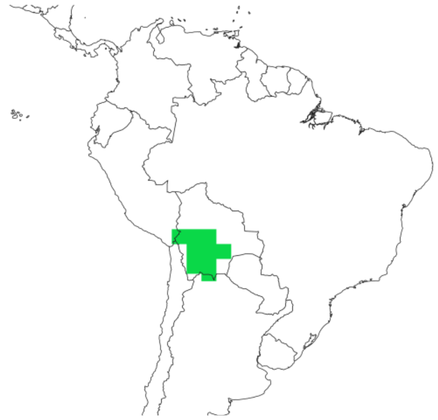

| Most common species (count) | Most indicative species (Score) |
|---|---|
| *Corryocactus melanotrichus* (126) | ***Lobivia calorubra**** (138.1) |
| *Cumulopuntia boliviana* (69) | ***Lobivia obrepanda**** (138.1) |
| ***Lobivia bridgesii**** (64) | ***Lobivia chrysochete**** (138.1) |
| *Opuntia sulphurea* (61) | ***Harrisia tetracantha**** (138.1) |
| *Leucostele atacamensis* (56) | ***Cleistocactus parviflorus**** (138.1) |
| *Austrocylindropuntia shaferi* (49) | ***Cleistocactus candelilla**** (138.1) |
| *Rhipsalis floccosa* (48) | ***Pereskia diaz-romeroana**** (138.1) |
| *Lobivia calorubra* (38) | ***Lobivia pamparuizii**** (138.1) |
| *Lobivia maximiliana* (37) | ***Lobivia lateritia**** (138.1) |
| ***Lobivia obrepanda**** (36) | ***Vatricania guentheri**** (138.1) |

*Species restricted to the bioregion are marked in bold.

**BIOREGION 04** (1414 records of 82 species)

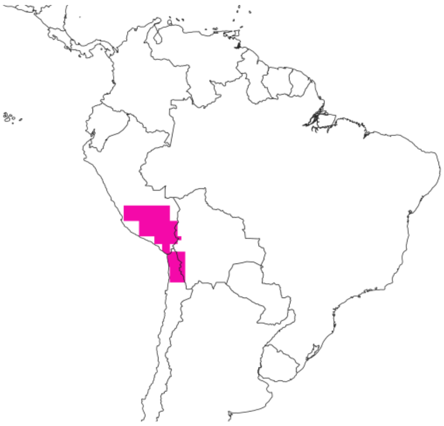

| Most common species (count) | Most indicative species (Score) |
|---|---|
| *Austrocylindropuntia subulata* (85) | ***Corryocactus brevistylus**** (204.7) |
| ***Corryocactus brevistylus**** (76) | ***Corryocactus aureus**** (204.7) |
| *Cumulopuntia leucophaea* (52) | ***Corryocactus erectus**** (204.7) |
| *Corryocactus erectus* (47) | ***Weberbauerocereus weberbaueri**** (204.7) |
| *Lobivia maximiliana* (46) | ***Lobivia pampana**** (204.7) |
| *Austrocylindropuntia floccosa* (43) | ***Punotia lagopus**** (204.7) |
| *Cumulopuntia boliviana* (41) | ***Lobivia hertrichiana**** (204.7) |
| *Browningia candelaris* (37) | ***Armatocereus riomajensis**** (204.7) |
| *Trichocereus cuzcoensis* (37) | ***Haageocereus platinospinus**** (204.7) |
| *Cumulopuntia sphaerica* (36) | ***Browningia hertlingiana**** (204.7) |

*Species restricted to the bioregion are marked in bold.

**BIOREGION 05** (708 records of 47 species)

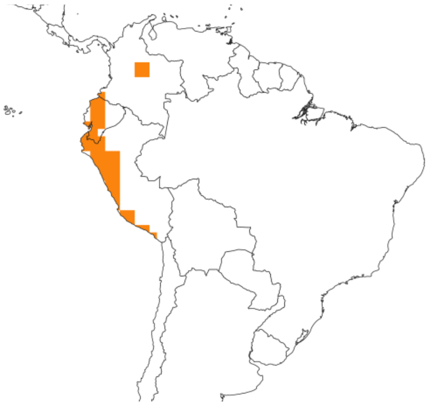

| Most common species (count) | Most indicative species (Score) |
|---|---|
| *Austrocylindropuntia cylindrica* (191) | ***Browningia microsperma**** (91.1) |
| *Borzicactus sepium* (152) | ***Melocactus andinus**** (91.1) |

| | |
|---|---|
| *Opuntia soederstromiana* (122) | ***Borzicactus leonensis**** (91.1) |
| *Rhipsalis micrantha* (55) | ***Lymanbensonia brevispina**** (91.1) |
| ***Opuntia quitensis**** (51) | ***Browningia altissima**** (91.1) |
| *Austrocylindropuntia subulata* (47) | ***Austrocylindropuntia pachypus**** (91.1) |
| *Trichocereus macrogonus* (40) | ***Armatocereus procerus**** (91.1) |
| ***Espostoa lanata**** (38) | ***Haageocereus versicolor**** (91.1) |
| *Austrocylindropuntia floccosa* (36) | ***Weberocereus rosei**** (91.1) |
| *Armatocereus matucanensis* (34) | ***Matucana weberbaueri**** (91.1) |

*Species restricted to the bioregion are marked in bold.

**BIOREGION 06** (11 records of 10 species)

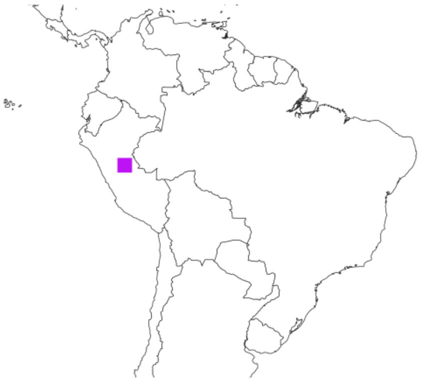

| Most common species (count) | Most indicative species (Score) |
|---|---|
| *Rhipsalis baccifera* (2) | *Trichocereus johnsonii* (8701.5) |
| *Armatocereus laetus* (1) | *Matucana madisoniorum* (8701.5) |
| *Borzicactus acanthurus* (1) | *Calymmanthium substerile* (8701.5) |
| *Borzicactus icosagonus* (1) | *Browningia utcubambensis* (8701.5) |
| *Borzicactus neoroezlii* (1) | *Armatocereus laetus* (8701.5) |
| *Borzicactus tenuiserpens* (1) | *Borzicactus tenuiserpens* (4350.8) |
| *Browningia utcubambensis* (1) | *Borzicactus acanthurus* (669.3) |
| *Calymmanthium substerile* (1) | *Borzicactus neoroezlii* (580.1) |
| *Matucana madisoniorum* (1) | *Borzicactus icosagonus* (280.7) |
| *Trichocereus johnsonii* (1) | *Rhipsalis baccifera* (14.6) |

**BIOREGION 07** (1404 records of 84 species)

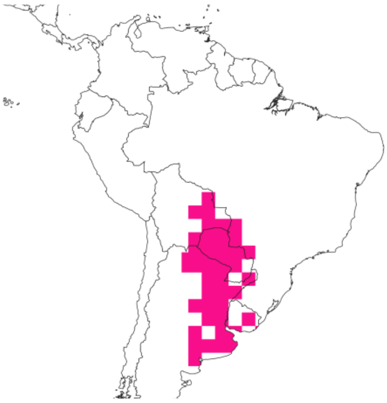

| Most common species (count) | Most indicative species (Score) |
|---|---|
| *Opuntia elata* (115) | ***Acanthocalycium rhodotrichum**** (151.3) |
| *Lepismium lumbricoides* (85) | ***Harrisia bonplandii**** (151.3) |
| *Cleistocactus baumannii* (77) | ***Echinopsis calochlora**** (151.3) |
| *Rhipsalis cereuscula* (48) | ***Cereus phatnospermus**** (151.3) |
| *Harrisia pomanensis* (48) | ***Cereus spegazzinii**** (151.3) |
| *Pereskia aculeata* (48) | ***Harrisia regelii**** (151.3) |
| *Pereskia sacharosa* (46) | ***Opuntia rioplatense**** (151.3) |
| *Lepismium cruciforme* (45) | ***Parodia schumanniana**** (151.3) |
| *Cereus stenogonus* (44) | ***Gymnocalycium friedrichii**** (151.3) |
| *Rhipsalis baccifera* (43) | ***Parodia nigrispina**** (151.3) |

*Species restricted to the bioregion are marked in bold.

**BIOREGION 08** (73 records of 21 species)

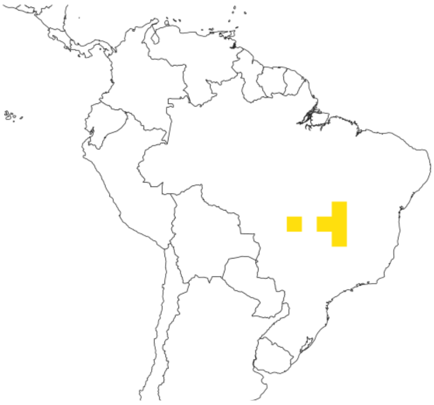

| Most common species (count) | Most indicative species (Score) |
|---|---|
| ***Pilosocereus vilaboensis**** (12) | ***Micranthocereus estevesii**** (1450.3) |
| *Epiphyllum phyllanthus* (9) | ***Cereus pierre-braunianus*** (1450.3) |

| | |
|---|---|
| *Discocactus heptacanthus* (8) | ***Discocactus diersianus**** (1450.3) |
| ***Micranthocereus estevesii**** (7) | ***Pilosocereus albisummus**** (1450.3) |
| *Pilosocereus machrisii* (7) | ***Pilosocereus diersianus**** (1450.3) |
| *Cereus jamacaru* (5) | ***Pilosocereus hermii**** (1450.3) |
| *Rhipsalis baccifera* (5) | ***Pilosocereus vilaboensis**** (1338.7) |
| ***Cereus pierre-braunianus**** (5) | *Pilosocereus machrisii* (922.9) |
| ***Discocactus diersianus**** (2) | *Discocactus heptacanthus* (725.1) |
| ***Pilosocereus albisummus**** (2) | *Discocactus catingicola* (161.1) |

*Species restricted to the bioregion are marked in bold.

**BIOREGION 09** (2757 records of 91 species)

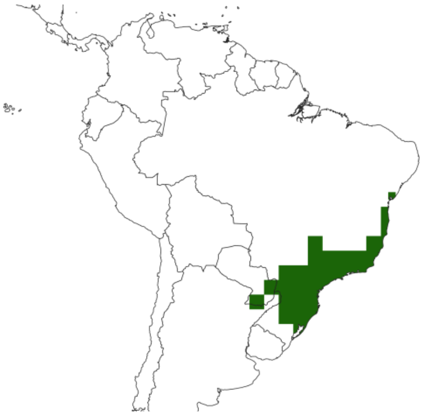

| Most common species (count) | Most indicative species (Score) |
|---|---|
| *Pereskia aculeata* (222) | ***Rhipsalis pulchra**** (78.4) |
| *Rhipsalis teres* (213) | ***Rhipsalis puniceodiscus**** (78.4) |
| *Hatiora salicornioides* (205) | ***Rhipsalidopsis gaertneri**** (78.4) |
| *Rhipsalis floccosa* (173) | ***Rhipsalis dissimilis**** (78.4) |
| *Lepismium houlletianum* (154) | ***Rhipsalis trigona**** (78.4) |
| *Lepismium cruciforme* (151) | ***Rhipsalis hoelleri**** (78.4) |
| *Brasiliopuntia brasiliensis* (130) | ***Rhipsalis flagelliformis**** (78.4) |
| *Rhipsalis elliptica* (120) | ***Rhipsalis ormindoi**** (78.4) |
| ***Rhipsalis pachyptera**** (96) | ***Parodia carambeiensis**** (78.4) |
| *Cereus fernambucensis* (94) | ***Cipocereus laniflorus**** (78.4) |

*Species restricted to the bioregion are marked in bold.

**BIOREGION 10** (1731 records of 59 species)

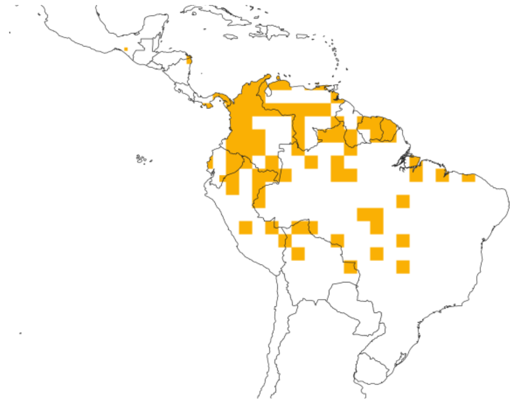

| Most common species (count) | Most indicative species (Score) |
|---|---|
| ***Leuenbergeria guamacho**** (291) | ***Melocactus schatzlii**** (59.8) |
| *Rhipsalis baccifera* (219) | ***Stenocereus humilis**** (59.8) |
| *Epiphyllum phyllanthus* (218) | ***Melocactus smithii**** (59.8) |
| *Pseudorhipsalis amazonica* (206) | ***Opuntia schumannii**** (59.8) |
| *Acanthocereus tetragonus* (102) | ***Selenicereus extensus**** (59.8) |
| *Melocactus curvispinus* (85) | ***Melocactus estevesii**** (59.8) |
| *Leuenbergeria bleo* (59) | ***Opuntia curassavica**** (59.8) |
| *Stenocereus griseus* (55) | ***Melocactus mazelianus**** (59.8) |
| *Armatocereus cartwrightianus* (38) | ***Mammillaria mammillaris**** (59.8) |
| *Opuntia pittieri* (33) | ***Browningia hernandezii**** (59.8) |

*Species restricted to the bioregion are marked in bold.

**BIOREGION 11** (411 records of 44 species)

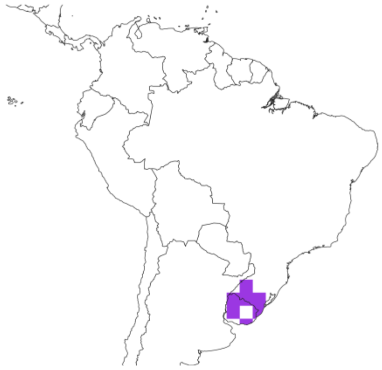

| Most common species (count) | Most indicative species (Score) |
|---|---|
| *Parodia ottonis* (118) | ***Parodia scopa**** (147.5) |
| *Parodia erinacea* (50) | ***Parodia crassigibba**** (147.5) |
| ***Parodia scopa**** (28) | ***Parodia tenuicylindrica**** (147.5) |

| | |
|---|---|
| *Frailea gracillima* (24) | ***Parodia mueller-melchersii**** (147.5) |
| *Frailea pygmaea* (23) | ***Frailea castanea**** (147.5) |
| *Parodia oxycostata* (22) | ***Parodia muricata**** (147.5) |
| *Lepismium lumbricoides* (17) | ***Parodia gaucha**** (147.5) |
| ***Parodia concinna**** (17) | ***Parodia fusca**** (147.5) |
| *Parodia mammulosa* (15) | ***Parodia curvispina**** (147.5) |
| *Opuntia elata* (11) | ***Frailea phaeodisca**** (147.5) |

*Species restricted to the bioregion are marked in bold.

**BIOREGION 12** (55 records of 13 species)

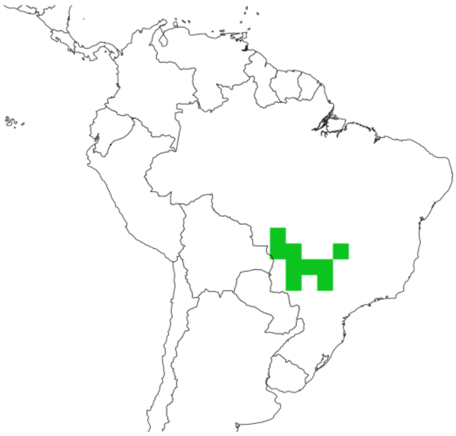

| **Most common species (count)** | **Most indicative species (Score)** |
|---|---|
| *Cereus bicolor* (18) | ***Cereus saddianus**** (966.8) |
| *Epiphyllum phyllanthus* (7) | ***Pilosocereus jauruensis**** (966.8) |
| *Discocactus heptacanthus* (7) | *Cereus bicolor* (828.7) |
| ***Pilosocereus jauruensis**** (6) | *Discocactus heptacanthus* (423) |
| *Brasiliopuntia brasiliensis* (5) | *Discocactus ferricola* (193.4) |
| ***Cereus saddianus**** (4) | *Cleistocactus baumannii* (17.4) |
| *Cleistocactus baumannii* (2) | *Brasiliopuntia brasiliensis* (16.2) |
| *Rhipsalis baccifera* (1) | *Pereskia sacharosa* (14) |
| *Hatiora salicornioides* (1) | *Epiphyllum phyllanthus* (10.8) |
| *Pereskia sacharosa* (1) | *Praecereus euchlorus* (8.6) |

*Species restricted to the bioregion are marked in bold.

**BIOREGION 13** (3385 records of 121 species)

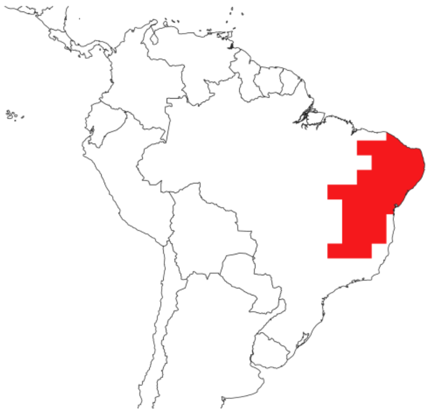

| Most common species (count) | Most indicative species (Score) |
|---|---|
| *Tacinga inamoena* (280) | ***Melocactus zehntneri**** (62.2) |
| *Cereus jamacaru* (210) | ***Pilosocereus pentaedrophorus**** (62.2) |
| *Xiquexique gounellei* (153) | ***Micranthocereus purpureus**** (62.2) |
| ***Tacinga palmadora**** (149) | ***Stephanocereus luetzelburgii**** (62.2) |
| *Pilosocereus pachycladus* (147) | ***Leocereus bahiensis**** (62.2) |
| *Pilosocereus catingicola* (127) | ***Pilosocereus chrysostele**** (62.2) |
| *Arrojadoa rhodantha* (116) | ***Micranthocereus flaviflorus**** (62.2) |
| ***Melocactus zehntneri**** (99) | ***Melocactus glaucescens**** (62.2) |
| *Arrojadoa penicillata* (93) | ***Micranthocereus polyanthus**** (62.2) |
| ***Pereskia bahiensis**** (82) | ***Melocactus paucispinus**** (62.2) |

*Species restricted to the bioregion are marked in bold.

**BIOREGION 14** (4159 records of 156 species)

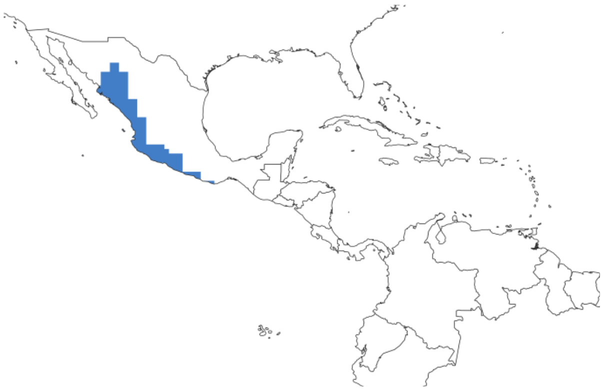

| Most common species (count) | Most indicative species (Score) |
|---|---|
| *Pachycereus pecten-aboriginum* (731) | ***Stenocereus martinezii**** (23.8) |
| *Opuntia decumbens* (237) | ***Pereskiopsis blakeana**** (23.8) |

| | |
|---|---|
| **_Stenocereus martinezii*_** (228) | **_Stenocereus chrysocarpus_** (23.8) |
| *Acanthocereus tetragonus* (228) | **_Stenocereus fricii*_** (23.8) |
| *Stenocereus alamosensis* (203) | **_Mammillaria marksiana*_** (23.8) |
| *Opuntia puberula* (113) | **_Pachycereus tepamo*_** (23.8) |
| *Pilosocereus purpusii* (97) | **_Echinocereus subinermis*_** (23.8) |
| **_Stenocereus kerberi*_** (94) | **_Selenicereus atropilosus*_** (23.8) |
| *Opuntia tomentosa* (90) | **_Acanthocereus cuixmalensis*_** (23.8) |
| *Opuntia excelsa* (87) | **_Acanthocereus tepalcatepecanus*_** (23.8) |

*Species restricted to the bioregion are marked in bold.

**BIOREGION 15** (12886 records of 102 species)

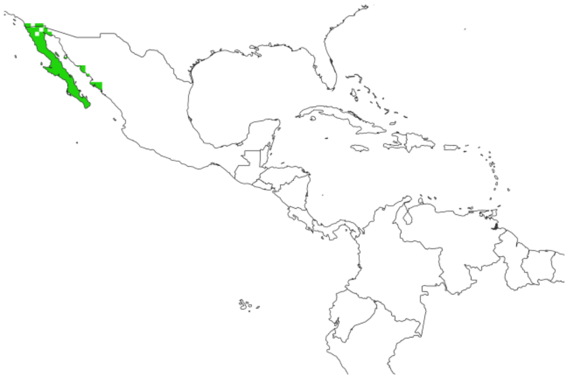

| Most common species (count) | Most indicative species (Score) |
|---|---|
| *Cochemiea dioica* (1127) | **_Echinocereus maritimus*_** (15.4) |
| *Pachycereus pringlei* (1087) | **_Ferocactus gracilis*_** (15.4) |
| *Stenocereus gummosus* (701) | **_Cylindropuntia tesajo*_** (15.4) |
| *Echinocereus engelmannii* (651) | **_Ferocactus fordii*_** (15.4) |
| *Ferocactus viridescens* (628) | **_Cochemiea schumannii*_** (15.4) |
| *Lophocereus schottii* (584) | **_Opuntia pycnantha*_** (15.4) |
| *Ferocactus cylindraceus* (550) | **_Cochemiea blossfeldiana*_** (15.4) |
| **_Echinocereus maritimus*_** (522) | **_Ferocactus diguetii*_** (15.4) |
| *Cylindropuntia cholla* (516) | **_Ferocactus chrysacanthus*_** (15.4) |
| *Bergerocactus emoryi* (513) | **_Opuntia bravoana*_** (15.4) |

*Species restricted to the bioregion are marked in bold.

**BIOREGION 16** (37659 records of 132 species)

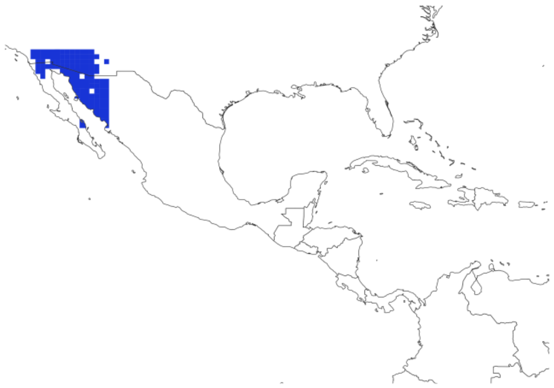

| Most common species (count) | Most indicative species (Score) |
|---|---|
| *Carnegiea gigantea* (17375) | ***Sclerocactus erectocentrus**** (1.0) |
| *Ferocactus wislizenii* (3240) | ***Echinocereus ledingii**** (1.0) |
| *Cylindropuntia fulgida* (2396) | ***Cochemiea thornberi**** (1.0) |
| *Echinocereus engelmannii* (1832) | ***Echinocereus klapperi**** (1.0) |
| *Cochemiea grahamii* (1539) | ***Echinocereus pseudopectinatus**** (1.0) |
| *Cylindropuntia acanthocarpa* (1332) | ***Grusonia emoryi**** (1.0) |
| *Ferocactus cylindraceus* (1140) | ***Cochemiea boolii**** (1.0) |
| *Cylindropuntia imbricata* (1036) | ***Echinocereus bristolii**** (1.0) |
| *Opuntia engelmannii* (1025) | ***Grusonia reflexispina**** (1.0) |
| *Cylindropuntia thurberi* (966) | ***Sclerocactus johnsonii**** (1.0) |

*Species restricted to the bioregion are marked in bold.

**BIOREGION 17** (13472 records of 223 species)

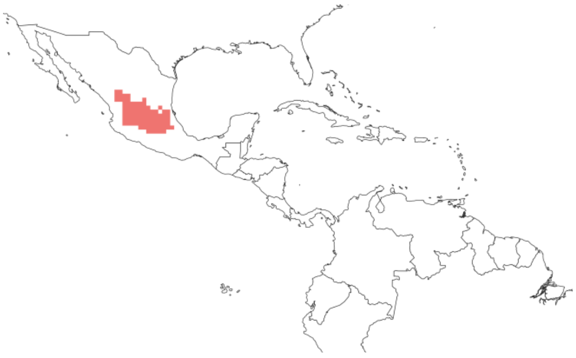

| Most common species (count) | Most indicative species (Score) |
|---|---|
| *Myrtillocactus geometrizans* (735) | ***Stenocactus phyllacanthus**** (23.7) |
| *Ferocactus histrix* (659) | ***Coryphantha vogtherriana**** (23.7) |

| | |
|---|---|
| *Opuntia tomentosa* (612) | ***Echinocereus pamanesii**** (23.7) |
| *Cylindropuntia imbricata* (587) | ***Stenocactus ochoterenianus**** (23.7) |
| *Opuntia robusta* (517) | ***Mammillaria wiesingeri**** (23.7) |
| *Mammillaria magnimamma* (435) | ***Coryphantha clavata**** (23.7) |
| *Opuntia microdasys* (381) | ***Thelocactus leucacanthus**** (23.7) |
| *Opuntia streptacantha* (371) | ***Mammillaria albiflora**** (23.7) |
| *Opuntia engelmannii* (371) | ***Turbinicarpus alonsoi**** (23.7) |
| *Opuntia leucotricha* (319) | ***Coryphantha glassii**** (23.7) |

*Species restricted to the bioregion are marked in bold.

**BIOREGION 18** (9077 records of 165 species)

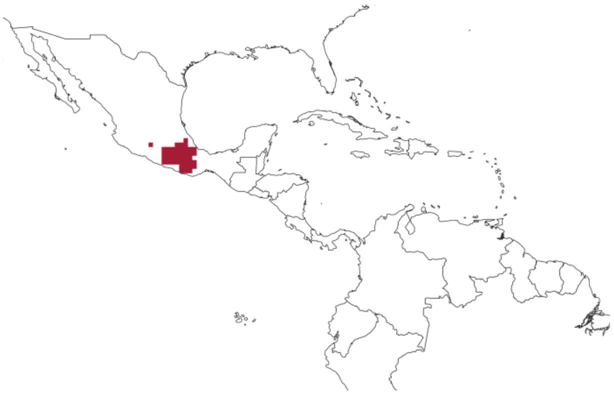

| Most common species (count) | Most indicative species (Score) |
|---|---|
| *Mammillaria haageana* (494) | ***Mammillaria pectinifera**** (35.2) |
| *Echinocactus platyacanthus* (412) | ***Stenocereus beneckei**** (35.2) |
| *Opuntia pilifera* (364) | ***Opuntia tehuacana**** (35.2) |
| ***Coryphantha pallida**** (343) | ***Stenocereus treleasei**** (35.2) |
| ***Mammillaria sphacelata**** (282) | ***Mammillaria flavicentra**** (35.2) |
| *Mammillaria carnea* (274) | ***Opuntia parviclada**** (35.2) |
| *Myrtillocactus geometrizans* (265) | ***Mammillaria knippeliana**** (35.2) |
| *Pachycereus weberi* (217) | ***Mammillaria varieaculeata**** (35.2) |
| *Cephalocereus tetetzo* (208) | ***Mammillaria magnifica**** (35.2) |
| *Opuntia depressa* (202) | ***Mammillaria tonalensis**** (35.2) |

*Species restricted to the bioregion are marked in bold.

**BIOREGION 19** (34003 records of 283 species)

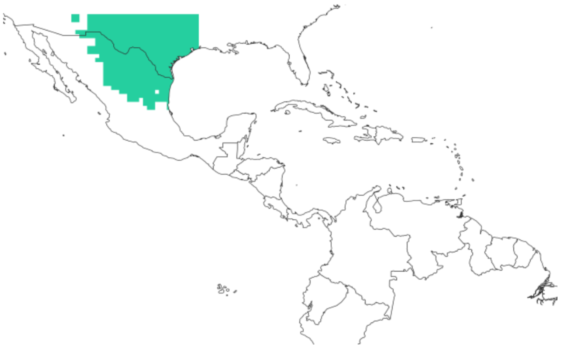

| Most common species (count) | Most indicative species (Score) |
|---|---|
| *Cylindropuntia leptocaulis* (3251) | ***Echinocereus longisetus**** (5.4) |
| *Echinocereus reichenbachii* (2180) | ***Echinocereus viridiflorus**** (5.4) |
| *Opuntia engelmannii* (1912) | ***Echinocereus viereckii**** (5.4) |
| *Echinocactus horizonthalonius* (1509) | ***Echinocereus russanthus**** (5.4) |
| *Cylindropuntia imbricata* (1405) | ***Coryphantha durangensis**** (5.4) |
| *Echinocereus coccineus* (1225) | ***Turbinicarpus valdezianus**** (5.4) |
| *Echinocereus enneacanthus* (1172) | ***Echinocereus parkeri**** (5.4) |
| *Mammillaria heyderi* (936) | ***Mammillaria sphaerica**** (5.4) |
| *Echinocereus dasyacanthus* (926) | ***Sclerocactus brevihamatus**** (5.4) |
| *Echinocactus texensis* (830) | ***Escobaria emskoetteriana**** (5.4) |

*Species restricted to the bioregion are marked in bold.

**BIOREGION 20** (335 records of 11 species)

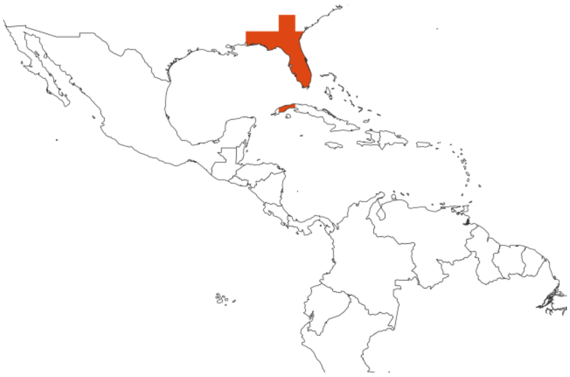

| Most common species (count) | Most indicative species (Score) |
|---|---|
| *Acanthocereus tetragonus* (169) | ***Leptocereus leonii**** (103.0) |
| *Opuntia humifusa* (73) | ***Opuntia drummondii* *** (101.5) |
| ***Opuntia drummondii* *** (70) | *Opuntia humifusa* (91.7) |

| | |
|---|---|
| *Opuntia stricta* (11) | *Opuntia triacantha* (51.5) |
| *Opuntia triacantha* (4) | *Acanthocereus tetragonus* (14.3) |
| *Mammillaria prolifera* (3) | *Melocactus harlowii* (4.7) |
| *Opuntia littoralis* (1) | *Opuntia stricta* (9.9) |
| *Rhipsalis baccifera* (1) | *Mammillaria prolifera* (1.3) |
| *Melocactus harlowii* (1) | *Opuntia littoralis* (0.9) |
| *Selenicereus gradiflorus* (1) | *Selenicereus gradiflorus* (0.6) |

*Species restricted to the bioregion are marked in bold.

**BIOREGION 21** (485 records of 39 species)

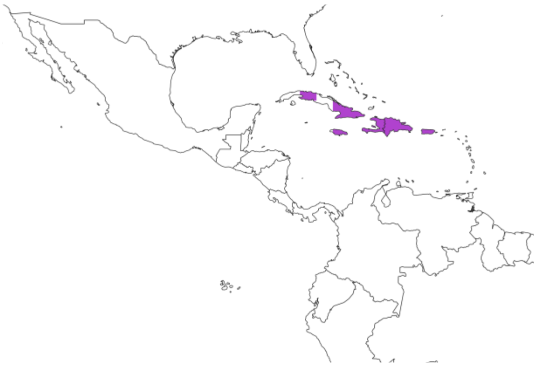

| Most common species (count) | Most indicative species (Score) |
|---|---|
| ***Melocactus intortus**** (61) | ***Melocactus lemairei**** (285.3) |
| ***Consolea moniliformis**** (54) | ***Leptocereus quadricostatus**** (285.3) |
| *Pilosocereus royenii* (45) | ***Leptocereus paniculatus**** (285.3) |
| *Rhipsalis baccifera* (37) | ***Leuenbergeria portulacifolia**** (285.3) |
| ***Stenocereus heptagonus**** (35) | ***Leptocereus weingartianus**** (285.3) |
| *Mammillaria prolifera* (23) | ***Consolea rubescens**** (285.3) |
| ***Harrisia gracilis**** (23) | ***Leptocereus sylvestris**** (285.3) |
| *Melocactus harlowii* (21) | ***Melocactus matanzanus**** (285.3) |
| ***Cylindropuntia caribaea**** (20) | ***Leuenbergeria marcanoi**** (285.3) |
| ***Melocactus lemairei**** (19) | ***Cylindropuntia hystrix**** (285.3) |

*Species restricted to the bioregion are marked in bold.

**BIOREGION 22** (5312 records of 161 species)

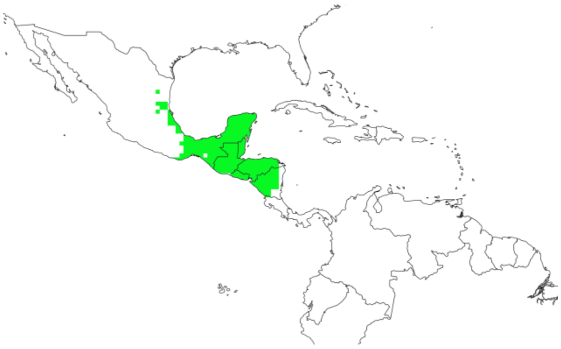

| Most common species (count) | Most indicative species (Score) |
| --- | --- |
| *Rhipsalis baccifera* (470) | ***Selenicereus pteranthus**** (37.0) |
| *Acanthocereus tetragonus* (369) | ***Epiphyllum chrysocardium**** (37.0) |
| *Opuntia decumbens* (343) | ***Selenicereus nelsonii**** (37.0) |
| *Pachycereus pecten-aboriginum* (330) | ***Deamia chontalensis**** (37.0) |
| ***Stenocereus chacalapensis**** (295) | ***Lemaireocereus lepidanthus**** (37.0) |
| *Leuenbergeria lychnidiflora* (291) | ***Selenicereus minutiflorus**** (37.0) |
| *Acanthocereus oaxacensis* (240) | ***Selenicereus escuintlensis**** (37.0) |
| *Pilosocereus leucocephalus* (219) | ***Cochemiea macdougallii**** (37.0) |
| *Selenicereus grandiflorus* (163) | ***Opuntia deamii**** (37.0) |
| *Opuntia stricta* (162) | ***Acanthocereus macdougallii**** (37.0) |

*Species restricted to the bioregion are marked in bold.

**BIOREGION 23** (863 records of 42 species)

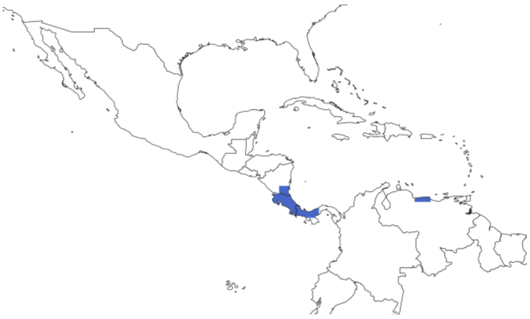

| Most common species (count) | Most indicative species (Score) |
| --- | --- |
| *Rhipsalis baccifera* (98) | ***Weberocereus imitans**** (177.6) |
| *Epiphyllum hookeri* (96) | ***Disocactus lepidocarpus**** (177.6) |
| ***Epiphyllum cartagense**** (58) | ***Pseudorhipsalis himantoclada**** (177.6) |

| | |
|---|---|
| *Selenicereus costaricensis* (53) | ***Weberocereus bradei**** (177.6) |
| ***Pseudorhipsalis himantoclada**** (51) | ***Pseudorhipsalis acuminata**** (177.6) |
| *Weberocereus tunilla* (47) | ***Selenicereus stenopterus**** (177.6) |
| *Acanthocereus tetragonus* (47) | ***Selenicereus tonduzii**** (177.6) |
| *Epiphyllum phyllanthus* (39) | ***Weberocereus trichophorus**** (177.6) |
| *Opuntia guatemalensis* (32) | ***Selenicereus calcaratus**** (177.6) |
| ***Weberocereus bradei**** (31) | ***Epiphyllum cartagense**** (177.6) |

*Species restricted to the bioregion are marked in bold.

**BIOREGION 24** (630 records of 3 species)

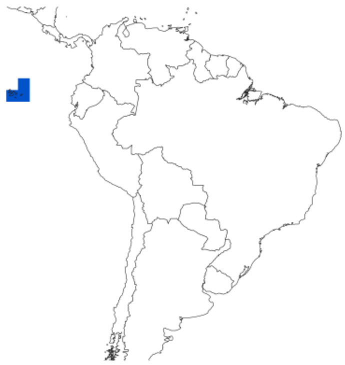

| Most common species (count) | Most indicative species (Score) |
|---|---|
| ***Opuntia galapageia**** (344) | ***Brachycereus nesioticus**** (50.6) |
| ***Jasminocereus thouarsii**** (170) | ***Opuntia galapageia**** (50.3) |
| ***Brachycereus nesioticus**** (116) | ***Jasminocereus thouarsii**** (50.0) |

*Species restricted to the bioregion are marked in bold.

S6. Hierarchical solution based on preserved specimens and human observations using Markov time = 1.18 showing in the second level a similar scheme to the non-hierarchical solution presented (Fig. 2). First level shows three bioregions, second level shows 22 bioregions, and third level shows 25 bioregions.

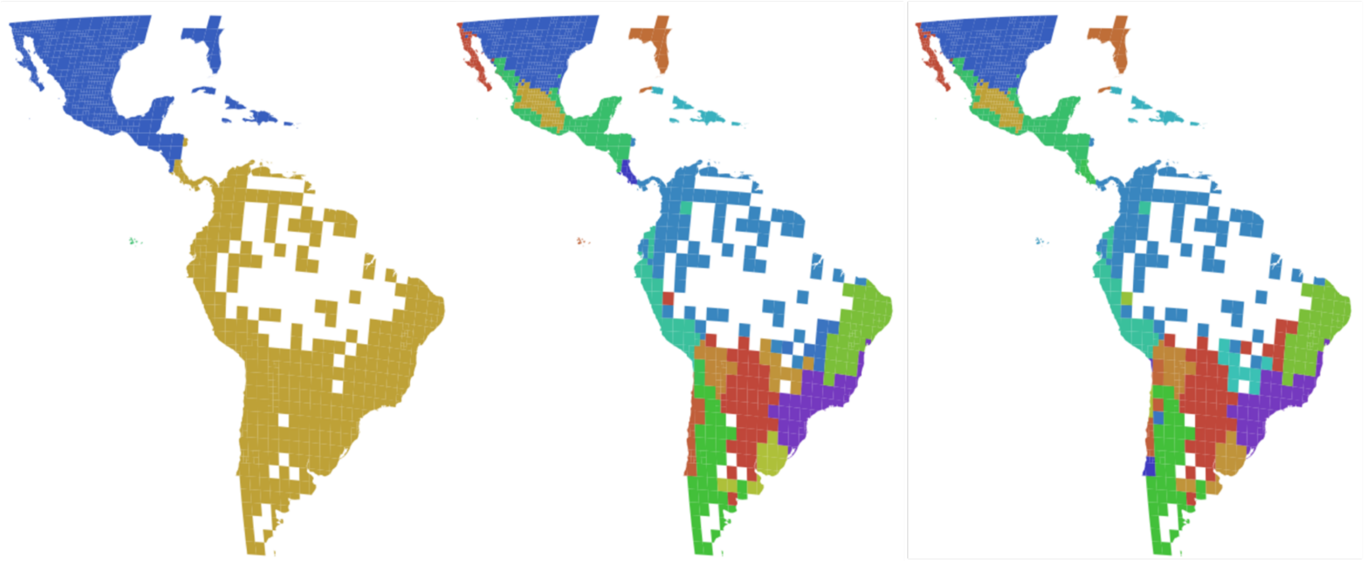

S7. Alternative hierarchical solution for North America (using Markov time = 1) based on preserved specimens and human observations. First level shows two bioregions for this area, second level shows 11 bioregions, and third level shows 14 bioregions.

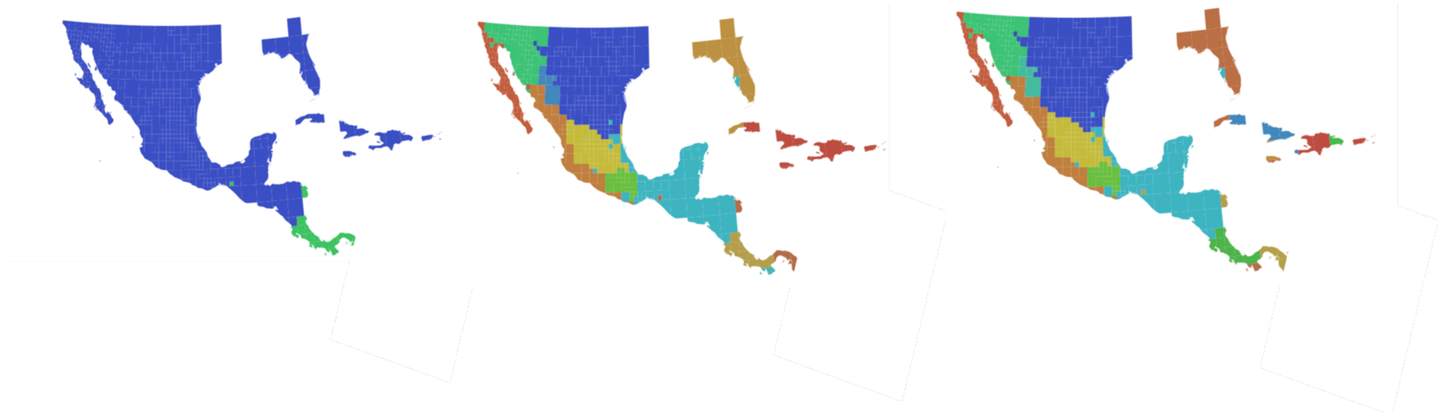

S8. Alternative first level of hierarchical solutions based on preserved specimens and human observations using Markov time = 1 (left) and Markov time = 0.96 (right).

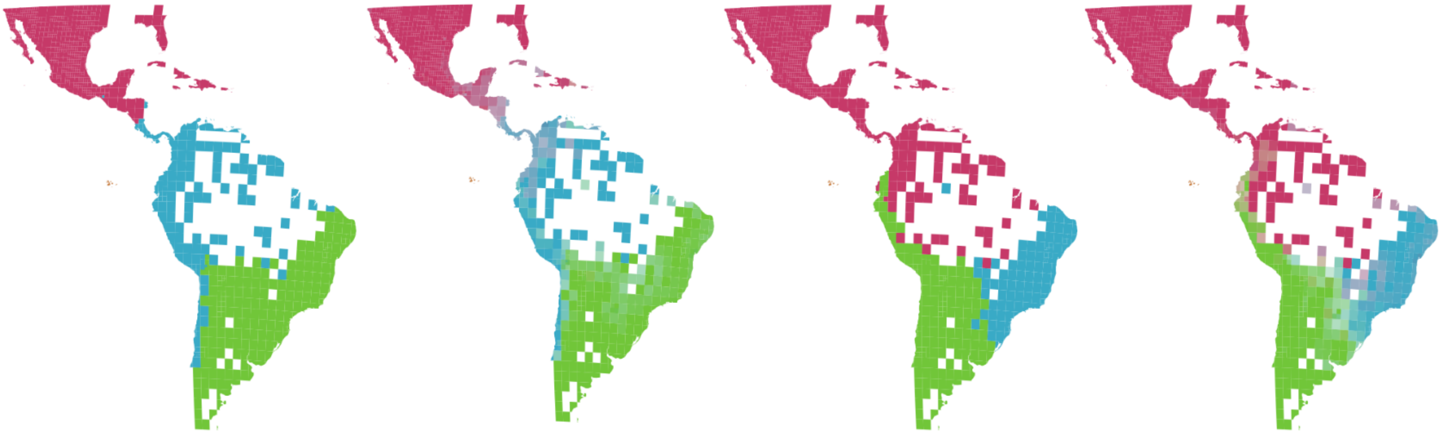

S9. Tree pruned from the megatree (Jin and Qian, 2022) including 616 species that matched our occurrence dataset. Major clades and genera in Cactaceae are collapsed.

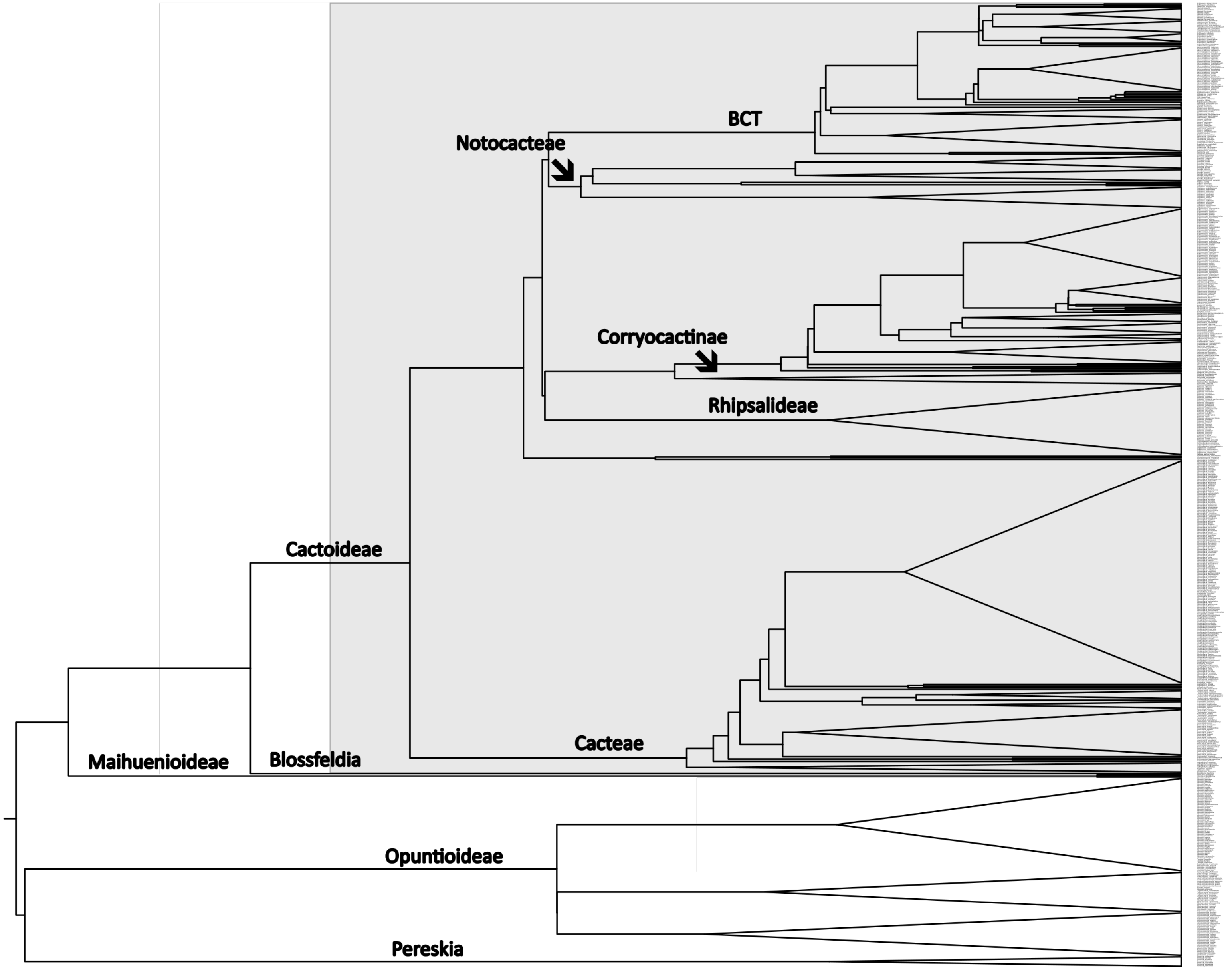
