## Supplementary material for "Spiny but photogenic: amateur sightings complement herbarium specimens to reveal the bioregions of cacti": non-hierarchical scheme shapefile: bioregions_nonhierarchical.qmd

```
<!DOCTYPE qgis PUBLIC 'http://mrcc.com/qgis.dtd' 'SYSTEM'>
<qgis version="3.24.1-Tisler">
  <identifier></identifier>
  <parentidentifier></parentidentifier>
  <language></language>
  <type>dataset</type>
  <title></title>
  <abstract></abstract>
  <links/>
  <fees></fees>
  <encoding></encoding>
  <crs>
    <spatialrefsys nativeFormat="Wkt">
      <wkt></wkt>
      <proj4></proj4>
      <srsid>0</srsid>
      <srid>0</srid>
      <authid></authid>
      <description></description>
      <projectionacronym></projectionacronym>
      <ellipsoidacronym></ellipsoidacronym>
      <geographicflag>false</geographicflag>
    </spatialrefsys>
  </crs>
  <extent/>
</qgis>
```
